# Epidermal maintenance of Langerhans cells relies on autophagy-regulated lipid metabolism

**DOI:** 10.1101/2022.09.16.507799

**Authors:** Florent Arbogast, Raquel Sal-Carro, Wacym Boufenghour, Quentin Frenger, Delphine Bouis, Louise Filippi De La Palavesa, Jean-Daniel Fauny, Olivier Griso, Hélène Puccio, Rebecca Fima, Thierry Huby, Emmanuel L. Gautier, Anne Molitor, Raphaël Carapito, Seiamak Bahram, Nikolaus Romani, Björn E. Clausen, Benjamin Voisin, Christopher G. Mueller, Frédéric Gros, Vincent Flacher

**Affiliations:** Laboratory CNRS I²CT/UPR3572 Immunology, Immunopathology and Therapeutic Chemistry, Strasbourg Drug Discovery and Development Institute (IMS), Institut de Biologie Moléculaire et Cellulaire, Strasbourg, France; Université de Strasbourg, Strasbourg, France; Laboratoire d’Immunorhumatologie Moléculaire, Plateforme GENOMAX, INSERM UMR_S 1109, Faculté de Médecine, Fédération Hospitalo-Universitaire OMICARE, ITI TRANSPLANTEX NG, Université de Strasbourg, 67085 Strasbourg, France. Strasbourg Federation of Translational Medicine (FMTS), Strasbourg University, Strasbourg, France; INSERM U1258 / CNRS UMR7104, Institut de Génétique et de Biologie Moléculaire et Cellulaire, Illkirch, France; Sorbonne Université, INSERM UMR_S 1166 ICAN, Paris, France; Service d’Immunologie Biologique, Plateau Technique de Biologie, Pôle de Biologie, Nouvel Hôpital Civil, Hôpitaux Universitaires de Strasbourg, Strasbourg, France; Department of Dermatology, Venereology and Allergology, Medical University of Innsbruck, Innsbruck, Austria; Institute for Molecular Medicine and Paul Klein Center for Immunotherapy (PKZI), University Medical Center of the Johannes Gutenberg-University Mainz, Mainz, Germany; INSERM UMR_S 1109 Immunorhumatologie Moléculaire, Fédération de Médecine Translationnelle de Strasbourg (FMTS), ITI Transplantex NG, Centre de Recherche en Biomédecine de Strasbourg (CRBS), Faculté des Sciences de la Vie, Université de Strasbourg, Strasbourg, France

**Keywords:** Autophagy, Langerhans cells, skin, homeostasis, lipid droplets, ferroptosis, metabolism

## Abstract

Macroautophagy (often-named autophagy), a catabolic process involving autophagy-related (*Atg*) genes, prevents accumulation of harmful cytoplasmic components and mobilizes energy reserves in long-lived and self-renewing cells. Autophagy deficiency affects antigen presentation in conventional dendritic cells (DCs) without impacting their survival. However, previous studies did not address epidermal Langerhans cells (LCs). Here, we demonstrate that deletion of either *Atg5* or *Atg7* in LCs leads to their gradual depletion. ATG5-deficient LCs showed metabolic dysregulation and accumulated neutral lipids. Despite increased mitochondrial respiratory capacity, they were unable to process lipids, eventually leading them to ferroptosis. Finally, metabolically impaired LCs upregulated proinflammatory transcripts and showed decreased expression of neuronal interaction receptors. Altogether, autophagy represents a critical regulator of lipid storage and metabolism in LCs, allowing their maintenance in the epidermis.

## INTRODUCTION

Langerhans cells (LCs) are resident antigen-presenting cells (APCs) of the epidermis^1,2^. LCs arise from hematopoietic precursors that emerge from the yolk sac and the fetal liver to colonise the skin before birth^3^. There, they are maintained lifelong by local proliferation^4^. LCs exhibit an exceptional longevity, with a half-life of several weeks. In contrast, conventional dendritic cells (cDCs), which represent a major skin APC subset, are replenished from bone marrow precursors within days^5^. Possibly as a consequence of UV exposure, LCs are endowed with a potent DNA-repair capacity, allowing the survival of at least a pool of self-renewing cells upon gamma irradiation^6^. Despite free diffusion of glucose from the blood into the lowest layers of the epidermis, their position in the suprabasal layers implies a limited supply of nutrients, which must be metabolised in a very hypoxic environment^7^. Similar to cutaneous DC subsets, LCs migrate to lymph nodes (LNs) following microorganism recognition or irradiation. There, LCs are important contributors to antigen presentation and differentiation of CD4^+^ and CD8^+^ T cells, either driving immune activation or tolerance^8–10^. LCs are among the first APCs that sense skin infections^11^ and are involved in inflammatory disorders such as psoriasis^12^. Therefore, a deeper understanding of their homeostasis appears critical.

Autophagy is a conserved mechanism of self-digestion, allowing the engulfment of cytoplasmic content into double-membrane vesicles, which fuse with lysosomes for degradation and recycling of the sequestered content^13,14^. The core autophagy proteins are encoded by autophagy-related (*Atg*) genes. Autophagy is promoted under energetic stress notably through the inhibition of the PI3K/AKT/mTOR pathway^15^. Autophagy also contributes to metabolic equilibrium in homeostatic conditions as it is a key process to support energy provision. For cells relying on oxidative phosphorylation to generate ATP, autophagy contributes to maintain a functional mitochondrial pool through degradation of defective mitochondria and to mobilize fatty acids through degradation of lipid droplets in a process called lipophagy^16^. To support homeostasis, autophagy also acts as a quality-control mechanism during the unfolded protein response (UPR), preventing the accumulation of misfolded protein aggregates and degrading excess or damaged endoplasmic reticulum (ER)^17^. These housekeeping forms of autophagy are particularly important in long-lived and self-renewing cells. In the immune system, B-1 B cells, memory B and T cells as well as plasma cells rely on autophagy for their maintenance^13,18–22^. ATG proteins are also involved in several non-autophagic processes such as LC3-associated phagocytosis (LAP). LAP requires Rubicon (*Rubcn*) to form an initiation complex, and is involved in microorganism clearance, efferocytosis and antigen presentation, which are highly relevant for DCs and macrophages^23,24^. Notably, autophagy impairment in DCs notably leads to defective CD4^+^ and CD8^+^ T cell responses^25–28^.

Overall, selective deletion of *Atg* genes in macrophages and DCs has demonstrated that autophagy modulates pathogen resistance, antigen presentation, and proinflammatory signals, i.e. inflammasome activity^29–32^. Similarly, recent reports support a role of autophagy for LCs in the regulation of inflammatory responses^33,34^ and in the immune response against intracellular bacteria^35^. Moreover, autophagy proteins participate in the intracellular routing of human immunodeficiency virus (HIV) particles towards degradative compartments in human LCs upon Langerin/CD207-mediated uptake^36^. Interestingly, enhancing autophagy by pharmacological agents limits HIV-1 mucosal infection and replication^37^. Yet, non-autophagic roles of ATG proteins cannot be ruled out to explain these results.

When autophagy defects were assessed *in vivo* for cDCs and macrophages, there was no report of an impaired cell survival^26,27,38,39^, except for a peritoneal macrophage subset^40^ Although some of the conditional deletion systems used for these investigations may have impacted LCs as well, no information is currently available on the consequences of constitutive autophagy impairment for their maintenance *in vivo*. Since LCs are self-renewing, long-lived APCs that are exposed to low availability of nutrients, UV irradiation or stress related to infection, we hypothesized that efficient autophagy might be a key element supporting their maintenance in the epidermis. To investigate this, we generated *Cd207*-specific deletion of *Atg5* in order to define primary roles of autophagy and related processes in LC biology.

## RESULTS

### ATG5 is necessary for Langerhans cell network maintenance

Since evidence for autophagosomes in primary LCs has been so far limited ^36^, we first verified whether LCs from digested murine epidermis comprise such compartments. Electron microscopy of LCs, including original images and reanalysis of previously published samples^41^, allowed the identification of double-membrane compartments as well as crescent-shape structures reminiscent of incipient phagophores and isolation membranes. The diameter of the autophagosomes was between 400 and 600nm **(Figure S1)**. In line with this observation, immunofluorescence revealed LC3-positive compartments within LCs **(Figure 1a, *Atg5^W^****^T^*). When LCs were treated with hydroxychloroquine to block lysosomal degradation of autophagosomes, we observed an accumulation of membrane-associated LC3 by flow cytometry, thereby demonstrating autophagic flux **(Figure 1b, *Atg5^W^****^T^*). Altogether, this showed constitutive autophagic activity in LCs of wild-type mice.

**Figure 1:**
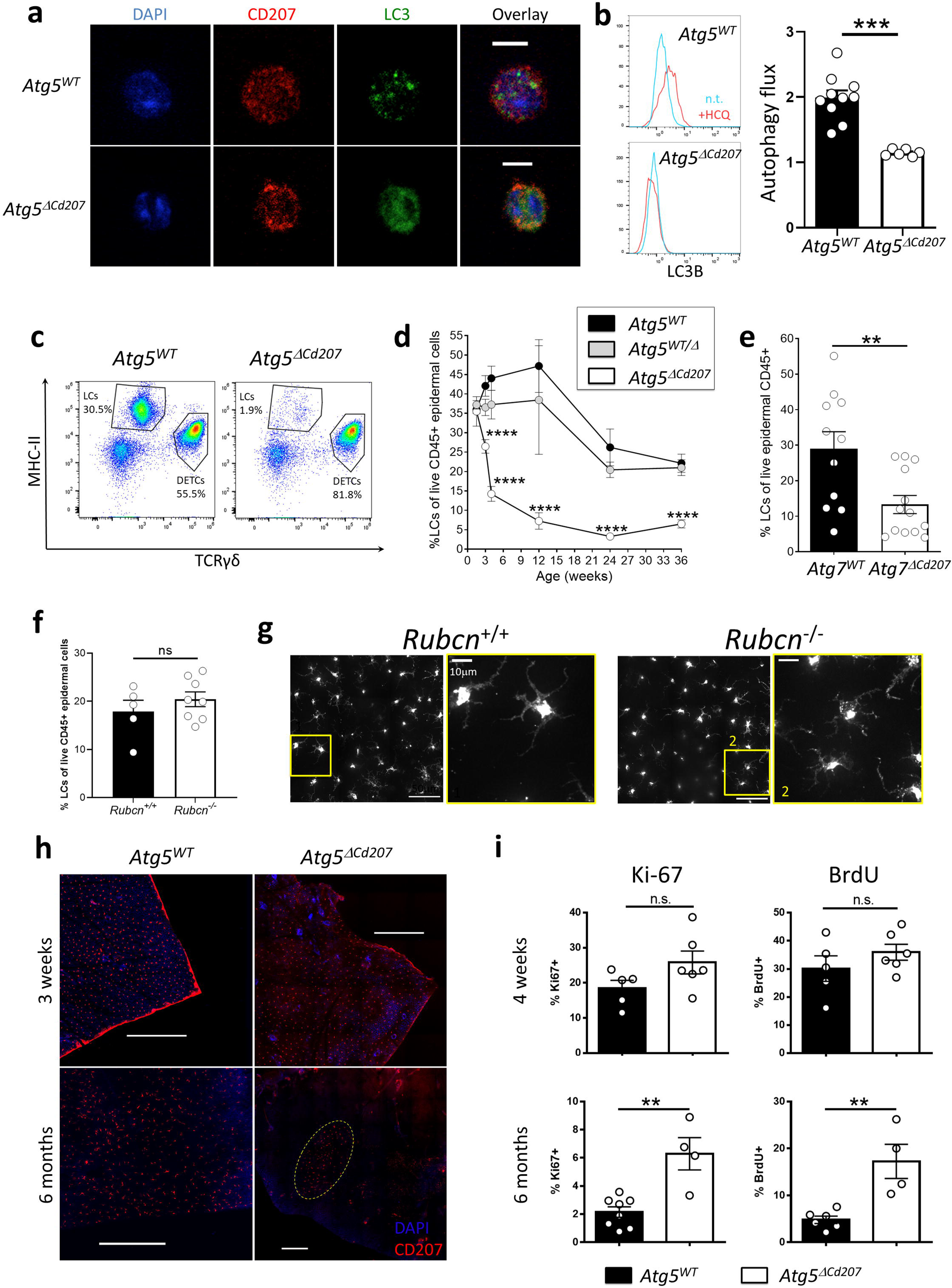
ATG5 deficiency in Langerhans cells disrupts autophagosomes and depletes their epidermal network. **(a)** Representative immunofluorescent stainings of MAP1LC3B (LC3) on LCs obtained by *in vitro* emigration from epidermal sheets of *Atg5^WT^* **(Supplementary Video SV1)** and *Atg5*^Δ*Cd207*^ **(Supplementary Video SV2)** mice. Scale bars: 10µm. **(b)** Representative histogram plot of LC3B staining and quantification of mean fluorescence intensity (MFI) for LCs of 3-week-old *Atg5^WT^* and *Atg5*^Δ*Cd207*^ mice, treated or not with chloroquine. Autophagy flux was calculated as a ratio between MFI for LC3B in treated and untreated cells. **(c)** Representative dot plots for the identification of MHCII^+^ TCRγδ^-^ Langerhans cells (LCs; all CD207^+^) and MHCII^-^ TCRγδ^+^ dendritic epidermal T Cells (DETCs) among CD45+ cells in freshly digested back skin epidermal suspension of 6-month-old *Atg5^WT^* and *Atg5*^Δ*Cd207*^ mice. **(d)** Comparison over time of the percentage of LCs among live CD45^+^ epidermal cells for control (*Atg5^WT^* and *Atg5^WT/^*^Δ^) and *Atg5*^Δ*Cd207*^ mice. **(e)** Percentage of LCs among live CD45+ cells in freshly digested back skin epidermis of 10-20 week-old *Atg7^WT^* and *Atg7*^Δ*Cd207*^ mice. **(f)** Percentage of CD45+ MHCII+ CD207+ LCs among live CD45+ cells obtained from fresh epidermal cell suspensions of 2-month-old *Rubicn*^-/-^ and *Rubicn*^+/+^ mice. **(g)** Immunofluorescence staining of CD207+ LCs in ear epidermal sheets of 2-month-old *Rubicn^-/-^* and *Rubicn^+/+^* mice. Images are representative of two independent experiments. Insets 1 and 2: close-up views. **(h)** Representative immunofluorescent staining of CD207 on epidermal sheets of ear skin from 3-week (upper panels) and 6-month-old (lower panels) *Atg5^WT^* and *Atg5*^Δ*Cd207*^ mice. Scale bars: 100µm. **(i)** Percentages of epidermal LCs stained for BrdU incorporation and Ki67 expression for 4-week (top) and 6-month-old (bottom) *Atg5^WT^* and *Atg5*^Δ*Cd207*^ mice. All data are pooled from at least 3 independent experiments, with each point representing one individual mouse (except d: n≥4 mice per time-point). Statistical analysis: Mann-Whitney U test (except (d): two-way ANOVA followed by Tukey’s multiple comparison test). **, p<0.01; ***, p<0.001; ****, p<0.0001; ns, p>0.05.

To determine the function of autophagy in LCs *in vivo* we generated *Cd207-cre x Atg5^flox/-^*(*Atg5*^Δ*Cd207*^) mice, in which the essential autophagy gene *Atg5* is deleted by CRE-mediated recombination in cells expressing CD207 **(Figure S2a)**. Efficiency of the deletion was verified by RT-qPCR of LCs sorted from the mouse epidermis **(Figure S2b)** and from skin-draining LNs of 4-week-old mice **(Figure S2c,d)**. This confirmed that the breeding strategy resulted in an optimal deletion efficiency, as *Atg5* mRNA was undetectable in LCs from *Atg5*^Δ*Cd207*^ mice, as compared with LCs from *Atg5^flox/+^* and *Cd207-cre x Atg5^flox/+^*control animals (respectively referred to as *Atg5^WT^* and *Atg5^WT/^*^Δ^ below). With respect to migratory dermal DCs isolated from LNs of *Atg5*^Δ*Cd207*^ mice, *Atg5* mRNA was absent from CD103^+^ dermal cDC1, which also express CD207^42^, but still present in CD207^-^ MHCII^high^ dermal DCs **(Figure S2d)**.

To address whether the *Atg5* deletion leads to autophagy impairment in LCs, formation of autophagic compartments was assessed by LC3 immunostaining. LC3^+^ punctate staining in the cytoplasm of *Atg5^WT^* LCs was clearly visible, whereas LC3 staining was diffuse in LCs from *Atg5*^Δ*Cd207*^ mice **(Figure 1a, and Supplementary Videos SV1, SV2)**. This reflects the expected consequences of ATG5 deficiency, i.e. the absence of LC3 conjugation with phosphatidylethanolamine (LC3-II) and lack of integration into autophagic compartments. Since this accumulation of LC3^+^ vesicles may also reveal an impaired degradation of autophagosomes, the acidification and lysosomal load were quantified by Lysosensor and Lysotracker probes, respectively. We could thus verify that it was unperturbed in the absence of ATG5 **(Figure S3)**. Finally, we observed that autophagic flux was abolished in LCs of *Atg5*^Δ*Cd207*^ mice **(Figure 1b)**. This shows that ATG5 deletion leads to autophagy impairment in LCs.

To monitor the possible involvement of autophagy in the homeostatic maintenance of LCs under steady state conditions, we evaluated their epidermal network at different ages. Since CD207 expression in MHCII^+^ epidermal LC precursors is completed around 7-10 days after birth^43^, we assessed the proportion of MHCII^+^ TCRγδ^-^ CD207^+^ LCs among CD45^+^ epidermal cells by flow cytometry from 10 days until 9 months of age **(Figure 1c)**. The basal proportion of LCs at 10 days was comparable for mice of all genotypes, suggesting that no major defect occurs in the seeding of MHCII^+^ CD207^-^ embryonic LC precursors in the epidermis, which also corresponds to the expected kinetics of *Cd207* promoter activity and CRE expression^43^ **(Figure 1d)**. In *Atg5^WT^* and *Atg5^WT/^*^Δ^ mice, we observed a moderate increase of LCs until 6-12 weeks, followed by a decrease in aging mice. Nevertheless, the proportion of LCs diminished sharply around 2-4 weeks of age in the epidermis of *Atg5*^Δ*Cd207*^, eventually stabilizing at around 5% of epidermal CD45^+^ leukocytes at 9 months.

To reinforce the conclusion that the loss of LCs is due to impaired autophagy and not other ATG5-related cellular homeostatic dysfunctions, we generated *Cd207-cre x Atg7^flox/flox^* (*Atg7*^Δ*Cd207*^) mice and compared their epidermal cell suspensions with that of *Atg7^flox/flox^*(*Atg7^WT^)* mice. Similar to our findings with *Atg5*^Δ*Cd207*^ mice, ATG7 deficiency resulted in a significant depletion of LCs from the epidermis of mice older than 10 weeks **(Figure 1e)**. Thus, we can exclude effects only linked to ATG5, such as direct regulation of apoptosis independently of the autophagy machinery^15^. Additionally, we could exclude a role for LAP or other endocytic processes requiring ATG5, as the density of the epidermal network of LCs appeared unaffected in Rubicon-deficient mice **(Figure 1f,g).**

We next performed an immunofluorescent staining of the LC network in epidermal sheets prepared from ear skin of *Atg5*^Δ*Cd207*^ and control mice **(Figure 1h)**. We did not observe any obvious difference in the LC network in mice, regardless of their genotype. However, and in accordance with flow cytometry results, very few LCs were visible in 6-month-old *Atg5*^Δ*Cd207*^ mice. LCs remaining in older mice were often assembled in disseminated patches. This pattern is reminiscent of the network reconstitution that occurs through slow *in situ* LC proliferation following induced depletion in Langerin-DTR mice^44^. Indeed, LCs ensure the integrity of their epidermal network by self-renewal^45,46^. Consequently, we assessed the proliferative capacity of ATG5-deficient LCs by 5-bromo-2’-deoxyuridine (BrdU) incorporation and Ki-67 staining by flow cytometry **(Figure 1i)**. We observed proliferation rates consistent with our previous observations in 4-week-old mice^47^, with comparable percentages of BrdU^+^ and Ki-67^+^ LCs in *Atg5*^Δ*Cd207*^ and control *Atg5^WT^* mice, thereby concluding that autophagy deficiency does not prevent maintenance of the LC network by a major proliferative impairment. On the other hand, LCs of 6-month-old *Atg5*^Δ*Cd207*^ mice displayed higher proliferation rates, presumably because of homeostatic compensation for the depletion of their epidermal niche.

Finally, since dermal cDC1 also express CD207 and showed a decrease of *Atg5* transcripts in *Atg5*^Δ*Cd207*^ mice **(Figure S2d)**, we quantified LC and cDC1 populations among total MHCII^+^ CD11c^+^ skin DCs **(Figure S2e)**. We confirmed the significant decrease of LCs present in whole skin cell suspensions, while the proportion of cDC1 among skin DCs was rather slightly increased **(Figure S2f)**. Thus, the core autophagic machinery is dispensable for the maintenance of dermal cDC1, yet appears essential to LCs.

### ATG5-deficient Langerhans cells undergo limited apoptosis

Since their self-renewal was not affected, the loss of ATG5-deficient LCs might be explained by an enhanced migration into lymph nodes or by increased cell death. LCs, similar to cDCs of peripheral tissues, undergo maturation and migrate to skin-draining LNs following inflammatory signals^1,2^. Alternatively, spontaneous maturation of LCs may result from disrupted TGF-β signalling, which, under physiological conditions, is required to maintain an immature state^48,49^. In both cases, an increased expression of maturation markers MHC-II and CD86 can be observed prior to the departure of LCs from the epidermis. We thus verified whether autophagy impairment might prompt spontaneous LC maturation. MHC-II and CD86 expression by LCs in the epidermis did not show any variation in *Atg5*^Δ*Cd207*^ compared to control mice **(Figure 2a,b)**. Moreover, because an overt LC emigration from the epidermis would lead to a noticeable accumulation in LNs, we determined LC numbers in inguinal and brachial LNs of 6-week-old mice. We observed instead a trend towards a decrease, only significant in percentage for LCs from *Atg5*^Δ*Cd207*^ mice **(Figure 2c,d)**. A similar pattern was observed for dermal cDC1 that also express CD207. In contrast, no differences were detected for CD207^-^ dermal DC subsets that lack *Atg5* deletion. Taken together, these results exclude that impaired autophagy leads to a massive spontaneous maturation and migration of LCs.

**Figure 2:**
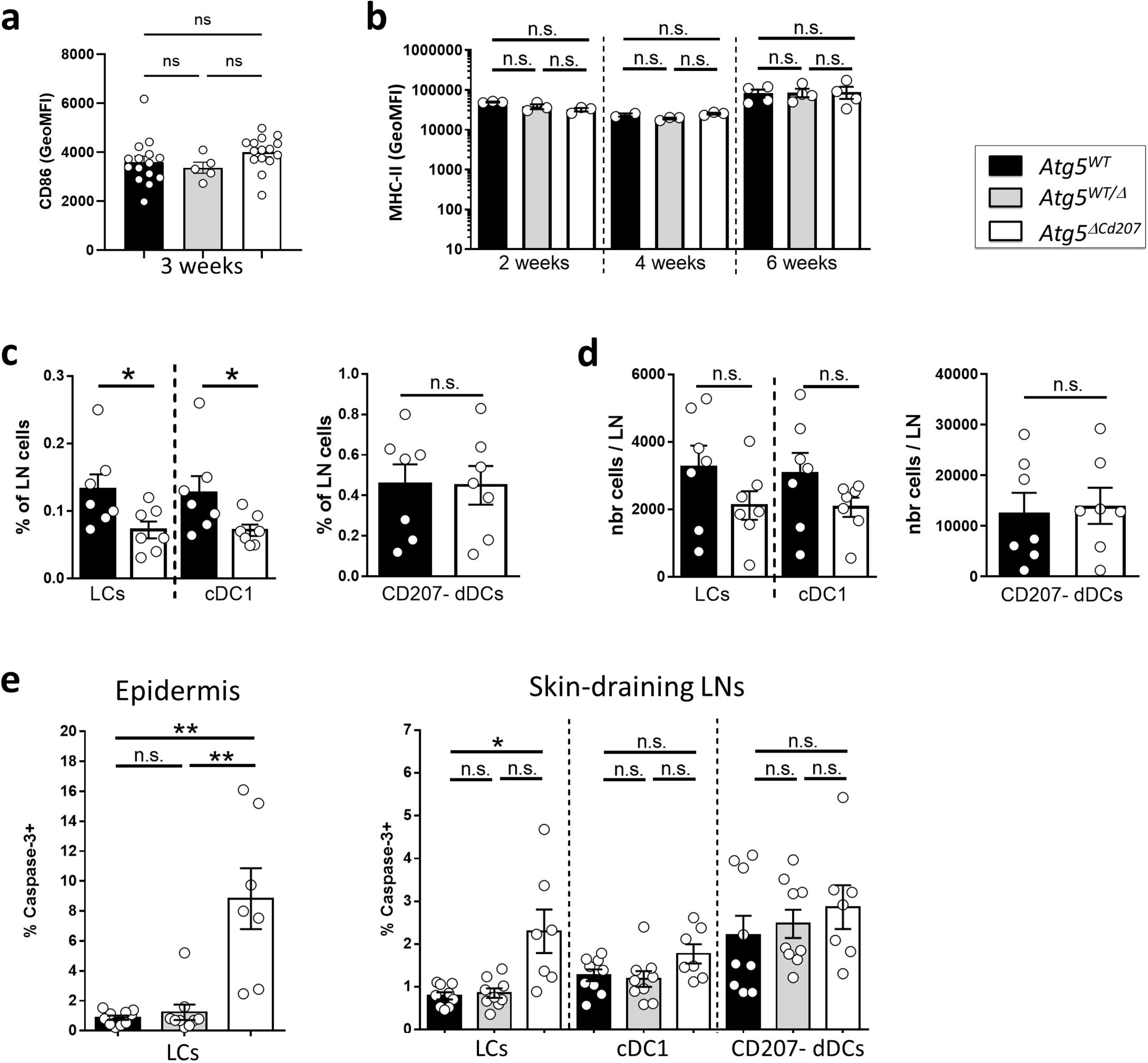
ATG5-deficient Langerhans cells remain immature and undergo apoptosis. **(a)** CD86 and **(b)** MHC-II MFI for epidermal LCs of 3-week-old control (*Atg5^WT^* and *Atg5^WT/^*^Δ^) and *Atg5*^Δ*Cd207*^ mice. Data are pooled from 5 independent experiments, with each point representing one individual mouse. Statistical analysis: Kruskal-Wallis one-way ANOVA followed by Dunn’s multiple comparison test (ns, p>0.05). **(c)** Percentages and **(d)** absolute numbers of LCs, cDC1 and CD207^-^ DCs in freshly digested skin draining lymph nodes of 3-week-old control (*Atg5^WT^* and *Atg5^WT/^*^Δ^) and *Atg5*^Δ*Cd207*^ mice. Data are pooled from at least 3 independent experiments, with each point representing one individual mouse. Statistical analysis: Mann-Whitney U test (*, p<0.05; ns, p>0.05). **(e)** Percentage of cells with activated caspase-3 in LCs of freshly digested back skin epidermis (left panel) and LCs, cDC1 and CD207-dermal DCs of skin-draining lymph nodes (right panel) for 3-week-old control (*Atg5^WT^* and *Atg5^WT/^*^Δ^) and *Atg5*^Δ*Cd207*^ mice. Data are pooled from at least 3 independent experiments, with each point representing one individual mouse. Statistical analysis: Kruskal-Wallis one-way ANOVA followed by Dunn’s multiple comparison test (*, p<0.05; **, p<0.01; ns, p>0.05).

Finally, using flow cytometry, we addressed whether the absence of *Atg5* might lead to increased apoptosis by measuring the proportion of LCs with active caspase-3. This major effector of apoptosis was detected in a small, yet significant proportion of ATG5-deficient LCs, both in the epidermis and in LNs **(Figure 2e)**. Altogether, these results demonstrate that reduced cell division or increased maturation and migration cannot account for LC network disintegration, whereas apoptosis of ATG5-deficient LCs, albeit limited, suggests autophagy as crucial, even in the steady state, for LC survival.

### ATG5-deficient Langerhans cells show endoplasmic reticulum swelling but no unfolded protein response

Functional autophagy is required for the maintenance of the endoplasmic reticulum (ER). ER damage triggers the inositol-requiring enzyme 1 (IRE1)/X-box binding protein 1 (Xbp1) axis of the Unfolded Protein Response (UPR), which is a master regulator of DC survival and maturation^50–52^ but has not been investigated in LCs so far. Autophagy regulates ER swelling, protein aggregation and thereby limits the extent of the UPR^53^. In line with this, exposure of wild-type LCs to the phosphatidylinositol-3-kinase inhibitor, wortmannin, which inhibits the initiation of the autophagosome formation, resulted in an increased labelling by ER-tracker, a fluorescent dye specific for ER membranes **(Figure 3a)**. Then, we stained *Atg5*-deficient LCs from 3-week-old mice with ER-tracker. Flow cytometry analysis showed a significantly increased ER-tracker staining in LCs of *Atg5*^Δ*Cd207*^ compared to wild-type mice **(Figure 3b)**. In line with this, confocal microscopy revealed an expanded ER compartment in these cells **(Figure 3c and Supplementary Videos SV3, SV4)**. These signs of ER expansion prompted us to study whether the expression of key intermediates of the UPR pathway might be elevated. Quantitative PCR was performed for *Ern1*, total *Xbp1*, spliced *Xbp1* and *Ddit3* mRNA. However, none of these genes showed increased expression, demonstrating that the UPR pathway was not constitutively engaged **(Figure 3d)**. Therefore, we conclude that ATG5-deficient LCs can cope with the observed ER swelling, which does not trigger a massive UPR that could lead to cell death.

**Figure 3:**
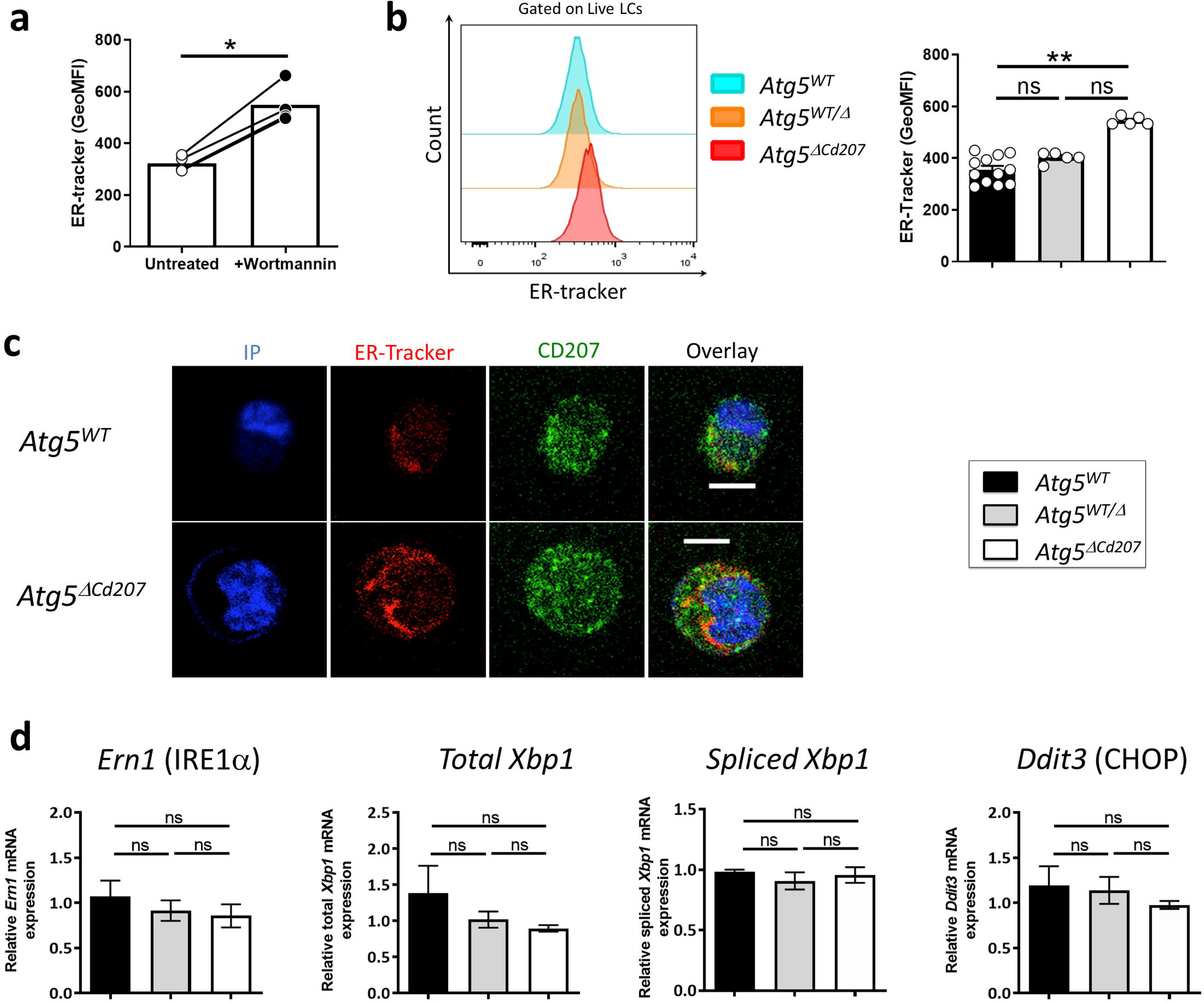
ATG5-deficient Langerhans cells present endoplasmic reticulum swelling but no unfolded protein response. **(a)** MFI for ER-tracker on epidermal LCs treated or not with Wortmannin. Data are pooled from at least 3 independent experiments, with each point representing one individual 3-week-old mouse. Statistical analysis: Mann-Whitney U test (*, p<0.05). **(b)** Representative half-set overlay (left panel) and MFI (right panel) of ER-tracker in epidermal LCs of control (*Atg5^WT^* and *Atg5^WT/^*^Δ^) and *Atg5*^Δ*Cd207*^ mice. Data are pooled from at least 3 independent experiments, with each point representing one individual 3-week-old mouse. Statistical analysis: Kruskal-Wallis one-way ANOVA followed by Dunn’s multiple comparison test (**, p<0.01). **(c)** Representative immunofluorescent stainings of the endoplasmic reticulum using the ER-tracker dye on epidermal LCs of 3-week-old *Atg5^WT^* **(Supplementary Video SV3)** and *Atg5*^Δ*Cd207*^ **(Supplementary Video SV4)** mice. Scale bar: 10µm. **(d)** Expression of *Ern1*, total and spliced *Xbp1*, and *Ddit3* mRNAs, in epidermal LCs of 3-week-old control (*Atg5^WT^* and *Atg5^WT/^*^Δ^) and *Atg5*^Δ*Cd207*^ mice. Fold changes were calculated relative to mRNA expression in cells of *Atg5^WT^* control mice. Data are pooled from at least 3 independent experiments, with each point representing one individual mouse. Statistical analysis: Kruskal-Wallis one-way ANOVA followed by Dunn’s multiple comparison test (ns, p>0.05).

### Autophagy-deficient Langerhans cells accumulate intracellular lipid storage

To identify dysregulated gene expression patterns that could be linked with impaired autophagy, RNA sequencing was performed on epidermal LCs sorted from 3-week-old *Atg5*^Δ*Cd207*^ or *Atg5^WT^* control mice. Analysis of *Atg5* mRNA sequencing reads confirmed the deletion of exon 3 in LCs upon CRE-mediated recombination **(Figure S4a)**. Principal component analysis revealed clear differences in transcriptomic profiles between *Atg5*^Δ*Cd207*^ and *Atg5^WT^* mice **(Figure S4b)**. Differentially expressed genes in *Atg5*^Δ*Cd207*^ LCs included 673 upregulated and 629 downregulated genes **(Table 2, Figure S4c,d)**. Gene ontology pathway enrichment analysis suggested in particular a dysregulation of cellular metabolism (GO:0046942, GO:0043269, GO:0017144, GO:0044272, GO:0051186, GO:0110096, GO:0046085, GO:1901615, GO:0015711, GO:0009166, GO:0007584; **Figure S4e)**.

Autophagy regulates cellular lipid metabolism by lipophagy, which has a crucial role in balancing energy supply in both steady state and under metabolic stress. Lipophagy mediates lysosomal degradation of proteins that coat cytoplasmic lipid droplets, and lipolysis of triglycerides, thus liberating free fatty acids to be consumed by beta-oxidation in mitochondria^54^. Accordingly, LCs of *Atg5*^Δ*Cd207*^ mice modulated the expression of several genes encoding actors of lipidic metabolism pathways **(Figure 4a)**. We noticed upregulation of mRNA of the solute carrier (SLC) family transporters MCT-4/SLC16A3 (lactate), SLC7A11 (cystein, glutamate) and SLC7A2 (lysine, arginine), which import molecules that directly or indirectly provide substrates to the tricarboxylic acid cycle. Upregulated expression of *Acss1* and *Acss2* (Acyl-CoA Synthetase Short Chain Family Member 1 and 2) is expected to favour synthesis of Acetyl-CoA, which can either be converted into lipids or fuel mitochondrial beta-oxidation. Fatty acid synthesis and energy storage in the form of lipid droplets appears to be favoured in ATG5-deficient LCs, as hinted by the upregulation of *Gyk*, encoding the Glycerol kinase which catalyses triglyceride synthesis, and *Acsl3,* encoding the Acyl-CoA Synthetase Long Chain Family Member 3, a key enzyme for neutral lipid generation^55^.

**Figure 4:**
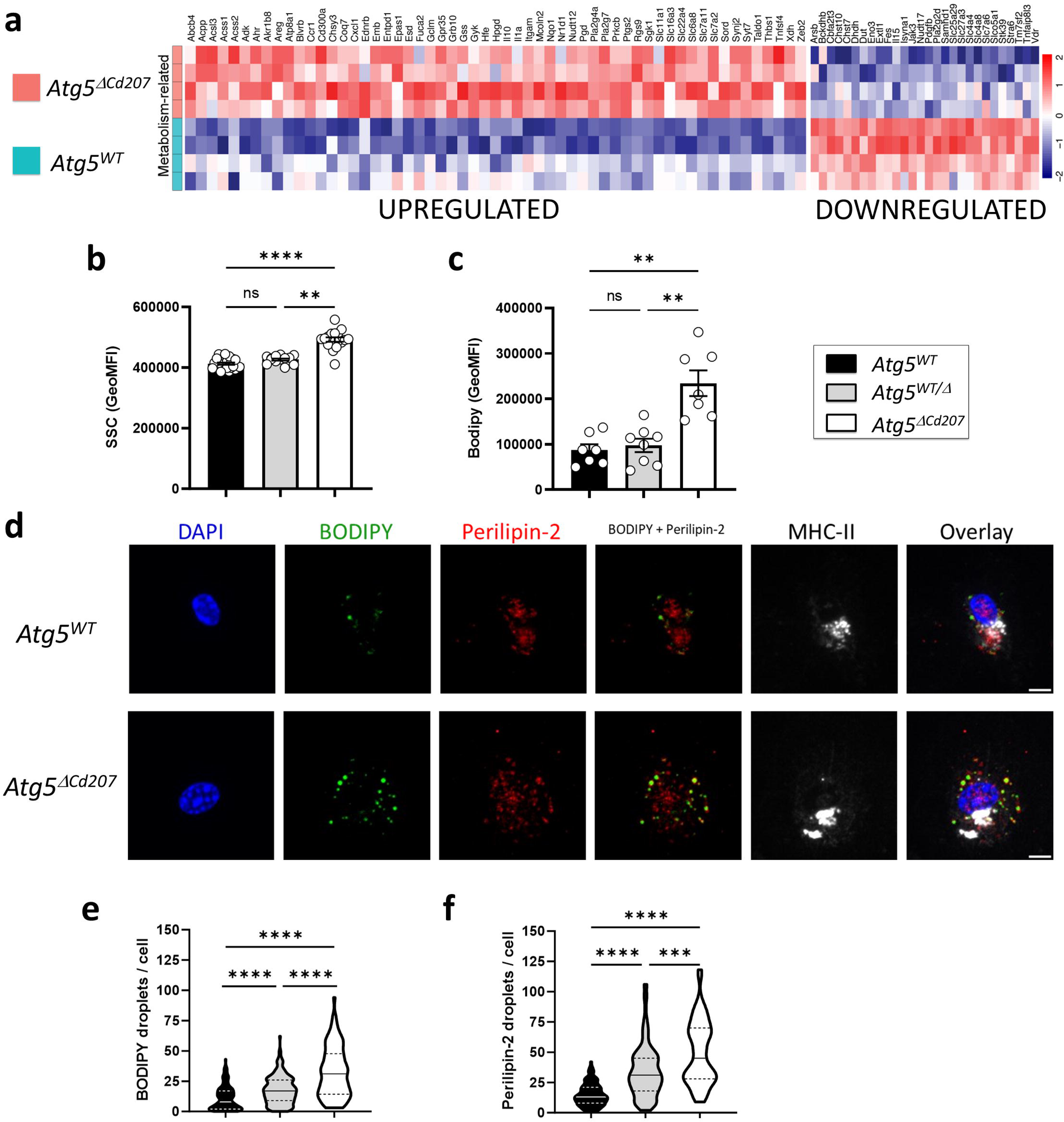
Impaired autophagy increases the lipid storage compartments of Langerhans cells. **(a)** Differentially expressed transcripts related to lipid metabolism pathways in *Atg5^WT^* vs. *Atg5*^Δ*Cd207*^ LCs (GO:0046942, GO:0043269, GO:0017144, GO:0044272, GO:0051186, GO:0110096, GO:0046085, GO:1901615, GO:0015711, GO:0009166, GO:0007584). **(b-c)** Flow cytometry quantification of the side scatter (b) and Bodipy (C) MFI in epidermal LCs obtained from control (*Atg5^WT^* and *Atg5^WT/^*^Δ^) and *Atg5*^Δ*Cd207*^ mice. Data are pooled from 10 (SSC) or 4 (Bodipy) independent experiments, with each point representing one individual 3-week-old mouse. Statistical analysis: Kruskal-Wallis one-way ANOVA followed by Dunn’s multiple comparison test (*, p<0.05; **, p<0.01; ns, p>0.05). **(d)** Immunofluorescent stainings of MHCII+ epidermal LCs obtained from *Atg5^WT^* (upper panels) and *Atg5*^Δ*Cd207*^ (lower panels) mice and stained with Bodipy or anti-Perilipin-2 antibody. Scale bars: 10µm. **(e-f)**. Quantification of (E) Bodipy+ and (F) Perilipin- 2+ vesicles in LCs obtained from control (*Atg5^WT^* and *Atg5^WT/^*^Δ^) and *Atg5*^Δ*Cd207*^ mice. Data are representative of 3 independent experiments and presented as violin plots (solid line, median; dashed lines, first and third quartiles; n>100 cells from a total of three 3-week-old mice). Statistical analysis: Kruskal-Wallis one-way ANOVA followed by Dunn’s multiple comparison test ***, p<0.001; ****, p<0.0001).

Using flow cytometry, we noticed an increased side scatter of *Atg5*^Δ*Cd207*^ LCs **(Figure 4b)**. Considering that lipid metabolism appeared deregulated in autophagy-deficient LCs, we hypothesized that this increased granularity may be due to a larger number of intracellular lipid-rich vesicles. Thus, epidermal cell suspensions were exposed to Bodipy, a lipid staining dye that targets neutral lipid-rich compartments, which we first quantified by flow cytometry. We first validated this experimental approach by treating wild-type LCs with etomoxir, a carnitine palmitoyltransferase I inhibitor that blocks the import of activated free fatty acids (acyl-CoA) by mitochondria. Etomoxir treatment resulted in a stronger intensity of Bodipy staining, reflecting higher neutral lipid storage by LCs **(Figure S5a)**. Treating LCs with the autophagy inhibitor wortmannin also resulted in an increased Bodipy staining, suggesting constitutive lipophagy in LCs **(Figure S5b)**. We then found that *Atg5*^Δ*Cd207*^ LCs retained more Bodipy as compared with LCs of control mice **(Figure 4c)**. We consistently observed by confocal microscopy that LCs of *Atg5*^Δ*Cd207*^ mice contained more Bodipy-positive vesicular structures that could correspond to lipid droplets **(Figure 4d,e and Supplementary Videos SV5, SV6)**. To unequivocally identify this lipid storage compartment, we interested in perilipins, a family of lipid droplet-specific proteins. According to Immgen gene expression datasets^56^, *Plin2*, encoding Perilipin-2, is the most strongly expressed in LCs among this family. Staining for Perilipin-2 lipid droplets confirmed their accumulation in autophagy-deficient LCs **(Figure 4d,f)**. Both Bodipy+ and Perilipin-2+ vesicles were already increased in LCs of *Atg5^WT/^*^Δ^ mice, as compared to *Atg5^WT^* mice **(Figure 4e,f)**, suggesting that even a moderate impairment of autophagy results in an abnormal increase of neutral lipid storage compartments. Of note, the enlargement of ER **(Figure 3)** could be linked to the increased production of lipid droplets, which originate from this compartment^57^.

### Disrupted lipid metabolism in autophagy-deficient Langerhans cells

ATG5 deficiency might lead to an accumulation of intracellular lipids if energy production in LCs strongly relies on lipophagy to mobilize these storage units and produce energy by the beta-oxidation pathway^54^. To assess the energy production in LCs, we focused on AMP-activated protein kinase (AMPK), a master regulator of energetic metabolism, which is phosphorylated on residues T183/T172 when ATP/AMP ratios decline^58^. As measured by flow cytometry, AMPK phosphorylation was indeed increased in LCs from *Atg5*^Δ*Cd207*^ mice **(Figure 5a)**. AMPK phosphorylation is expected to promote different pathways that help restore optimal ATP production, including import of glucose or fatty acid uptake and synthesis. To quantify the glucose uptake intensity, we first monitored the expression of the glucose transporter GLUT-1 by LCs. However, no difference could be observed between LCs of *Atg5*^Δ*Cd207*^ mice and control mice **(Figure 5b)**. Next, we quantified the glucose uptake of these cells using 2-(N-(7-Nitrobenz-2-oxa-1,3-diazol-4-yl) Amino)-2-Deoxyglucose (2-NBDG), a fluorescent glucose analogue which can be tracked by flow cytometry. In line with the unmodified GLUT-1 expression, autophagy-deficient LCs were not more efficient at capturing glucose than LCs of wild-type mice **(Figure 5c)**. On the other hand, LCs from *Atg5*^Δ*Cd207*^ mice exhibited a stronger expression of CD36 **(Figure 5d)**. This scavenger receptor has a key role in the capture of free fatty acids and lipids, which was indeed increased when LCs incubated with Bodipy-labelled C16 fatty acid **(Figure 5e)**. Altogether, these assays demonstrate that autophagy-deficient LCs show a deficit in energy production, despite an increased ability to capture and store lipids.

**Figure 5:**
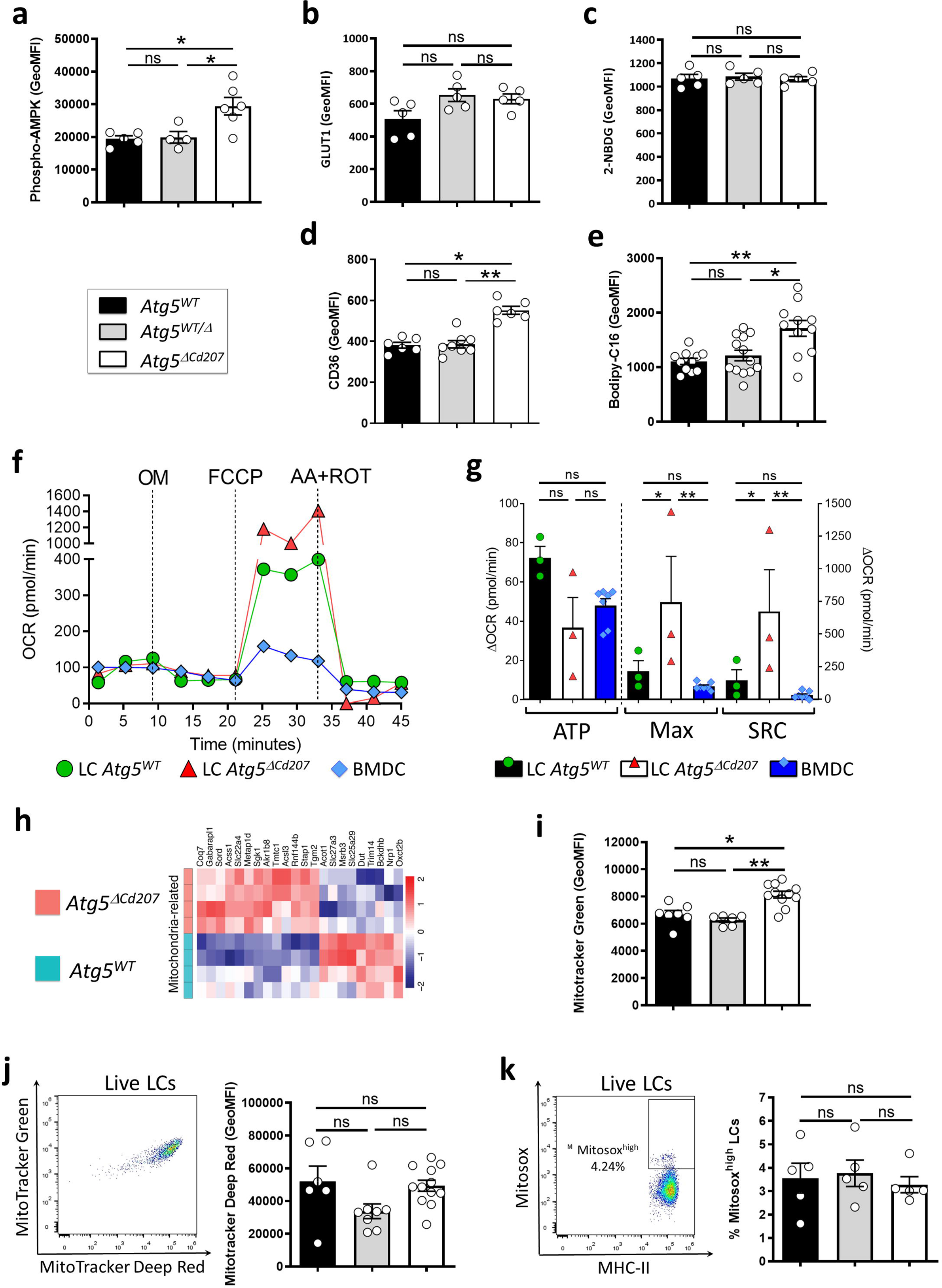
Impaired lipid metabolism in ATG5-deficient Langerhans cells. **(a-e)** Flow cytometry quantification of mean intensity of fluorescence for **(a)** Phosphorylated AMPK, **(b)** GLUT1, **(c)** 2-NDBG uptake, **(d)** CD36 and **(e)** Bodipy C16 uptake in epidermal LCs obtained from 3-week-old control (*Atg5^WT^* and *Atg5^WT/^*^Δ^) and *Atg5*^Δ*Cd207*^ mice. **(f)** Epidermal LCs sorted from *Atg5^WT^* or *Atg5*^Δ*Cd207*^ mice and BMDCs derived from C57BL/6 mice were sequentially exposed to Oligomycin (OM), Carbonyl cyanide 4- (trifluoromethoxy)phenylhydrazone (FCCP), rotenone (ROT) and antimycin A (AA), and oxygen consumption rates (OCR) were measured by a Seahorse XF96 analyzer throughout the experiment. Data are from one representative experiment out of three. **(g)** ATP production (OCR_baseline_ – OCR_OM_), maximum respiration (Max; OCR_FCCP_ – OCR_AA+ROT_) and spare respiratory capacity (SRC; OCR_FCCP_ – OCR_baseline_) were calculated from the OCR curves. **(h)** Differentially expressed transcripts related to mitochondria (GO: 0005739) in *Atg5^WT^* vs. *Atg5*^Δ*Cd207*^ LCs. **(i)** Mitochondrial load for epidermal LCs of *Atg5^WT^*, *Atg5^WT/^*^Δ^ and *Atg5*^Δ*Cd207*^ mice, as measured by MFI of Mitotracker Green staining. **(j)** Representative dot plot of Mitotracker Green and Deep-Red staining and comparison of Mitotracker Deep-Red MFI of epidermal LCs obtained from control (*Atg5^WT^* and *Atg5^WT/^*^Δ^) and *Atg5*^Δ*Cd207*^ mice. **(k)** Representative dot plot of Mitosox Red staining and comparison of Mitosox^high^ percentage of epidermal LCs obtained from control (*Atg5^WT^* and *Atg5^WT/^*^Δ^) and *Atg5*^Δ*Cd207*^ mice. All data are pooled from at least 3 independent experiments, with each point representing one individual 3-week-old mouse. Statistical analysis: Kruskal-Wallis one-way ANOVA followed by Dunn’s multiple comparison test (except g: two-way ANOVA followed by Šídák’s multiple comparisons test). (*, p<0.05; **, p<0.01; ***, p<0.001; ns, p>0.05).

Recent insights into different tissue macrophages showed that LCs heavily rely on mitochondrial oxidative phosphorylation^59^. To evaluate whether a lack of autophagy affects this pathway, we quantified the metabolic flux of LCs exposed to a series of inhibitors of mitochondrial respiratory complexes **(Figure 5f).** We included bone marrow-derived DCs (BMDCs) because their metabolism has been scrutinized in many previous investigations and relies mostly on beta-oxidation^60^. The decrease of oxygen consumption rates for wild-type and ATG5-deficient LCs upon oligomycin treatment was similar to that of BMDCs, indicating a similar basal production of ATP **(Figure 5g: ATP)**. Wild-type LCs and BMDCs also displayed a comparable profile after exposure to FCCP, which unleashes the maximal respiratory capacity of a cell **(Figure 5g: Max)**. On the other hand, LCs of *Atg5*^Δ*Cd207*^ mice reacted to FCCP by a strikingly prominent peak of their oxygen consumption. This implies that, in the absence of autophagy, the potential of LCs to mobilize oxidative phosphorylation, also called spare respiratory capacity, had massively increased **(Figure 5g: SRC)**. Despite this, ATG5-deficient LCs appeared unable to use this capacity to promote ATP production **(Figure 5g: ATP)**. The respiratory capacity of a cell relies on mitochondria, and transcriptome analysis demonstrated that mitochondria-related genes were differentially regulated upon loss of autophagy **(Figure 5h)**. Thus, we performed a double staining with mitotracker (MT) Green and Deep Red on LCs extracted from 3-week-old mice. While MT Deep Red is sensitive to mitochondrial membrane potential, MT Green stains mitochondrial membranes independently of the membrane potential, thus allowing normalization of the membrane potential to the mitochondrial load. An increased mitochondrial mass was detected **(Figure 5i)**, in line with the increased capacity of energy production that we measured by the mitochondrial stress assay. ATG5-deficient LCs did not display decreased membrane potential **(Figure 5j),** suggesting that mitochondrial function was preserved. Mitophagy is a key process to eliminate defective mitochondria, in particular when they produce reactive oxygen species (ROS). We used Mitosox staining to quantify ROS produced under altered mitochondria function, but this assay did not reveal any difference between *Atg5*^Δ*Cd207*^ and *Atg5^WT^* mice **(Figure 5k)**. All things considered, since mitochondria and lysosomes of autophagy-deficient LCs remained functional, we conclude that shortage in the lipophagy-dependent fatty acid supply could not be compensated by increased mitochondrial mass.

### Excess lipid oxidation causes ferroptosis in ATG5-deficient Langerhans cells

Mitosox detects superoxide O2·-, yet other ROS may cause cellular damage and death, i.e. H_2_O_2_ and HO·. Interestingly, ATG5-deficient LCs showed upregulated transcription of *Gss* (Glutathione-S Synthetase), *Slc7a11* (Cysteine/glutamate antiporter xCT), *Esd* (S-formylglutathione hydrolase) and *Gclm* (Glutamate-Cysteine Ligase Modifier Subunit), which are key elements in the glutathione-dependent response to such ROS **(Figure 6a)**. The glutathione pathway is notably involved in preventing ferroptosis, in which cell death occurs as a consequence of iron-dependent lipid peroxidation^61^. In favour of this hypothesis, we noticed several genes that showed significant moderate or high (more than two-fold) upregulation in ATG5-deficient LCs. They included ferroptosis-related genes such as *Hfe* (Homeostatic iron regulator), *Ftl1* (Ferritin Light Chain), *Sat1* (Spermidine/Spermine N1-Acetyltransferase 1), *Lpcat3* (Lysophosphatidylcholine Acyltransferase 3) and *Tfrc* (Transferrin receptor protein 1) **(Figure 6a)**. Accordingly, LCs of *Atg5*^Δ*Cd207*^ mice had increased surface expression of CD71/Transferrin receptor, which is required for iron import into the cells **(Figure 6b)**. Consistently, the lack of autophagy led LCs to accumulate ferrous iron **(Figure 6c),** which catalyses the production of oxidated lipid species. We also noticed elevated expression of *Acsl4* (acyl-CoA synthetase long-chain family member 4) and *Ptgs2* (Prostaglandin-Endoperoxide Synthase 2/Cyclooxygenase 2)^62^, which are considered as strong indicators of ongoing ferroptosis. Finally, in RNA expression analyses based on Kyoto Encyclopedia of Genes and Genomes (KEGG) pathways, 26 out of 137 genes (19%) in the apoptosis network were differentially expressed by autophagy-deficient LCs, whereas 39% of ferroptosis-related genes (16 out of 41) were affected **(Figure S4f, g; Table 3)**.

**Figure 6:**
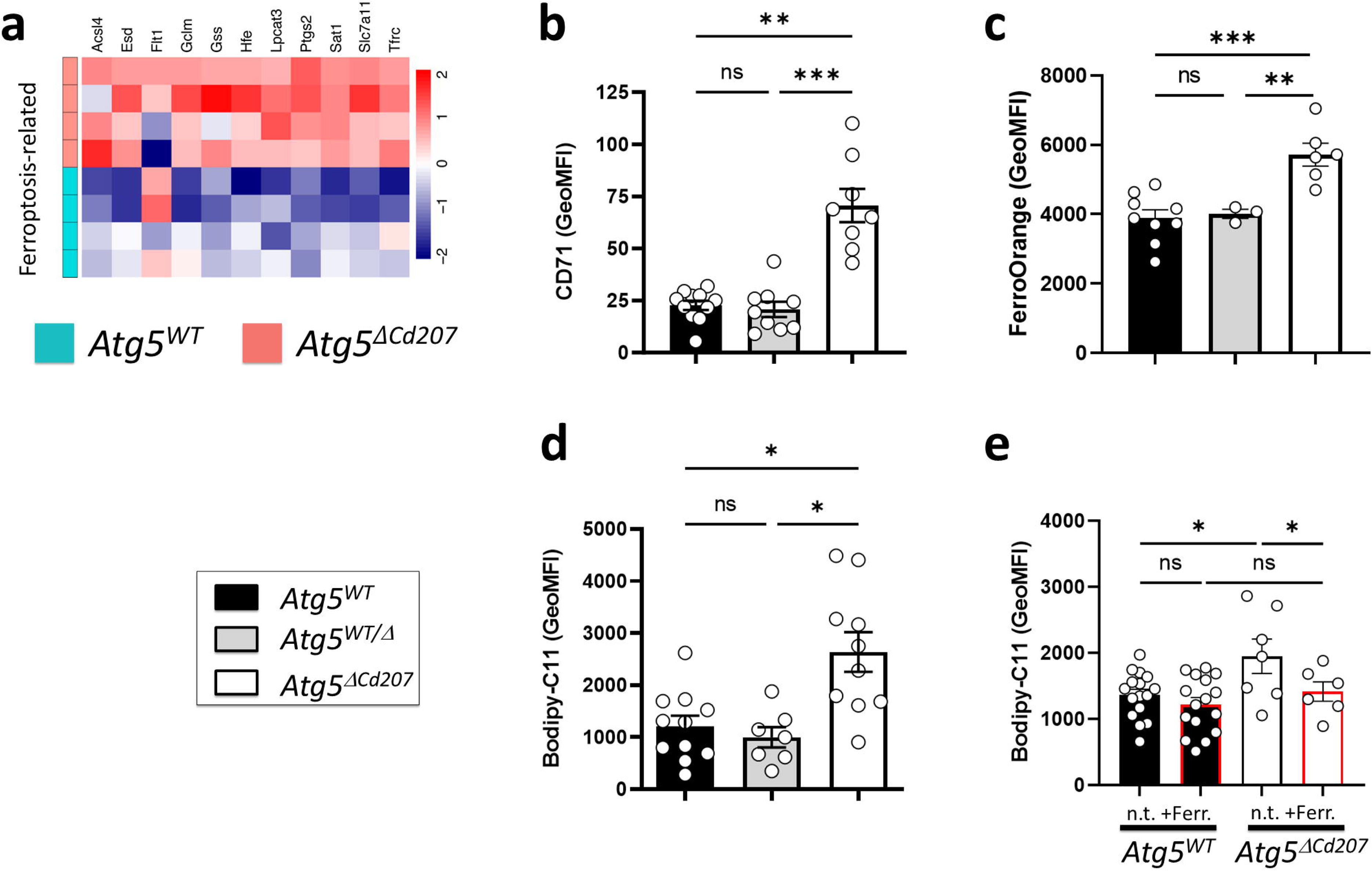
Autophagy-deficient Langerhans cells undergo ferroptosis. **(a)** Differentially expressed transcripts included in the KEGG ferroptosis pathway (mmu04216) in *Atg5^WT^* vs. *Atg5*^Δ*Cd207*^ LCs. No threshold was applied to fold changes**. (b)** MFI of CD71 at the surface of LCs. **(c)** MFI of FerroOrange staining, reflecting the concentration of ferrous iron (Fe2+) in LCs. Data are pooled from 5 independent experiments, with each point representing one individual 3-week-old mouse. **(d)** Quantification of lipid peroxidation for epidermal LCs of control (*Atg5^WT^* and *Atg5^WT/^*^Δ^) and *Atg5*^Δ*Cd207*^ mice, as measured by MFI of Bodipy-C11. Data are pooled from 10 independent experiments, with each point representing one individual 3-week-old mouse. **(e)** Lipid peroxidation of *Atg5^WT^* and *Atg5*^Δ*Cd207*^ LCs after overnight incubation with 50µM ferrostatin-1. Data are pooled from 4 independent experiments, with each point representing one individual 3-week-old mouse. Statistical analyses: (b-d) Kruskal-Wallis one-way ANOVA followed by Dunn’s multiple comparison test (***, p<0.001; **, p<0.01; *, p<0.05; ns, p>0.05). (e) two-way ANOVA followed by Sidak’s multiple comparison tests. (*, p<0.05; ns, p>0.05).

To confirm that lipid peroxidation occurs in the absence of functional autophagy, LCs were exposed to Bodipy-C11, a derivative of undecanoic acid that emits fluorescence upon oxidation. Upon treatment with this compound, LCs harvested from *Atg5*^Δ*Cd207*^ mice displayed significantly higher fluorescence intensity than LCs of *Atg5^WT^* mice **(Figure 6d)**. Finally, we exposed enriched LCs in vitro to the specific ferroptosis inhibitor ferrostatin-1^63^. Whereas Bodipy-C11 oxidation in *Atg5^WT^* LCs was unaffected, the ferrostatin-1 treatment normalized the elevated levels observed in *Atg5*^Δ*Cd207*^ LCs **(Figure 6e)**. Altogether, autophagy-deficient LCs concomitantly show an increase in lipid peroxidation, high expression of CD71, an accumulation of Fe^2+^ and sensitivity to ferrostatin-1, thereby supporting the notion that oxidation of the accumulated lipids leads them to ferroptosis, which contributes to their progressive depletion from the epidermis.

### Langerhans cells under metabolic stress have a proinflammatory signature

Besides metabolic imbalance, RNA sequencing revealed dysregulation of inflammation-related genes **(Figures S4e and 7a)**. In particular, and as confirmed by RT-qPCR **(Figure 7b)**, autophagy-deficient LCs increased their expression of mRNA encoding chemokines CXCL1, CXCL2 and CXCL3, known to attract neutrophils through CXCR2. Despite this observation, no obvious signs of inflammation were observed on the skin of *Atg5*^Δ*Cd207*^ mice: when analysing ear skin of 3-week-old mice for myeloid infiltrates **(Figure 7c)**, the proportions of Gr1^+^ Ly6G^+^ neutrophils or Gr1^low^ Ly6G^-^ monocytes did not differ between *Atg5^WT^* and *Atg5*^Δ*Cd207*^ mice in untreated conditions **(Figure 7d-g)**. Since *Nlrp3*, encoding a key inflammasome component, was markedly upregulated in LCs of *Atg5*^Δ*Cd207*^ mice, we challenged this pathway by injecting intradermally a small dose of alum hydroxide into the ears of 3-week-old mice. In *Atg5^WT^* mice, this resulted in only a modest increase of neutrophils **(Figure 7d)**, whereas monocytes were not attracted **(Figure 7e)**. On the other hand, alum hydroxide injection was able to drive a significant monocyte infiltration into the ears of *Atg5*^Δ*Cd207*^ mice **(Figure 7e**). Despite this, no difference could be demonstrated when comparing the extent of immune infiltrates of alum-treated *Atg5^WT^* and *Atg5*^Δ*Cd207*^ mice. As an alternative danger signal, we evaluated poly(I:C), which engages TLR3 and/or RIG-I signalling pathways. Poly(I:C) injection promoted recruitment of both neutrophils and monocytes in *Atg5^WT^* mice **(Figure 7f,g)**. Yet, inflammatory infiltrates were identical in *Atg5^WT^* and *Atg5*^Δ*Cd207*^ mice. These results suggest that autophagy-deficient LCs, although they exhibit a proinflammatory profile, do not prompt an exacerbated response to the challenges tested here.

**Figure 7:**
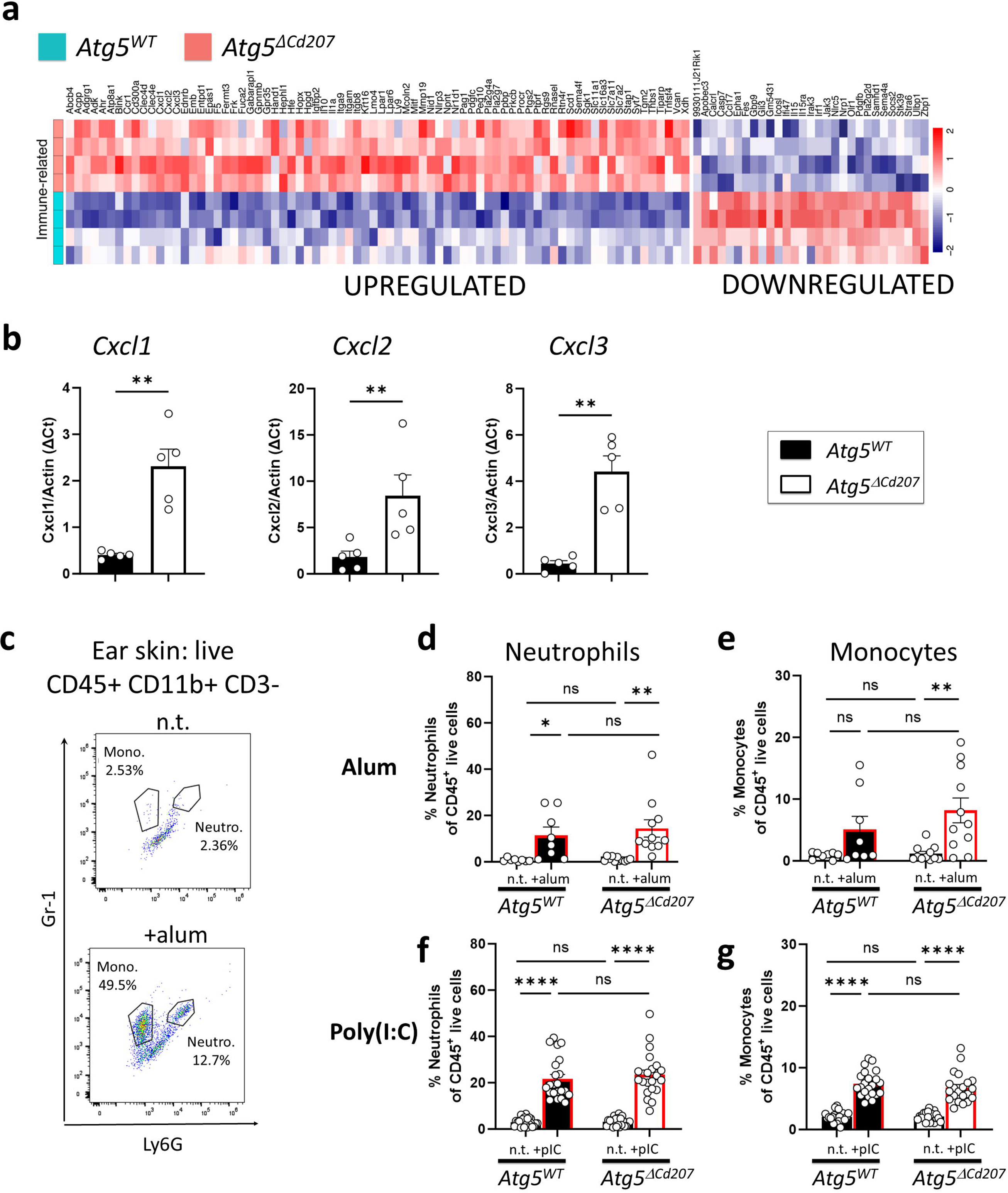
Langerhans cells under metabolic stress have a proinflammatory signature. **(a)** Differentially expressed transcripts related to immune function in *Atg5^WT^* vs. *Atg5*^Δ*Cd207*^ LCs (GO:0048871, GO:0009611, GO:0001660, GO:0035458, GO:0070672, GO:0019221, GO:0097529, GO:0006954, GO:0002274, GO:0050865, GO:1905517, GO:0001773). **(b)** Expression of *Cxcl1*, *Cxcl2* and *Cxcl3* genes quantified by RT-qPCR in purified LCs of the indicated genotypes. Data are pooled from 2 independent experiments, with each point representing one individual 3-week-old mouse. **(c-g)** One ear of *Atg5^WT^* or *Atg5*^Δ*Cd207*^ mice was injected intradermally with 2.5µg alum hydroxide **(c,d,e)** or 10µg poly(I:C) **(f,g)** and the contralateral ear was left untreated. 4h later, whole skin was digested and cell suspensions were monitored by flow cytometry for CD45+ CD3-CD11b+ Gr1+ Ly6G+ neutrophils and CD45+ CD3-CD11b+ Gr1 low Ly6G-monocytes (C: representative example). Percentages of neutrophils **(d,f)** and monocytes **(e,g)** were calculated as a proportion of live CD45+ cells. Data are pooled from 6-9 independent experiments, with each point representing one individual 3-week-old mouse. Statistical analysis: Mann-Whitney U test (b) or two-way ANOVA followed by Sidak’s multiple comparison tests. (*, p<0.05; **, p<0.01; ****, p<0.0001; ns, p>0.05).

### Autophagy deficiency affects neuronal interaction genes in Langerhans cells

Several reports showed that a long-term absence of LCs leads to a decrease of intraepidermal nerve endings, although the pathways governing this remain unidentified^64,65^. In our model, autophagy-deficient LCs downregulated a set of genes involved in neuronal interactions and axonal guidance (GO:0045664, GO:0050808, GO:0071526, GO:0030031; **Figures S4e and S6a)**. Thus, we sought to investigate the epidermal neuronal network of mice aged 6 months and older. Since neuronal development is particularly sensitive to autophagy impairment^66^, we chose to compare the density of LCs and epidermal sensory neurons of *Atg5*^Δ*Cd207*^ mice with that of *Atg5^WT^* and heterozygous *Atg5^WT/-^* mice **(Figure S6b)**. As we have demonstrated for mice of this age, there is a decrease in LC density in *Atg5*^Δ*Cd207*^ epidermis, but not in control mice **(Figure S6c)**. In parallel, quantification of β3-tubulin staining demonstrated that only *Atg5*^Δ*Cd207*^ mice presented significantly less epidermal nerve endings as compared to *Atg5^WT^* mice, although a slight reduction was observed in *Atg5^WT/-^* mice **(Figure S6d)**. Intriguingly, we found that LC numbers were correlated with the density of epidermal nerve endings and *Atg5^WT^* mice, but not in *Atg5^WT/-^*or *Atg5*^Δ*Cd207*^ mice **(Figure S6e).** This suggests that the autophagy-deficient LCs that remain in aging mice are unable to support neuronal epidermal growth, in line with their decreased neuron-related transcripts. Thus, in accordance with previous findings, the metabolic stress resulting from reduced autophagy in LCs may have an impact on the maintenance of the epidermal neuronal network.

## DISCUSSION

In contrast to conventional DCs, epidermal LCs self-renew to maintain their population and they are exposed in the steady state to environmental conditions (hypoxia, irradiation) that may favour autophagy. We report here a major role for autophagy in LC homeostasis. Indeed, constitutively autophagy-deficient LCs progressively disappeared from the epidermis. The depletion of autophagy-deficient LCs was not due to their emigration or decreased proliferation, suggesting cell death as the most likely explanation. Our analyses excluded a significant contribution of the ER stress response and major mitochondrial damages. In the absence of the key autophagy mediator ATG5, LCs displayed clear signs of lipid-related metabolic stress and underwent progressive depletion from the epidermis. They enhanced production of inflammation-related transcripts and showed less expression of innervation-regulating genes, which is of great interest in the context of a decreased network of epidermal nerves when these mice were aging.

The similar depletion of LCs observed using *Cd207*-specific *Atg7* deletion supports the idea that ATG5-dependent non-autophagic functions, are not directly involved in the maintenance of LCs. We could also rule out lysosomal alterations or the impairment for endocytic mechanisms such as LAP, since the LC network appeared unperturbed in *Rubcn*^-/-^ mice. The importance of autophagy for other skin APCs *in vivo* has been investigated previously. An earlier report did not find any consequence on the maintenance for CD207^+^ cDC1, in line with our findings^26^, although the absence of Vps34, which plays a role in autophagy and other biological pathways, resulted in the depletion of splenic cDC1^67^. Nevertheless, the ontogeny and features of LCs differ strikingly from those of cDC subsets^1^. Surprisingly, LCs had a continuous presence in skin-draining LNs, despite their depletion from the epidermis. This could be due to LCs remaining proliferative in older mice, which allowed the steady-state flux of LCs towards LNs to be kept constant. Emigration to LNs also entails a major environmental change as compared to the epidermis, and this may be beneficial to extended survival of LCs.

The metabolic requirements of LCs have rarely been studied and were mostly extrapolated from those of keratinocytes nearby. In the epidermal layer where they reside, LCs are relatively distant from dermal blood capillaries and have limited access to glucose and oxygen, suggesting that fatty acids may represent a major resource. This was recently substantiated by Wculek et al., who showed that disrupting mitochondrial function led to the partial depletion of LCs^59^. In line with this, we showed a remarkable accumulation of lipid droplets when autophagy was abrogated, in addition to a metabolic stress in freshly isolated LCs, revealed by AMPK phosphorylation. The seemingly unaltered ATP production could be due to normoxic in vitro culture conditions during Seahorse assays.

Interestingly, a hypoxic environment is expected to bias cellular metabolism towards an increase of lipid storage under the control of HIF-1α^68^. We postulate that lipophagy may be important for LCs to maintain a sustainable level of lipids, limit their potential cellular toxicity and provide free fatty acids. In this context, we propose that ferroptosis, resulting from uncontrolled supply of lipids and their peroxidation by ROS, represents one important mechanism explaining LC depletion in vivo. Ferroptosis was demonstrated by several key features of autophagy-deficient LCs: Bodipy-C11 assays, upregulation of CD71 and relevant detoxification pathways, and inhibition of lipid peroxidation by ferrostatin-1^61^. Another mechanism contributing to depletion could be apoptosis, also occurring in a proportion of ATG5-deficient LCs, most likely as a result of the critically low ATP:AMP ratio^69^ revealed by increased phosphorylation of AMPK.

Epidermal hypoxia appears to regulate the functional properties of human LCs^70^. Hypoxic tissues exhibit low levels of phosphorylated AKT^7^, which is expected to promote autophagy. LCs attempted to compensate autophagy impairment by inhibiting the PI3K/AKT pathway: they decreased expression of Inositol-3-phosphate synthase 1 (*Isyna1*) and of TNF-α-induced protein 8-like protein 3 (*Tnfaip8l3*), which shuttles PIP2 and PIP3 across the plasma membrane^71^, whereas Protein Kinase C beta (*Prkcb*) was upregulated. Although historically autophagy has been shown to be regulated through mTOR complex 1 (in particular under energetic stress), it is now clear that several induction pathways coexist with constitutive activity, especially in immune cells. Intriguingly, progressive disappearance of epidermal LCs at a young age has been reported in mice with selective disruption of critical intermediates of the mTOR pathway^72–74^. Therefore, since mTOR is recognized as a negative regulator of autophagy, it could be interpreted that excessive autophagy is detrimental to LCs. However, autophagy has not been investigated in these mouse models. In addition, mTOR regulates many other cellular processes and, beyond autophagy, is critical to survival, lysosomal trafficking or cytokine signalling pathways. By impairing the recycling of receptors through lysosomes, LAMTOR2 deletion leads to a defect in TGF-β signalling, which is essential for the differentiation of LCs^75^. Moreover, LCs deficient in Raptor, an adaptor of the mTOR complex 1, leave the epidermis, which may also result from an impaired TGF-β signalling that normally maintains the epidermal LC network by restricting their spontaneous migration to skin-draining LNs^48,49^. Altogether, there is no direct evidence to date that the deleterious impact of genetic ablations affecting the mTOR pathway in LCs may depend solely on altered autophagy. Finally, the Nrf2/Keap1 pathway plays an important role in the electrophilic and oxidative stress response of cutaneous antigen-presenting cells, including LCs^76^. In this context, it is worth noting that Keap1, which prevents the activity of transcription factor Nrf2, can be degraded through autophagy, and that Nrf2 target genes *Sqstm1* and *Hmox1* show increased expression in LCs of *Atg5*^Δ*Cd207*^ mice.

ATG5-deficient LCs displayed an accumulation of lipid droplets that likely resulted from impaired lipophagy. Consistent with the data reported here, a similar phenotype of lipid accumulation has been described in murine LCs in a model of imiquimod-induced psoriasis, and, strikingly, the authors also found signs of dysregulated autophagy for LCs in this model and in psoriatic patients^77^. In our selective autophagy impairment model, LCs did not promote glycolysis and were unable to take advantage of their increased respiratory capacity linked to higher mitochondrial mass, highlighting the critical importance of lipophagy for their energy production and limitation of potentially toxic accumulation of lipids. The limited ER swelling observed in our model, which was not sufficient to trigger the UPR pathway in LCs, may be related to defective energy mobilisation from lipid storage^50,78^, multiple budding of lipid droplets^55^ or modification of ER membrane dynamics following lipid peroxidation^79^. Accumulation of lipid droplets as a result of an impaired autophagy machinery has been observed in other cell types that rely on lipophagy. This catabolic process is key in the development of neutrophils^80^. Of note, DCs derived from *Atg5^-/-^*fetal liver display elevated CD36 expression and lipid droplets, although no cell death was reported during the time frame of this *in vitro* experiment^38^. Yet, the lifespan of bone marrow-derived or conventional DCs is not comparable to that of LCs, in which alterations in lipid metabolism may have deleterious consequences when they accumulate over long periods of time. Considering that CD207 is expressed in LCs about 10 days after birth^43^, the time period at which LC depletion becomes visible (around 20 days of age) suggests that they cannot survive more than 2 weeks to autophagy deficiency in the steady state. This represents a relatively short delay as compared to other, unrelated cell types previously found to rely on autophagy, i.e. B-1 B cells that survive 6 weeks after deletion of *Atg5*^19^.

Some features of LCs are reminiscent of tissue-resident macrophages^1^. Although autophagy regulates many functions of macrophages, it was not considered to play a prominent role in their homeostatic maintenance^39^. Absence of autophagy affects the survival of a subset of peritoneal macrophages^40^, which are of embryonic rather than monocytic origin, similar to steady-state LCs. It remains to be demonstrated whether the dependence on autophagy can be associated with the origin and/or long-term residency for macrophages within other organs.

The consequences for the epidermis of a long-term absence of LCs have been investigated through constitutive diphtheria toxin-mediated depletion in the huLangerin-DTA mouse strain^64,81^. However, since LCs are absent at birth, it is difficult to identify which of the genes that they normally express may affect epidermal homeostasis. Here, we were able to document transcriptome alterations of ATG5-deficient LCs that are still present in young mice, albeit in a metabolically stressed state. First and foremost, our data suggested a potential for supporting inflammation. Yet, cutaneous immune infiltration did not occur spontaneously, and inflammatory challenges by alum hydroxide or poly(I:C) did not result in a larger immune infiltration in mice where LCs were impaired for autophagy. This may be explained by the fact that autophagy-deficient LCs progressively disappear, limiting their capacity to induce inflammation. Secondly, several genes involved in neuronal interactions were downregulated by ATG5-deficient LCs, including EPH receptor A1, Semaphorin 4A, Neuropilin-1 and Neuregulin-1. The long-term absence of LCs in our model may lead to a decrease of epidermal nerve endings. These findings thus represent a milestone for future investigations on neuroimmune interactions, considering the putative role of LCs and dermal macrophages^82^ in regulating sensory neuron growth and repair in the skin.

Altogether, we shed light on the metabolic adaptations of LCs that ensure their long-lasting tissue residency. It will be of great interest to translate these findings in the context of human skin diseases, considering that lipid supply and autophagy capacity of LCs may perturb their homeostasis and favour inflammation.

## MATERIAL AND METHODS

### Mice

Mice were bred and maintained on a C57BL/6J background at the animal facility of the Institut de Biologie Moléculaire et Cellulaire (IBMC). *Atg5^flox/flox^*and *Cd207-cre* mice were gifted by N. Mizushima and B.E. Clausen, respectively^66,83^. *Atg5^+/-^* mice were generated at the IBMC^18^. [*Atg5^+/-^*; *Cd207-cre*] were obtained from a first cross between *Cd207-cre* and *Atg5^+/-^*, then bred to *Atg5^flox/flox^* to obtain [*Atg5^flox/-^*; *Cd207-cre*] (*Atg5*^Δ*Cd207*^) and littermates [*Atg5^flox/+^*; *Cd207-cre*] (*Atg5^WT/^*^Δ^) and [*Atg5^flox/+^*] (*Atg5^WT^*). Mice were genotyped for their *Atg5* allele and the *Cd207-cre* transgene as previously described^18,83^. All mice were bred and maintained in accordance with guidelines of the local institutional Animal Care and Use Committee (CREMEAS).

### Antibodies and reagents for flow cytometry and immunofluorescence microscopy

Antibody stainings for flow cytometry or immunofluorescent microscopy were performed in SE buffer (fetal calf serum 2%, EDTA 2.5mM). All reagents and antibodies are listed in **Table 1**.

### Cell preparation and culture

#### Lymph nodes

Brachial and inguinal lymph nodes were digested for 1h at 37°C under shaking in R2 buffer (RPMI-1640 medium containing L-glutamine (Lonza) plus 2% fetal calf serum (Dutscher)) supplemented with 50μg/mL DNAse and 10μg/mL collagenase D (Roche).

#### Digestion of back skin epidermis (electron microscopy, LC proportions, caspase-3 activation, proliferation assays)

Back skin was incubated with 0.25% Trypsin (VWR) for 45min at 37°C. After removal of the dermis, the epidermis was teased apart with forceps, followed by 15min of gentle shaking on a rotating wheel. Where indicated, CD11b^+^ LCs were enriched by magnetic bead separation (Miltenyi-Biotec).

#### Ear skin digestion (skin DC subsets, quantification of immune infiltrates)

Ear skin was cut into small pieces, digested in R2 buffer containing 0.15mg/ml LiberaseTM and 0.12mg/ml DNAse (Roche) for 45min at 37°C and filtered through 100µm cell strainers.

#### Epidermal crawl-out assay (Bodipy C16 and glucose uptake, in vitro treatment with inhibitors, Seahorse assay)

Back skin was incubated overnight at 4°C in R2 buffer, containing 1mg/mL dispase II (Roche). The separated epidermis was then laid upon cell culture medium (RPMI-1640 medium supplemented with 10% fetal calf serum, 50mM β-Mercaptoethanol (Gibco), 1% Gentamicin (Gibco), and 10mM HEPES (Lonza) in a Petri dish for 24h at 37°C, allowing emigration of LCs.

#### Bone marrow-derived DCs

Femurs and tibias were collected from C57BL/6J mice. Bone marrow was flushed out, red blood cells lysed, filtered and cultured for 7 days in complete RPMI medium (RPMI-1640 medium containing L-glutamine plus 10% fetal calf serum) containing 20ng/mL recombinant GM-CSF (Peprotech).

### Induction of cutaneous inflammation

For each mouse, one ear was injected intradermally with 25µL of 100µg/mL alum hydroxide (Roche), and the contralateral ear was left untreated. 4h later, whole skin was digested and cell suspensions were monitored by flow cytometry for CD45^+^ CD3^-^ CD11b^+^ Gr1^+^ Ly6G^+^ neutrophils and CD45^+^ CD3^-^ CD11b^+^ Gr1^+^ Ly6G^-^ monocytes.

### Electron microscopy

Epidermal cell suspensions (freshly isolated or cultured for 3 days) were processed for electron microscopy essentially as described^84^. Briefly, after pre-enrichment on bovine serum albumin density gradient, cells were washed and fixed using Karnovsky’s formaldehyde-glutaraldehyde fixative for 1h at room temperature. Specimens were post-fixed in aqueous 3% osmium tetroxide and contrasted with 0.5% veronal-buffered uranyl acetate. Dehydration of samples was done in graded series of ethanol concentrations, followed by embedding in Epon 812 resin. Ultrathin sections were mounted on nickel grids, contrasted with lead citrate and examined by transmission electron microscopy (Phillips EM 400; Fei Company Electron Optics, Eindhoven, The Netherlands) at an operating voltage of 80kV. LCs were identified within epidermal cell suspensions by their electron-lucent cytoplasm, the absence of keratin tonofilament bundles, the presence of cytoplasmic processes (dendrites) and their ultrastructural hallmarks, the Birbeck granules.

### Autophagy flux assessment by flow cytometry

Measurements of autophagy fluxes were carried out using the Guava Autophagy LC3 Antibody-based Detection Kit (Luminex). LCs isolated through epidermal crawl-outs were cultured 18h at 37°C with or without the lysosome inhibitor provided with the kit (60 µM hydroxychloroquine). After labelling by FVD450, cells were stained for CD45, I-A/I-E, and TCRγ/δ. Cells were permeabilized with 0.05% saponin (Merck Millipore) to wash out the cytosolic LC3-I, then membrane-associated LC3 (LC3-II) was preferentially stained with anti-LC3 FITC (clone 4E12). Flow cytometry analysis allowed to calculate autophagy fluxes, dividing the LC3-FITC mean fluorescence intensities (MFI) of treated cells by that of untreated cells.

### Glucose uptake

Cells obtained by crawl-out were glucose-starved for 24h in PBS (Lonza) supplemented with 0.5% fetal calf serum for 8 hours. Cells were then incubated for 30min at 37°C with 150µM of 2-[N-(7-nitrobenz-2-oxa-1,3-diazol-4-yl) amino]-2-deoxy-D-glucose (ThermoFisher).

### Pharmacological inhibitions

Cells obtained by crawl-out were incubated for 24h at 37°C with the phosphatidylinositol-3-kinase inhibitor wortmannin or the carnitine palmitoyltransferase-1 inhibitor etomoxir (both from Sigma-Aldrich), at 10 and 200μM respectively.

### 5-bromo-2’-deoxyuridine incorporation

1mg of 5-bromo-2’-deoxyuridine (BrdU; Sigma) was administered by intraperitoneal injection 72 hours prior to analysis. Drinking water also contained 0.8mg/mL BrdU. Following staining of surface markers CD45, I-A/I-E and TCRγδ, epidermal single-cell suspensions were fixed with Cytofix/Cytoperm buffer (BD Biosciences) and permeabilized with permeabilization buffer (BD Biosciences). DNA was then denatured with a DNAse solution (100µg/mL, BD Biosciences) to improve the accessibility of the incorporated BrdU to the detection antibody.

### Quantitative real-time RT-PCR analysis

RNA was extracted from cells sorted from lymph nodes or epidermis on a FACS Melody cell sorter (BD Biosciences) with RNeasy microKit (Qiagen) for lymph nodes and Trizol for epidermis (ThermoFisher). cDNA was obtained with Maxima Reverse Transcriptase Kit (ThermoFisher) using a T100 Thermal cycler (Biorad) for *Atg5* quantification and with RevertAid H Minus First Strand cDNA Synthesis Kit (ThermoFisher) for *Cxcl1*, *Cxcl2* and *Cxcl3* quantification. Quantitative real-time PCR was performed on cDNA using Taqman preAmp MasterMix and Taqman Universal Mastermix (ThermoFisher) and Assays-on-Demand probes (*Gapdh*: Mm03302249_g1; *Atg5: Mm00504340_m1*) for *Atg5* quantification and Fast SYBR Green Master Mix (ThermoFisher) with primers Actb (β-Actin) forward: 5’-CATTGCTGACAGGATGCAGAAGG-3’; β-Actin reverse: 5’-TGCTGGAAGGTGGACAGTGAGG-3’; Cxcl1 forward: 5’-TCCAGAGCTTGAAGGTGTTGCC-3’; Cxcl1 reverse: 5’-AACCAAGGGAGCTTCAGGGTCA-3’; Cxcl2 forward: 5’-CATCCAGAGCTTGAGTGTGACG-3’; *Cxcl2* reverse:5’-GGCTTCAGGGTCAAGGCAAACT-3’; *Cxcl3* forward: 5’-TGAGACCATCCAGAGCTTGACG-3’; *Cxcl3* reverse: 5’-CCTTGGGGGTTGAGGCAAACTT-3’ (Eurogentec) for the quantification of *Cxcl1*, *Cxcl2* and *Cxcl3*. Each sample was amplified in triplicate or duplicate in a StepOnePlus real-time PCR system (Applied Biosystems). mRNA levels were calculated with the StepOne v2.1 software (Applied Biosystems), using the comparative cycle threshold method, and normalized to the mean expression of *Gapdh* (for *Atg5*) and *Actb* (for *Cxcl1*, *Cxcl2* and *Cxcl3*) housekeeping genes.

### Immunofluorescence microscopy of epidermal sheets

Ear epidermis was separated from the dermis by ammonium thiocyanate digestion (0.15M) for 20min at 37°C. Alternatively, for optimal preservation of neuronal networks, epidermal sheets were separated after 10mM EDTA diluted in PBS for 1h. Epidermis was then fixed by incubation in PBS 4% PFA or in glacial acetone for 15min at 4°C followed by incubation with PBS 5% BSA 0.1% Triton. Primary antibodies were incubated overnight at 4°C. After fixation, epidermal sheets were washed four times in blocking buffer consisting in 5% BSA in PBS for 15 minutes each time at room temperature. Sheets were then incubated overnight at 4°C with the primary antibodies: anti-β3-tubulin and AF647 anti-CD207 diluted in blocking buffer. After washing the sheets as described above, they were incubated with a solution of goat anti-mouse AF594, and 4’,6-diamidino-2-phenylindole (DAPI) in blocking buffer for 1h at room temperature. After additional washings, epidermal sheets were mounted in Fluoromount-G mounting medium (ThermoFisher) and observed under a confocal microscope (Yokogawa Spinning Disk, Zeiss). Whole-mount epidermal images were processed using the open-source software FIJI to measure the total analysed area for each sample and to quantify the mean fluorescence intensity.

### Immunofluorescence microscopy of epidermal cell suspensions

Cell suspensions were deposited on Lab-Tek chamber slides (Thermo Scientific Nunc) previously coated with a poly-L-Lysine solution (Sigma-Aldrich) diluted in ultra-pure water at 0.02% (v/v) to enhance cellular adhesion. Epidermal cells were then incubated with Mitotracker, Mitosox, ER-tracker, Bodipy or Bodipy-C16 according to the manufacturer’s instructions (Invitrogen), before fixation using 2% PFA in PBS for 15min at RT. DAPI was incubated 15min at RT. Tissues were mounted and observed under a confocal microscope (Yokogawa Spinning Disk, Zeiss).

### RNA sequencing

Total RNA was isolated from 10^5^ sorted LCs with the RNeasy Mini Kit (Qiagen). RNA integrity was evaluated on an Agilent Bioanalyzer 2100 (Agilent Technologies). Total RNA Sequencing libraries were prepared with SMARTer® Stranded Total RNA-Seq Kit v2 - Pico Input Mammalian (TaKaRa) according to the manufacturer’s protocol. Briefly, random priming was used for first strand synthesis and ribosomal cDNA has been cleaved by ZapR v2 in the presence of mammalian R-probes V2. Libraries were pooled and sequenced (paired-end 2*75bp) on a NextSeq500 using the NextSeq 500/550 High Output Kit v2 according to the manufacturer’s instructions (Illumina). For analysis, quality control of each sample was carried out and assessed with the NGS Core Tools FastQC (http://www.bioinformatics.babraham.ac.uk/projects/fastqc/). Sequence reads were mapped on the GRCm38 reference genome using STAR^85^ and unmapped reads were remapped with Bowtie2^86^ using a very sensitive local option to optimize the alignment. The total mapped reads were finally available in BAM (Binary Alignment Map) format for raw read counts extraction. Read counts were found by the HTseq-count tool of the Python package HTSeq^87^ with default parameters to generate an abundance matrix. Finally, differential analyses were performed using the DEseq2^88^ package of the Bioconductor framework. Differentially expressed genes between *Atg5*^Δ*Cd207*^ and *Atg5^WT^* were selected based on the combination of adjusted p-value < 0.05 and FDR < 0.1, with fold changes < -2 or > 2, unless otherwise stated in figure legends. Pathway enrichment analysis was performed using Metascape (https://metascape.org)^89^.

Finally, to highlight the genes that are part of the ferroptosis or the apoptosis pathways, we screened the differentially expressed genes (p<0.05) for their presence in the entries of the respective KEGG pathways using Excel **(Table 3)**, with https://www.genome.jp/pathway/mmu04216 for ferroptosis and https://www.genome.jp/entry/mmu04210 for apoptosis. The genes that overlapped with the pathway genes were then labeled using the visualization tool of KEGG.

### Metabolic parameter quantitation by extracellular flux assay

CD45^+^ MHCII^+^ CD207^+^ CD103^-^ TCRDD^-^ LCs were sorted from epidermal crawl-out suspensions on a FACSFusion cell sorter (Becton-Dickinson). Purified LCs or BMDCs (2.10^5^ cells/well) were seeded in Seahorse XF96 culture plate coated with poly-lysine (Sigma). After overnight culture, a Mitochondrial Stress Test was performed. In this assay, culture wells are injected sequentially with different inhibitors of the mitochondrial respiration. Energy production resulting from mitochondrial respiration was determined after each injection by measuring oxygen consumption rates (OCR, pmoles/min) on a Seahorse XF96 according to the manufacturer’s instructions (Agilent). Oligomycin (OM) injection allowed calculating the oxygen consumption used for mitochondrial ATP synthesis. Carbonyl cyanide 4-(trifluoromethoxy)phenylhydrazone (FCCP) uncoupled mitochondrial respiration, allowing to calculate maximal respiration and spare respiratory capacity. Finally, rotenone (ROT) and antimycin A (AA) blocked mitochondrial complex I and III to determine the non-mitochondrial oxygen consumption. The following metabolic parameters were calculated:

ATP production = OCR_baseline_ – OCR_OM_

Maximum respiration = OCR_FCCP_ – OCR_AA+ROT_

Spare respiratory capacity (SRC) = OCR_FCCP_ – OCR_baseline_

### Lipid peroxidation assay

50 000 enriched CD11b+ LCs were seeded and incubated for 10min at 37°C with 2mM Bodipy-C11 (581/591) (4,4-difluoro-5,7-dimethyl-4-bora-3a,4a-diaza-s-indacene-3-undecanoic acid; Invitrogen) in PBS. Cells were then resuspended in SE buffer and incubated with the following antibodies: CD3e-PerCP-Cy5.5, MHC-II-AF700 and CD45-APC-Cy7. Upon gating on CD45^+^ CD3-MHCII^+^ cells, the fluorescence of Bodipy-C11 was collected from the FITC channel on a Gallios cytometer (Beckman-Coulter).

### In vitro ferroptosis inhibition

At least 50 000 enriched CD11b+ LCs were seeded and incubated overnight in complete RPMI medium and 50µM ferrostatin-1 or DMSO (untreated control). Cells were then stained and analyzed as indicated for the lipid peroxidation assay.

### Detection of intracellular Fe^2+^ with FerroOrange

To detect intracellular iron levels, 100 000 enriched CD11b+ LCs were incubated with 1µM FerroOrange (Sigma-Aldrich) in PBS for 30min at 37°C. After incubation, the cells were washed in PBS and stained for flow cytometry analysis. Upon gating on CD45^+^ CD3-MHCII^+^ cells, the fluorescence of FerroOrange was collected from the PE channel on a Gallios cytometer (Beckman-Coulter).

### Quantification of the density of epidermal nerve endings

The open-source software iLastik was used to segmentate images of whole-mount epidermal sheets stained for β3-tubulin and CD207, using machine learning to differentiate background from β3-tubulin signal. Images were then processed using the open-source software FIJI to measure the total area of each scan, as well as the area that was determined positive for β3-tubulin.

### Statistical analyses

Statistical significance was calculated with the indicated tests using Prism software (GraphPad, versions 6-9). All data were presented as mean ± standard error of the mean (SEM). P-values < 0.05 were considered statistically significant (*, p<0.05, **, p<0.01, ***, p<0.001, ****,p<0.0001).

## Supporting information

Graphical abstract

## ABBREVIATIONS

2-NBDG: 2-(N-(7-Nitrobenz-2-oxa-1,3-diazol-4-yl) Amino)-2-Deoxyglucose
AMPK: AMP-activated protein kinase
APCs: Antigen-presenting cells
*Atg*: Autophagy-related genes
DCs: Dendritic cells
cDCs: Conventional dendritic cells
DETCs: Dendritic epidermal T cells
ER: Endoplasmic Reticulum
FCS: Fetal calf serum
IRE1: Inositol-requiring enzyme 1
LCs: Langerhans cells
LN: Lymph nodes
MFI: Mean fluorescence intensity
MHC-II: Major histocompatibility complex
MT: Mitotracker
ROS: Reactive oxygen species
UPR: Unfolded protein response
TGF-β1: Tumor growth factor β 1
Xbp1: X-box binding protein 1

## AKNOWLEDGEMENTS

We thank Pr. Noboru Mizushima for his gift of *Atg5^flox/flox^* mice, Delphine Lamon and Fabien Lhericel for mouse breeding at the IBMC animal facility, as well as Claudine Ebel and Muriel Philipps at the cell sorting facility of the IGBMC.

## AUTHOR CONTRIBUTIONS

F.A, R.S.-C., F.G. and V.F. designed the research.

N.R. contributed electron microscopy images.

B.E.C., E.L.G. and T.H. contributed essential reagents and critically reviewed the paper

F.A, R.S.-C., D.B., W.B., Q.F., L.F., R.F., F.G. and V.F. performed the experiments.

O.G. and H.P. helped design and analyze the metabolic measurement assays

R.C., A.M., S.B. and B.V. designed and analyzed the RNA sequencing assay

F.A, W.B., R.S.-C, J.-D.F., F.G. and V.F. analyzed the data.

F.A, C.G.M., F.G. and V.F. wrote the paper.

## Supplementary figures

**Figure S1: Autophagosomes are detectable in murine Langerhans cells. (a)** Transmission electron microscopy of LCs in a bulk epidermal cell suspension from C57BL/6 mice, either freshly isolated (right panel) or cultured for 3 days (left and center panels). The inset image in the right panel highlights a Birbeck granule (arrow). **(b)** Close-up micrographs of autophagic structures corresponding to the boxes in the low power overview micrographs. 1 and 3 appear to be limiting membranes of incipient autophagy; 2 and 4 show double membrane-limited autophagosomes. Scale bars: 1µm (a); 500nm (b).

**Figure S2: Efficient deletion of *Atg5* in CD207+ DC subsets affects Langerhans cells but not cDC1. (a)** Representative electrophoresis of genotyping PCR. Left panel: floxed, wild-type (WT) and exon 3-deleted (KO) alleles of *Atg5*. Right panel: wild-type and *Cre* knock-in alleles of *Cd207*. **(b)** *Atg5* mRNA expression in sorted epidermal CD45+ MHCII+ TCRγδ^-^ LCs from control (*Atg5^WT^*and *Atg5^WT/^*^Δ^) and *Atg5*^Δ*Cd207*^ mice. Fold changes were calculated relative to mRNA expression in LCs of *Atg5^WT^*control mice. Data are pooled from at least 3 independent experiments, with each point corresponding to one individual mouse. Statistical analysis: Kruskal-Wallis one-way ANOVA followed by Dunn’s multiple comparison test (*, p<0.05; ns, p>0.05). **(c)** Gating strategy used to sort lymph nodes MHC-II^+^ CD207-FSA high dermal DCs (dDCs), MHC-II^+^ CD207^+^ CD103^-^ LCs (LCs) and MHC-II^+^ CD207^+^ CD103^+^ cDC1 (CD103^+^). Red dots in the top panel depict backgating of CD207^+^ LCs/cDC1. **(d)** *Atg5* mRNA expression in LCs, cDC1 and CD207-dDCs sorted from pooled lymph node cell suspensions of at least 3 control (*Atg5^WT^* and *Atg5^WT/^*^Δ^) or *Atg5*^Δ*Cd207*^ mice. Fold changes were calculated relative to mRNA expression in cells of *Atg5^WT^* control mice. ND, not detectable. **(e)** Representative dot plots for the identification of CD207^+^ CD11b^+^ LCs, CD207^+^ CD11b^-^ cDC1, CD207^-^ CD11b^+^ cDC2/macrophages and CD207^-^ CD11b^-^ (DN, double-negative) DCs among live CD45^+^ lineage-CD11c^+^ MHCII^+^ skin DCs in whole skin cell suspensions from *Atg5^WT^* and *Atg5*^Δ*Cd207*^ mice (lineage markers: B220, NK1.1, Ly6G and CD3). **(f)** Percentages of LCs and cDC1 among skin DCs. Data are pooled from at least 3 independent experiments, with each point corresponding to one individual mouse. Statistical analysis: Kruskal-Wallis one-way ANOVA followed by Dunn’s multiple comparison test (*, p<0.05; ns, p>0.05).

**Figure S3: ATG5-deficient Langerhans cells have functional lysosomes.** Representative half-set overlay of LysoTracker-Red (left panel) and LysoSensor (right panel) stainings and comparison of the ratio of MFI of each marker for epidermal LCs of control (*Atg5^WT^* and *Atg5^WT/^*^Δ^) and *Atg5*^Δ*Cd207*^ mice. Data are pooled from at least 3 independent experiments, with each point corresponding to one individual mouse. Statistical analysis: Kruskal-Wallis one-way ANOVA followed by Dunn’s multiple comparison test (ns, p>0.05).

**Figure S4: Lack of autophagy alters the transcriptome of Langerhans cells. (a)** Visualization of the exon 3 region of *Atg5* gene from RNA-seq of sorted LCs of indicated mouse genotype using integrative Genomic Viewer tool. **(b)** Principal component analysis of RNA-seq transcriptome analysis from sorted LCs of *Atg5^WT^* and A*tg5*^Δ*Cd207*^ mice. **(c)** Heatmap showing the differentially expressed genes (FDR<0.1, Absolute Log2 Fold Change value > 1, p-value<0.05) between LCs of indicated mouse genotypes **(d)** Volcano plot showing the differential expression of genes between LCs of indicated mouse genotypes. Gene names refer to the top 20 up and downregulated genes, based on the following combinations of p-value and fold-change criteria: blue dots: p-value<0.05 with no cutoff on Absolute Log2 Fold Change; red dots: p-value<0.05 and Absolute Log2 Fold Change value > 1. **(e)** Metascape pathway analysis of genes significantly upregulated or downregulated in *Atg5*^Δ*Cd207*^ LCs. **(f,g)** Differentially expressed genes related to **(f)** apoptosis and **(g)** ferroptosis KEGG pathways. Blue boxes: downregulated in *Atg5*^Δ*Cd207*^ LCs, red boxes: upregulated in *Atg5*^Δ*Cd207*^ LCs; dashed boxes: FDR>0.1.

**Figure S5: Inhibition of mitochondrial metabolism or autophagy leads to neutral lipid accumulation in epidermal Langerhans cells.** Flow cytometry quantification of the Bodipy MFI in epidermal LCs obtained from C57Bl/6 mice then treated with **(a)** etomoxir or **(b)** wortmannin. Data are pooled from at least 3 independent experiments, with each point corresponding to one individual mouse. Statistical analysis: Kruskal-Wallis one-way ANOVA followed by Dunn’s multiple comparison test (*, p<0.05; **, p<0.01).

**Figure S6: Depletion of autophagy-deficient Langerhans cells alters the epidermal neuronal network. (a)** Differentially expressed genes in *Atg5^WT^* vs. *Atg5*^Δ*Cd207*^ LCs of 3-week old mice: transcripts related to neuronal interactions (GO:0030031, GO:0071526, GO:0050808, GO:0045664). **(b)** Representative immunofluorescence microscopy image of epidermal sheets obtained from ears of 6–12-month-old *Atg5^WT^* and *Atg5*^Δ*Cd207*^ mice and stained with antibodies against β3-tubulin (neurons) and CD207 (LCs). Scale bar: 50µm. Quantification in *Atg5^WT^*, *Atg5^WT/-^* and *Atg5*^Δ*Cd207*^ mice: **(c)** Number of CD207^+^ LCs per mm². **(d)** Relative area of β3-tubulin staining. Data are pooled from 3 independent experiments, with each point corresponding to one field of view (n=3 mice per condition, at least 10 fields per mouse were scanned). Statistical analysis: Kruskal-Wallis one-way ANOVA followed by Dunn’s multiple comparison test (**, p<0.01; ****, p<0.0001; ns, p>0.05). **(e)** Correlation of epidermal nerve and Langerhans cell densities in 6–12-month-old *Atg5^WT^*, *Atg5^WT/-^* and *Atg5*^Δ*Cd207*^ mice. Based on epidermal sheet stainings (c, d), the number of CD207^+^ LCs per mm² (X axis) was plotted against the relative areas for β3-tubulin^+^ nerves (Y axis). Statistical analysis: Pearson correlation test. R² correlation coefficients and p-values are indicated.

**Supplementary Video SV1: Autophagosome staining of *Atg5^WT^* Langerhans cells.** LC3: green; CD207: red.

**Supplementary Video SV2: Autophagosome staining of *Atg5*^Δ*Cd207*^ Langerhans cells.** LC3: green; CD207: red.

**Supplementary Video SV3: Endoplasmic reticulum staining of *Atg5^WT^* Langerhans cells.** CD207: green; ER-tracker: red.

**Supplementary Video SV4: Endoplasmic reticulum staining of *Atg5*^Δ*Cd207*^ Langerhans cells.** CD207: green; ER-tracker: red.

**Supplementary Video SV5: Lipid droplets of *Atg5^WT^* Langerhans cells.** CD207: green; Bodipy: red.

**Supplementary Video SV6: Lipid droplets of *Atg5*^Δ*Cd207*^ Langerhans cells.** CD207: green; Bodipy: red.

**Table 1. Antibodies and fluorescent reagents.** This table identifies the exact references for all reagents used in the study.

**Table 2. Differentially expressed genes between Langerhans cells of *Atg5^WT^* and *Atg5***^Δ***Cd207***^ **mice.** The listed genes present with a significant difference (false discovery rate <0.1; p<0.05) with fold changes >0 (increased in *Atg5*^Δ*Cd207*^) or <0 (decreased in *Atg5*^Δ*Cd207*^).

**Table 3. Analysis of the ferroptosis and apoptosis KEGG pathways for dysregulated genes in Langerhans cells of *Atg5^WT^* vs. *Atg5*^Δ*Cd207*^ mice.** Differentially expressed genes in *Atg5*^Δ*Cd207*^ (Fold change>0; p<0.05; FDR as indicated) were screened among entries of the respective KEGG pathways (mmu04216 for ferroptosis; mmu04210 for apoptosis). The proportion of differentially expressed genes out of all genes involved in the pathway was calculated.

