## Supplementary Table 1 for "Epidermal maintenance of Langerhans cells relies on autophagy-regulated lipid metabolism"

| Antibody target or Reagent | Fluorochrome | Clone | Supplier | Reference |
|---|---|---|---|---|
| Perilipin-2/ADFP | Uncoupled | EPR3713 | Abcam | ab108323 |
| phospho-AMPKα (T183/T172) | Uncoupled | Polyclonal | Abcam | ab23875 |
| SQSTM1 / p62 | Uncoupled | 2C11 | Abcam | ab56416 |
| active Caspase-3 | FITC | C92-605 | BD Biosciences | 559341 |
| BrdU | FITC | B44 | BD Biosciences | 552598 |
| CD103 | FITC | M290 | BD Biosciences | 557494 |
| CD103 | PE | M290 | BD Biosciences | 557495 |
| CD11c | PerCP-Cy5.5 | HL3 | BD Biosciences | 560584 |
| CD36 | PE | CRF D-2712 | BD Biosciences | 562702 |
| CD3ε | FITC | 145-2C11 | BD Biosciences | 553062 |
| CD71 | FITC | C2 | BD Biosciences | 553266 |
| CD8alpha | APC | 53-6.7 | BD Biosciences | 561093 |
| Gr-1 | PE | RB6-8C5 | BD Biosciences | 553128 |
| I-A/I-E | Biotinylated | M5114.15.2 | BD Biosciences | 553622 |
| Ki67 | PerCP-Cy5.5 | B56 | BD Biosciences | 561284 |
| Ly-6G | FITC | 1A8 | BD Biosciences | 551460 |
| Beta3-tubulin | Uncoupled | TUJ1 | Biolegend | 801202 |
| CD11b | PerCP-Cy5.5 | M1/70 | Biolegend | 101228 |
| CD3e | APC | 145-2C11 | Biolegend | 100312 |
| CD3ε | PerCP-Cy5.5 | 145-2C11 | Biolegend | 100328 |
| CD45 | PE-Cy7 | 30F11 | Biolegend | 103114 |
| CD45 | APC-Cy7 | 30F11 | Biolegend | 103116 |
| CD86 | PE | GL-1 | Biolegend | 105008 |
| I-A/I-E | AlexaFluor 700 | M5114.15.2 | Biolegend | 107622 |
| TCRgd | PE | GL3 | Biolegend | 118107 |
| Fixable Viability Dye | eFluor450 | N/A | eBioscience | 65-0863-14 |
| Fixable Viability Dye | eFluor780 | N/A | eBioscience | 65-0865-14 |
| I-A/I-E | PE | M5114.15.1 | eBioscience | 12-5321-81 |
| TCRgd | APC | GL3 | eBioscience | 17-5711-82 |
| CD207 | AlexaFluor 647 | 929F3 | Eurobio/Dendritics | DDX0362A647-100 |
| CD207 | AlexaFluor 488 | 929F3 | Eurobio/Dendritics | DDX0362A488-100 |
| Bodipy | Bodipy 493/503 | N/A | Invitrogen | D3922 |
| Bodipy C11 | Bodipy 581/591 | N/A | Invitrogen | D3861 |
| Bodipy FL C16 | Bodipy 505/512 | N/A | Invitrogen | D3821 |
| DAPI | N/A | N/A | Invitrogen | D3571 |
| Donkey anti Rabbit IgG | AlexaFluor 647 | polyclonal | Invitrogen | A31573 |
| ER tracker Blue/White DPX | N/A | N/A | Invitrogen | E12353 |
| Lysosensor Green DND-189 | N/A | N/A | Invitrogen | L7535 |
| Lysotracker Red DND-99 | N/A | N/A | Invitrogen | L7528 |
| Mitosox Red | N/A | N/A | Invitrogen | M36008 |
| Mitotracker Deep Red 633 | N/A | N/A | Invitrogen | M22426 |
| Mitotracker Green FM | N/A | N/A | Invitrogen | M7514 |
| Mouse IgG(H+L) | AlexaFluor 555 | polyclonal | Invitrogen | A31570 |
| Mouse IgG(H+L) | AlexaFluor 594 | polyclonal | Invitrogen | A11032 |
| Streptavidin | AlexaFluor 488 | N/A | Invitrogen | S11223 |
| Streptavidin | AlexaFluor 546 | N/A | Invitrogen | S11225 |
| Poly (I:C) | N/A | N/A | InvivoGen | tlrl-pic |
| Guava Autophagy LC3 antibody based assay kit | FITC | 4E12 | Luminex | FCCH100171 |
| CD11b-coupled Microbeads | N/A | N/A | Miltenyi Biotec | 130-049-601 |
| 2-NBDG | N/A | N/A | Sigma-Aldrich | 72987 |
| Aluminium hydroxide | N/A | N/A | Sigma-Aldrich | 239186 |
| Etomoxir | N/A | N/A | Sigma-Aldrich | E1905 |
| Ferrostatin-1 | N/A | N/A | Sigma-Aldrich | SML0583 |
| Wortmannin | N/A | N/A | Sigma-Aldrich | W1628 |
| Fast SYBR Green Master Mix | N/A | N/A | ThermoFisher | 4385612 |
| RevertAid H Minus First Strand cDNA Synthesis Kit | N/A | N/A | ThermoFisher | K1632 |
| GLUT1 | Uncoupled | EPR3915 | Abcam | ab115730 |
