## Supplementary Table 2 for "Epidermal maintenance of Langerhans cells relies on autophagy-regulated lipid metabolism"

### Differentially expressed genes between $Atg5^{WT}$ and $Atg5^{\Delta\ Cd207}$

Genes with fold change (FC) >0 are increased in Langerin-cre x Atg5-floxed
Genes with fold change (FC) <0 are decreased in Langerin-cre x Atg5-floxed
FDR <0.1
p value <0.05

| Genes | baseMean | log2FoldChange | lfcSE | stat | pvalue | FDR |
|---|---|---|---|---|---|---|
| Oxct2b | 16.3971266001068 | -3.86859360361883 | 0.831108515955247 | -4.65473945862822 | 3.24390638420828e-06 | 0.000327929445385418 |
| Art2a-ps | 18.4102620289763 | -3.09631432478514 | 0.791863638648498 | -3.91016100962753 | 9.22346195040843e-05 | 0.00410259587554167 |
| Clu | 18.7374051105534 | -2.79505159369541 | 0.912478846387835 | -3.0631412495314 | 0.00219026649422018 | 0.0349939129536328 |
| Stra6 | 26.2206530492531 | -2.66014807815719 | 0.66710305834699 | -3.98761187626504 | 6.67417337298488e-05 | 0.00329852479589297 |
| Hcn3 | 20.7135432723618 | -2.62592466051472 | 0.742840042093054 | -3.53497995761755 | 0.000407806088740912 | 0.0117122489076641 |
| Gm12216 | 18.8021126816527 | -2.27875278864893 | 0.726133560629001 | -3.13820061790699 | 0.00169988487251056 | 0.029535499659871 |
| Mir17hg | 39.248794400744 | -2.15354450532231 | 0.560929918389386 | -3.83923986708687 | 0.000123415812691046 | 0.00514001437125256 |
| Pmel | 27.9876501353941 | -2.11681170433873 | 0.772326308187421 | -2.74082558356285 | 0.00612850301921884 | 0.0676300314890438 |
| Dqx1 | 20.0221076789565 | -2.07732523739429 | 0.719539942585185 | -2.88701865518517 | 0.00388911170674545 | 0.0506997915345949 |
| Dut | 21.3243583388932 | -2.04688406801382 | 0.572243151184179 | -3.57694812734425 | 0.000347629123893957 | 0.0106198787299473 |
| Cdc42ep3 | 57.3720248249748 | -2.02530263684992 | 0.413686366605021 | -4.89574421673808 | 9.79343829767342e-07 | 0.000132808577890401 |
| Slc4a8 | 55.4062209786223 | -2.0018304626321 | 0.566062416097585 | -3.53641295677719 | 0.000405600135434252 | 0.0117026063078899 |
| Chst7 | 24.0555306884723 | -1.9893840203953 | 0.787690852323698 | -2.52558984851302 | 0.0115504241920106 | 0.0994893237917567 |
| Slco5a1 | 396.585369251828 | -1.98154083484284 | 0.236818229432983 | -8.36734925173314 | 5.89288399387613e-17 | 1.07966847968394e-13 |
| Nrp1 | 24.9456525792033 | -1.97556290778168 | 0.736321100035891 | -2.68301819367309 | 0.00729610210006569 | 0.0747077857759949 |
| Nup210 | 47.7928636428196 | -1.95810881969532 | 0.61206428602144 | -3.19918816440588 | 0.00137815188323848 | 0.02549468761835 |
| Impg2 | 16.9341110926523 | -1.91240162351514 | 0.62274652204938 | -3.07091498033849 | 0.0021340390724722 | 0.034227506100699 |
| Tnfaip8l3 | 19.7068336531706 | -1.8677958915656 | 0.599817840254232 | -3.11393854303156 | 0.00184607938092365 | 0.0312456662342024 |
| Slc4a4 | 111.601936955643 | -1.86515875340635 | 0.404835529536583 | -4.60720123933146 | 4.08124980032035e-06 | 0.000395968592230861 |
| Cc2d2a | 72.3760340508022 | -1.86005161902814 | 0.363546044681177 | -5.11641275222611 | 3.11401226989578e-07 | 4.9468309201773e-05 |
| AW046200 | 42.0292291660399 | -1.81038363664006 | 0.449318295627915 | -4.02917854504472 | 5.59720962070338e-05 | 0.00292211131371932 |
| Gm9895 | 16.5633008183027 | -1.79091453377688 | 0.545096782072941 | -3.2854982687042 | 0.00101802168024188 | 0.0208984458871468 |
| Epb4.1l3 | 145.456743052766 | -1.78027471676007 | 0.363284492202323 | -4.90049741999057 | 9.5594313219012e-07 | 0.00013123564975252 |
| Catsperg2 | 29.067237161504 | -1.76841490689872 | 0.579533976370505 | -3.05144301974133 | 0.00227744270248124 | 0.0358170719032573 |
| Pla2g2d | 64.0738637538716 | -1.76606401376332 | 0.321874826304127 | -5.48680378034484 | 4.09271313070437e-08 | 9.68318510924099e-06 |
| Ccbl2 | 17.1611331613882 | -1.75725790225301 | 0.551723022923376 | -3.18503638463726 | 0.00144735809451857 | 0.0262984019788341 |
| Tgif2 | 23.3585973138008 | -1.73251030755263 | 0.660891985189228 | -2.62147271623603 | 0.00875507698842983 | 0.0843643467169321 |
| Oprd1 | 91.2414484966893 | -1.72287293399532 | 0.33628468779193 | -5.12325715841487 | 3.00302336618639e-07 | 4.83965504811487e-05 |
| Gm13315 | 189.032445003815 | -1.72198641481435 | 0.300523036418731 | -5.72996478185131 | 1.00451487892917e-08 | 2.93952775097168e-06 |
| Tnfrsf11b | 406.121739001542 | -1.69801533904225 | 0.290833889523238 | -5.83843699173364 | 5.26928026830096e-09 | 2.09265702083952e-06 |
| Dhdh | 36.3286498786432 | -1.69070062297746 | 0.359173926950928 | -4.70719196499039 | 2.5115237901881e-06 | 0.000271231811971041 |
| Ulbp1 | 25.4845008652404 | -1.68374152298861 | 0.508876445519758 | -3.30874328692667 | 0.000937157232255961 | 0.0200022810416243 |
| Ifi47 | 28.88128528552 | -1.67959201377206 | 0.524093070149315 | -3.20475905795424 | 0.00135175584715882 | 0.025136329465562 |
| Catsperg1 | 82.4091514847359 | -1.67138211181238 | 0.291167545869213 | -5.74027612460329 | 9.45223027003974e-09 | 2.88157041654001e-06 |
| Zbtb12 | 22.848104148279 | -1.65596058446415 | 0.618413057248733 | -2.67775811822502 | 0.0074116720638017 | 0.0753361913614944 |
| Heatr9 | 22.5382638771393 | -1.64638160644408 | 0.573103098921754 | -2.87274943992034 | 0.00406916703405469 | 0.051920407984837 |
| Trim14 | 164.775616574132 | -1.64348478542217 | 0.317509378791063 | -5.17617713114443 | 2.26478450489568e-07 | 3.95850486793888e-05 |
| Ccdc15 | 72.0772625565176 | -1.61388170185291 | 0.314471274903101 | -5.13204807768275 | 2.86606262261866e-07 | 4.75680841246559e-05 |
| Fam132b | 24.6063161753254 | -1.60937597731815 | 0.533844096406003 | -3.01469284413362 | 0.00257239620739342 | 0.0381400611016198 |
| Gm12359 | 21.6349061257357 | -1.60882516783653 | 0.540505263690426 | -2.97652081471306 | 0.00291539291186867 | 0.0417235124581462 |
| Pou5f2 | 43.7369223942442 | -1.59106644820284 | 0.511764076308188 | -3.10898424070838 | 0.0018773173141049 | 0.0314763302452612 |
| Ccl17 | 205.878564862702 | -1.57994564832262 | 0.47249740836918 | -3.34381865453142 | 0.000826337338624187 | 0.0183777424110019 |
| Arhgef9 | 191.890809878395 | -1.56934668854116 | 0.288702164987652 | -5.4358674054567 | 5.45304961420029e-08 | 1.18897866097857e-05 |
| Fam161a | 16.4159955241747 | -1.56258341820864 | 0.523240799273927 | -2.98635622523501 | 0.00282323559420883 | 0.0408197047166315 |
| Adam11 | 133.792006731641 | -1.54200852617032 | 0.450925194239781 | -3.4196548471194 | 0.00062700635677473 | 0.0152566973464661 |
| Gm5431 | 132.143993785818 | -1.54073716861303 | 0.35182865083624 | -4.37922598103066 | 1.19101577207138e-05 | 0.000843572954486223 |
| 4932438H23F | 142.846800792287 | -1.53526776790895 | 0.530765200473109 | -2.89255543984507 | 0.00382121736684213 | 0.0501459327170544 |
| Nudt17 | 692.88186833623 | -1.53156178244291 | 0.324659336603377 | -4.71744259218383 | 2.38827760360726e-06 | 0.000265576469521128 |
| Neurl1a | 30.2152894262207 | -1.52539941398719 | 0.471922840268875 | -3.2323068176105 | 0.00122795132295107 | 0.0233017384150443 |
| AU040972 | 31.680563812625 | -1.52269588653357 | 0.421063974981965 | -3.61630530514701 | 0.000298837893568952 | 0.00963210833764274 |
| Ccser1 | 76.2527241082074 | -1.52134819728725 | 0.430193690116544 | -3.53642610814469 | 0.000405579941987825 | 0.0117026063078899 |
| Stk39 | 34.9553435812687 | -1.5155823874926 | 0.559113720600337 | -2.71068716729984 | 0.00671439414780054 | 0.071448864041667 |
| Snord71 | 21.253724490363 | -1.49985971536552 | 0.506897772986948 | -2.95889979260994 | 0.00308739477922664 | 0.0434030719911473 |
| Phf11b | 35.293402403325 | -1.49672114640259 | 0.443251139022363 | -3.37668877671418 | 0.000733640094344283 | 0.0170314777643182 |
| Nlrc5 | 586.504473720151 | -1.48849411510256 | 0.239315101610913 | -6.21980854982817 | 4.97761862651021e-10 | 3.25594818392903e-07 |
| Gm10584 | 36.5157812446196 | -1.48731297315708 | 0.51277300110303 | -2.90052902543171 | 0.00372533334770163 | 0.049538755819916 |
| Ranbp17 | 42.7752497498449 | -1.47721533285922 | 0.476412505980504 | -3.10070645567745 | 0.00193059575164228 | 0.0319467630331282 |
| Prss53 | 37.7142541584081 | -1.47680325007189 | 0.482393103126862 | -3.0614103736129 | 0.00220296933930361 | 0.0350458069428558 |
| Arhgap22 | 114.063877117449 | -1.45866804327112 | 0.4569422040686 | -3.19223750899606 | 0.00141175195364403 | 0.0258627376021774 |

| | | | | | | |
|---|---|---|---|---|---|---|
| Pdgfb | 87.0518217095495 | -1.45477573797977 | 0.319920388052898 | -4.54730549320046 | 5.43370985305184e-06 | 0.000479546456872511 |
| Calcrl | 82.4827267129549 | -1.44417406937878 | 0.355063834117877 | -4.06736459928874 | 4.75478191726555e-05 | 0.00256511623193471 |
| Bcas3os1 | 47.9432937112571 | -1.42604708868304 | 0.384785095691429 | -3.70608712408818 | 0.000210485919013259 | 0.00747796619625381 |
| Apobec3 | 1035.65196281837 | -1.39984732547919 | 0.275750108819919 | -5.0765068832425 | 3.84436877897031e-07 | 5.77694335434458e-05 |
| Cetn4 | 49.5021435303144 | -1.39775025980266 | 0.437456856286814 | -3.19517282610891 | 0.0013974712908046 | 0.0256433675804409 |
| Gm5141 | 31.1820422801756 | -1.39538822255721 | 0.434081081433025 | -3.21457967702863 | 0.001306356544529 | 0.0244567078075687 |
| Msrb3 | 230.224699621125 | -1.39429921690312 | 0.412993919742613 | -3.37607686275884 | 0.00073527374963483 | 0.0170338418665402 |
| Gm20125 | 106.032414720803 | -1.39291656871548 | 0.402151433381736 | -3.46366182759138 | 0.000532875961377799 | 0.0137291739619298 |
| 1110028F11R | 31.0678342981078 | -1.39267005784461 | 0.511581633050765 | -2.72228314675718 | 0.00648325675610874 | 0.0699261058466821 |
| Arsb | 25.3352930308351 | -1.38686295517676 | 0.536261737608868 | -2.58616801817827 | 0.00970496097133862 | 0.0894854223781694 |
| Jak3 | 122.902144553898 | -1.36444902619845 | 0.267436475839218 | -5.10195560241662 | 3.3616162258202e-07 | 5.1918295043223e-05 |
| Htr7 | 326.198659832049 | -1.36440008881006 | 0.347072161148347 | -3.93117121320163 | 8.45330428575823e-05 | 0.00383676504725027 |
| Scube3 | 26.0262956634437 | -1.35276384376399 | 0.518881007587645 | -2.60707912600846 | 0.00913182553453919 | 0.0864220425055964 |
| Slc7a6 | 40.9343380828835 | -1.35178597394014 | 0.51790306605541 | -2.61011386597103 | 0.00905120897710174 | 0.0862463100474476 |
| Dpf3 | 21.0166465870187 | -1.34850545432902 | 0.47784951020044 | -2.82202958367243 | 0.00477207749407889 | 0.0578055574446157 |
| Kank3 | 67.9752504379948 | -1.34128135965636 | 0.290607003089762 | -4.6154474785388 | 3.92249233036461e-06 | 0.000386001015165083 |
| Pyhin1 | 403.50125322369 | -1.33054674662209 | 0.32823817900492 | -4.05360141424057 | 5.04351206295049e-05 | 0.00268343799712964 |
| Gm3696 | 39.8594851754878 | -1.32748795241313 | 0.488064218808328 | -2.71990426926679 | 0.00653008182344102 | 0.0703628971673102 |
| Armc9 | 18.5893159752676 | -1.32694090852341 | 0.518977752838559 | -2.55683582054468 | 0.0105629058586439 | 0.0942692721895028 |
| Pkp1 | 80.5120048229748 | -1.30067060124135 | 0.412373326674953 | -3.15410943702134 | 0.00160988703449181 | 0.0284243910192768 |
| Tm7sf2 | 31.4282224140363 | -1.29421190182154 | 0.462288878910212 | -2.79957394794459 | 0.00511700947416829 | 0.0604046129010099 |
| Igsf9 | 64.7794615514052 | -1.28402892632965 | 0.425140165886746 | -3.02024844830047 | 0.00252567412181656 | 0.0376426074219372 |
| Tmem201 | 50.2055738633344 | -1.27812118911245 | 0.34896267342893 | -3.6626300932233 | 0.0002496389003647 | 0.00846171996505804 |
| F630111L10R | 308.343417768654 | -1.27807310295135 | 0.247314472460765 | -5.16780554827463 | 2.36858546901972e-07 | 4.05210314084604e-05 |
| Sema4a | 853.677127350477 | -1.27798111522666 | 0.285768553274396 | -4.47208449139448 | 7.74608085822632e-06 | 0.000633356023113799 |
| Eno3 | 151.59112491347 | -1.27482968774342 | 0.230043614228895 | -5.54168691887685 | 2.99571566450341e-08 | 7.48272761481306e-06 |
| Il15ra | 337.062744583124 | -1.26768150559758 | 0.279226748808536 | -4.53997158584122 | 5.62618096588221e-06 | 0.000492623089296143 |
| Slc27a3 | 1025.62159969429 | -1.26413786406194 | 0.495372681683647 | -2.55189256655303 | 0.0107139549892103 | 0.0951166669810377 |
| Acot1 | 35.6776604888679 | -1.25629184083345 | 0.458779414869944 | -2.73833524372401 | 0.00617510925599497 | 0.0676300314890438 |
| Slc25a29 | 122.330808366085 | -1.25610313466659 | 0.391558302191959 | -3.20795939617389 | 0.00133680370409484 | 0.0249417066938501 |
| Extl1 | 233.303812949817 | -1.25331541290636 | 0.275150766166156 | -4.5550133491161 | 5.23822517118859e-06 | 0.000469719716928102 |
| Pde1b | 207.874812444011 | -1.24431395843393 | 0.230870004245854 | -5.38967356326142 | 7.05857803927581e-08 | 1.4809695810707e-05 |
| Sh3bp1 | 508.52567830231 | -1.24198955041485 | 0.274062469524293 | -4.53177537431755 | 5.84900267002406e-06 | 0.00050666643535405 |
| Casp7 | 136.509460268506 | -1.23633014026956 | 0.271084055567175 | -4.56068925810799 | 5.09859850078706e-06 | 0.000460946466087415 |
| 4921511I17R | 249.784972517925 | -1.23536786460045 | 0.318263394269741 | -3.88158954766071 | 0.000103775925121961 | 0.00451496046078988 |
| Isyna1 | 29.6690347740127 | -1.23256422078878 | 0.434150359059508 | -2.83902614628457 | 0.00452514488753492 | 0.0557249292905739 |
| Snora52 | 22.7306027952009 | -1.22036524704576 | 0.416872004019524 | -2.92743392523092 | 0.00341771673164745 | 0.0465747672253917 |
| Gli3 | 101.953180280839 | -1.21352296304908 | 0.415185918863501 | -2.92284229284771 | 0.00346852147309323 | 0.0472092518736802 |
| Il15 | 209.240346530033 | -1.20643944800977 | 0.241541060624247 | -4.99475925497639 | 5.89092068653578e-07 | 8.50740753691921e-05 |
| Gpr85 | 305.652610870742 | -1.20294971193956 | 0.272764336883782 | -4.41021625364502 | 1.03267439229548e-05 | 0.000772766640307479 |
| Mir155hg | 493.740249308643 | -1.2010485607314 | 0.302837068552352 | -3.96598925776412 | 7.30921543709271e-05 | 0.00350338257157202 |
| Fah | 90.3658548743602 | -1.19831416332375 | 0.357969137557692 | -3.34753485034901 | 0.000815337354369936 | 0.0182793374608744 |
| Gpr52 | 111.538276239045 | -1.1927115934251 | 0.341903520718937 | -3.48844490082211 | 0.000485838942552005 | 0.012771936267561 |
| 2610206C17F | 31.2989803690609 | -1.19265984642401 | 0.466925567279051 | -2.55428258806661 | 0.010640685848644 | 0.0947353295731959 |
| F730043M19 | 89.8348976204775 | -1.18964857278501 | 0.405018607759214 | -2.93726893035063 | 0.00331116825350787 | 0.0454570259000093 |
| 9930111J21R | 75.5446618230923 | -1.18741591725257 | 0.387944461928203 | -3.06078842149401 | 0.00220755029265554 | 0.0350685132204708 |
| Lmo2 | 307.880530090909 | -1.18663270576788 | 0.217924754176468 | -5.44514876362778 | 5.17621069391215e-08 | 1.17468291665925e-05 |
| Zfp296 | 152.763421572572 | -1.17419940191333 | 0.288442380883424 | -4.07082828229702 | 4.68462731270589e-05 | 0.00254112466913607 |
| Epha1 | 96.7984728225612 | -1.17169570030298 | 0.426059032039978 | -2.75007830415632 | 0.00595810249595361 | 0.0666435371140802 |
| Vdr | 604.28386583523 | -1.17130634349889 | 0.275249844227685 | -4.25542963261404 | 2.08647978492575e-05 | 0.00133171521934482 |
| Mks1 | 28.2353168423433 | -1.16846287513127 | 0.438068424371066 | -2.66730677247242 | 0.0076461843070322 | 0.0769462167368308 |
| Icosl | 78.0279037633074 | -1.15835597423882 | 0.431669793550024 | -2.68343069528348 | 0.00728710770069647 | 0.0746844586467693 |
| A530099J19R | 125.786154090552 | -1.15670408084015 | 0.269517961922915 | -4.29175136450084 | 1.77269379267322e-05 | 0.00116641153695421 |
| 5430437J10R | 83.9121937064343 | -1.14703740569374 | 0.409027762241064 | -2.80430208308873 | 0.00504256356205819 | 0.0598677099666391 |
| 9330133O14I | 28.4333868842631 | -1.14503038764648 | 0.418044484129589 | -2.73901565770101 | 0.00616234385373127 | 0.0676300314890438 |
| Wee1 | 51.9424152977771 | -1.14451839420226 | 0.446301639420226 | -2.56445034727872 | 0.0103339363866333 | 0.0929719843198728 |
| Rab39 | 88.8553701365288 | -1.14206576548737 | 0.326775200486534 | -3.49495850293092 | 0.000474135465911921 | 0.0127045454962423 |
| Adat1 | 45.2123574387132 | -1.13543344984893 | 0.36290862831482 | -3.12870337396318 | 0.00175579457914713 | 0.0302235847060621 |
| Bckdhb | 42.0058553013178 | -1.13511636851706 | 0.414894274239531 | -2.73591716973593 | 0.00622066819829438 | 0.0678174807500329 |
| Fbxw17 | 118.294535314723 | -1.13308194034215 | 0.27984929224826 | -4.04890050369314 | 5.14588141393327e-05 | 0.00272486672966371 |
| 4930520O04I | 99.5908090964843 | -1.13236682604253 | 0.394224396159335 | -2.87239155433916 | 0.00407377871022444 | 0.051920407984837 |
| 2200002D01I | 72.601716262475 | -1.12980021412848 | 0.320133530166017 | -3.52915301793781 | 0.000416892002358129 | 0.0117122489076641 |
| Pla2g16 | 305.841541742135 | -1.11719393117309 | 0.253794588087303 | -4.40196120647297 | 1.0727674116618e-05 | 0.000795278241178612 |
| Gm10432 | 427.776933559322 | -1.10998744586763 | 0.260008326643395 | -4.26904576556118 | 1.96310991034537e-05 | 0.00126183712156303 |
| Gbp9 | 189.704891437662 | -1.106175368073 | 0.248668285779868 | -4.44839744884974 | 8.65133368346154e-06 | 0.000682289578440371 |
| Tyw3 | 54.9125746881925 | -1.10410716852332 | 0.421801222915275 | -2.61760068141172 | 0.00885503612750982 | 0.0849594492993177 |
| Plek2 | 1325.36865247543 | -1.10292006374521 | 0.263845877936389 | -4.18016787820015 | 2.91293967024101e-05 | 0.00174149941575699 |
| Yes1 | 715.88084654252 | -1.10200040048973 | 0.216269260499258 | -5.09550177378771 | 3.47818764242363e-07 | 5.29828035393846e-05 |
| Zbp1 | 41.5767496307673 | -1.09951579104124 | 0.387623027719718 | -2.83655952410618 | 0.00456024770167556 | 0.0559712521441856 |

| | | | | | | |
|---|---|---|---|---|---|---|
| Rragd | 79.583928707668 | -1.09875581165574 | 0.346650257430091 | -3.16963795094635 | 0.00152628986706655 | 0.027286725597717 |
| Suv39h2 | 562.085787381353 | -1.09469795545921 | 0.279703737520004 | -3.9137766451223 | 9.08636895635431e-05 | 0.00405784830500642 |
| Far1os | 52.082952557101 | -1.08326435939406 | 0.31456168947038 | -3.44372628853161 | 0.000573756232386443 | 0.0144021880454565 |
| Fam196b | 127.264724486916 | -1.08279827795817 | 0.306691528632399 | -3.53057772018219 | 0.000414653173329111 | 0.0117122489076641 |
| Irak3 | 209.133825927038 | -1.08174086004908 | 0.394600575996356 | -2.74135651555429 | 0.00611860779152768 | 0.0676300314890438 |
| Nrg1 | 287.41456568513 | -1.0749030798167 | 0.310614892891936 | -3.46056517061678 | 0.000539042739455423 | 0.0138165894044565 |
| Fes | 697.498240695771 | -1.06934707779075 | 0.237603352227969 | -4.50055551726714 | 6.7776099401114e-06 | 0.000562440466671931 |
| Mycl | 961.602398343311 | -1.06009400656069 | 0.24201925936704 | -4.38020515116517 | 1.18567627882312e-05 | 0.000843572954486223 |
| Chst10 | 69.7151115803465 | -1.05791652576947 | 0.386327801817279 | -2.73839087115411 | 0.00617406472397457 | 0.0676300314890438 |
| 4632415L05R | 67.9492419697652 | -1.05575579462567 | 0.324424376532146 | -3.25424311795222 | 0.00113694933758834 | 0.0223311958855458 |
| Lrrc4 | 528.085172786225 | -1.05554415982268 | 0.237624612143162 | -4.44206578730466 | 8.90993076019661e-06 | 0.000697735422911171 |
| Tk1 | 501.838325980169 | -1.05249259683757 | 0.317088352307493 | -3.31924080206179 | 0.00090262556822454 | 0.0195656848316898 |
| Fam212a | 63.9398157309771 | -1.05183306151812 | 0.403666735148403 | -2.60569665501772 | 0.00916876231964346 | 0.086477215432091 |
| Eps8l1 | 108.245398023547 | -1.04673892979119 | 0.34253516729552 | -3.05585828764882 | 0.00224417297016033 | 0.0353974516711814 |
| Irf1 | 1529.64199512788 | -1.04500269129615 | 0.159337825241919 | -6.55840940284919 | 5.43847311439267e-11 | 4.31970150228903e-08 |
| 4930594C11F | 157.125687790898 | -1.04434392364896 | 0.320969065786936 | -3.25372141732253 | 0.00113903920865621 | 0.0223311958855458 |
| Cbfa2t3 | 1694.54005794704 | -1.03918819053926 | 0.194400810835111 | -5.34559596781047 | 9.01200331592777e-08 | 1.78952637273423e-05 |
| Gramd1b | 250.747438684645 | -1.03835234524467 | 0.29763032048282 | -3.4887317379501 | 0.000485317940338701 | 0.012771936267561 |
| Cd48 | 379.360214745581 | -1.03690044632886 | 0.272710494729155 | -3.80220221212487 | 0.000143415596078129 | 0.00569564795853141 |
| 4930487H11F | 180.693481200834 | -1.03541690241554 | 0.31475303128011 | -3.28961693618795 | 0.0010032386037118 | 0.020813457599394 |
| Ptp4a3 | 31.6693752592607 | -1.03347813525587 | 0.378258289812198 | -2.73220220968318 | 0.00629125155538563 | 0.0683190598592659 |
| Thap2 | 211.486774133382 | -1.0302208781971 | 0.224191840048364 | -4.59526483200662 | 4.32199988313238e-06 | 0.000414315850865794 |
| Rundc3b | 85.2742613785078 | -1.02544348056713 | 0.386533112366561 | -2.65292531935703 | 0.00797975239222975 | 0.0790158918981254 |
| Samhd1 | 3752.6843713263 | -1.02192953550419 | 0.17596549109865 | -5.8075565221551 | 6.33911548900184e-09 | 2.34969880792335e-06 |
| Coro2a | 2953.82527175038 | -1.01835606061129 | 0.292423798336456 | -3.48246642853462 | 0.000496817510444649 | 0.0129381983984649 |
| Pard3b | 41.6658451369118 | -1.01604167947763 | 0.392022405383143 | -2.59179492173312 | 0.00954766672614519 | 0.0888132097150725 |
| Cbx7 | 184.464653746997 | -1.01475288765491 | 0.230747452282989 | -4.39767753713013 | 1.09415376152879e-05 | 0.000805760915774844 |
| Socs2 | 2774.2654490614 | -1.01461142780973 | 0.221304103587144 | -4.58469324049474 | 4.54653435211876e-06 | 0.00042999819888111 |
| Wfdc13 | 95.2660715610508 | -1.01368083363548 | 0.295834921564761 | -3.42650836579337 | 0.000611394928079049 | 0.015022150300123 |
| Olr1 | 294.389801395007 | -1.01254005132683 | 0.241574692636935 | -4.19141608036149 | 2.77218727772906e-05 | 0.001693775963096 |
| Tmcc2 | 88.8947248520483 | -1.00952612990314 | 0.300921846092601 | -3.35477846826874 | 0.000794285648228348 | 0.018025421241427 |
| Zfp39 | 79.2062229092097 | -1.00325229435382 | 0.228435422663146 | -4.39184204733971 | 1.12394342541345e-05 | 0.000816879143176314 |
| Vangl1 | 150.252448267764 | -1.00128754193063 | 0.283391831589767 | -3.53322654472297 | 0.000410520531311051 | 0.0117122489076641 |
| Cacna1d | 139.301803527539 | -0.999136728773638 | 0.292835725835456 | -3.41193591022105 | 0.000645032768888739 | 0.0155591418439106 |
| Gda | 252.46469499865 | -0.996177346722944 | 0.356891108179937 | -2.79126412480048 | 0.00525026094011689 | 0.0613910637792848 |
| Prdm8 | 218.978988652664 | -0.995624195246181 | 0.33853382994179 | -2.94098877922297 | 0.00327166384950079 | 0.0450258688198623 |
| L1cam | 451.853980504449 | -0.988751806589321 | 0.228514017918578 | -4.3268759422086 | 1.51239084853661e-05 | 0.00101925977186225 |
| Suv39h1 | 215.716379121371 | -0.986914350870141 | 0.266087546188028 | -3.7089836221525 | 0.00020809288612179 | 0.00744049161953154 |
| Zfp961 | 235.83746878233 | -0.978074707745352 | 0.169121093758548 | -5.78328040582411 | 7.32577985808414e-09 | 2.51426248195699e-06 |
| Tmem173 | 410.048358600078 | -0.976414509214688 | 0.280544732182698 | -3.48042360880553 | 0.000500621564004864 | 0.0130068032517152 |
| Ttll3 | 187.559727057778 | -0.969275648057435 | 0.281298306066978 | -3.44572159573065 | 0.00056953679316591 | 0.0143611091610089 |
| Atp10d | 227.512840375934 | -0.967668538673857 | 0.284049983468625 | -3.40668401686685 | 0.000657572067131298 | 0.0158198992577674 |
| Fscn1 | 56746.7470419766 | -0.966884683891194 | 0.231491735522368 | -4.1767568147061 | 2.956948593707e-05 | 0.00174900363627776 |
| Nfix | 51.1890148769438 | -0.96359437215083 | 0.341423670397083 | -2.82228344341254 | 0.00476830133686368 | 0.0578055574446157 |
| Tusc1 | 52.1616139821169 | -0.962734825636809 | 0.362140262523476 | -2.65845840760222 | 0.0078499039263993 | 0.0781476559190333 |
| Cpne8 | 110.494102915444 | -0.962432738061801 | 0.307301230854254 | -3.13188702624579 | 0.00173686695071211 | 0.0300476613112945 |
| Stap2 | 249.69883020229 | -0.961901125977026 | 0.229195785675313 | -4.19685345933756 | 2.70648927960418e-05 | 0.00167200893273325 |
| Cdkl5 | 82.8126745695314 | -0.95820635478354 | 0.279365103176894 | -3.42994290942918 | 0.000603708257309945 | 0.0148852235505246 |
| Gbp5 | 456.011176226864 | -0.953085194110999 | 0.340972082391361 | -2.79520008625535 | 0.00518676025433516 | 0.0610134356419227 |
| Mex3b | 154.532874991742 | -0.951796473381763 | 0.276743738306364 | -3.43927013202408 | 0.000583284896755125 | 0.0145755686559932 |
| Arhgef4 | 34.4989615766601 | -0.94990326203397 | 0.352334378276036 | -2.69602775261905 | 0.00701718199923833 | 0.0729262278799348 |
| Gm16617 | 98.0681726151253 | -0.949435179362821 | 0.335243188422751 | -2.83207895686028 | 0.00462464274652987 | 0.0566990378626375 |
| Mamld1 | 646.8482167363 | -0.948459523503147 | 0.204250021886483 | -4.64362018051719 | 3.42356694068896e-06 | 0.000342973552977128 |
| Gm10389 | 113.374314765439 | -0.947977652876575 | 0.273711769323459 | -3.46341575015104 | 0.000533363592765618 | 0.0137291739619298 |
| Adap1 | 581.819924973986 | -0.945601064393245 | 0.204191246366196 | -4.63095789472487 | 3.63977892282341e-06 | 0.00036137805019461 |
| Fam120c | 58.2272161182946 | -0.945331164284684 | 0.311706822335656 | -3.03275737502697 | 0.00242330350617432 | 0.0367628035315941 |
| Tmem176a | 527.540947551391 | -0.937030947593796 | 0.237842596123166 | -3.93971039194576 | 8.15800309802861e-05 | 0.00377987476875325 |
| Mab21l3 | 155.200462095374 | -0.936415449342312 | 0.283466602741893 | -3.30344188798478 | 0.000955057616765078 | 0.0201905716700146 |
| Ndnf | 108.433204578079 | -0.935206211575683 | 0.342619771374909 | -2.72957455964309 | 0.00634161069001311 | 0.0687316870106684 |
| Rn4.5s | 639.30262800319 | -0.929887120520759 | 0.266521612622631 | -3.48897453895189 | 0.000484877331264999 | 0.012771936267561 |
| Adgrl1 | 64.8998541445275 | -0.926775346921403 | 0.331885963966402 | -2.79245116559139 | 0.00523103629127589 | 0.0612574214906234 |
| Ptpro | 643.52203094598 | -0.925194452000229 | 0.130814112576084 | -7.07258898738557 | 1.52069402227577e-12 | 1.69101175277066e-09 |
| Chka | 100.398639336512 | -0.92449136453938 | 0.32207090245844 | -2.87045913642781 | 0.0040987617033698 | 0.051920407984837 |
| Uba7 | 507.073731114986 | -0.915907524253158 | 0.154593210222328 | -5.92462969709954 | 3.13001894974975e-09 | 1.45024211338405e-06 |
| Dcaf4 | 102.589891441111 | -0.907425005848448 | 0.3481902740597 | -2.60611818724398 | 0.00915748573774318 | 0.086477215432091 |
| Kynu | 290.826805590841 | -0.906098762428742 | 0.25871668099703 | -3.50228195158062 | 0.000461291185068219 | 0.0125111170194112 |
| Ttc7 | 594.264886658794 | -0.905904547571394 | 0.189919824060848 | -4.76993148056599 | 1.84288594377831e-06 | 0.000219890846249786 |
| Tmem176b | 1613.67365621972 | -0.902338167870135 | 0.265518868574708 | -3.39839564967206 | 0.000677823155376779 | 0.0161400288817768 |
| Oasl2 | 651.6206826062 | -0.899419532667931 | 0.326925387949546 | -2.75114618142393 | 0.00593871354822164 | 0.0665710631615167 |

| | | | | | | |
|---|---|---|---|---|---|---|
| Slc50a1 | 55.3737080217337 | -0.89716940783496 | 0.332144731632848 | -2.7011399621617 | 0.0069102251947742 | 0.0724921737414048 |
| Prune2 | 50.1123911229979 | -0.894128492443807 | 0.32650321747406 | -2.73849825848912 | 0.00617204873156576 | 0.0676300314890438 |
| Sh2b3 | 1117.20172137169 | -0.888262700228601 | 0.221370457714173 | -4.01256206180636 | 6.00632808880093e-05 | 0.00303592583397574 |
| Crtap | 300.780830026072 | -0.884296614476071 | 0.238056268005277 | -3.71465377444491 | 0.000203482101716926 | 0.00732272158929521 |
| Tbrg3 | 301.239188776231 | -0.87937806462404 | 0.23838746382171 | -3.68886035585212 | 0.000225260807835767 | 0.00790189332218843 |
| Selplg | 4879.53744680852 | -0.877151864211359 | 0.258818875325879 | -3.38905677998538 | 0.000701334864582033 | 0.0165229739282886 |
| Klf8 | 104.883854023486 | -0.875450947941073 | 0.298952248232674 | -2.92839727119132 | 0.00340714401623353 | 0.0465447683790133 |
| Dtymk | 59.5574910005044 | -0.874988588658625 | 0.328362534305179 | -2.66470287333516 | 0.0077056372297028 | 0.0771952126074731 |
| 2310009A05F | 40.263713003342 | -0.874708601293045 | 0.342911157089415 | -2.55083155858053 | 0.0107466250593117 | 0.095297026044295 |
| Prr33 | 692.748166606634 | -0.867830772120763 | 0.237799350019789 | -3.64942449190271 | 0.000262828479801774 | 0.00880317076926424 |
| St3gal4 | 361.128900935894 | -0.865968742828134 | 0.232887985450748 | -3.71839166005956 | 0.000200495246881705 | 0.00726223825838619 |
| Gne | 124.230190375176 | -0.865629304300979 | 0.229577850473883 | -3.77052621807457 | 0.000162903686290376 | 0.00624272372995241 |
| Flt3 | 4265.03384864169 | -0.857985471328833 | 0.258450899770432 | -3.31972329015273 | 0.000901067129572766 | 0.0195656848316898 |
| Dennd3 | 1298.97398346296 | -0.85488379085157 | 0.271045682950143 | -3.15402105485229 | 0.00161037467105615 | 0.0284243910192768 |
| Slc22a23 | 1269.91323240637 | -0.853489399693434 | 0.225591836707973 | -3.78333459290138 | 0.000154741172403104 | 0.00599554647081016 |
| Paqr3 | 117.151775428784 | -0.851653684674303 | 0.274024863955755 | -3.10794309822854 | 0.00188394337022745 | 0.0314763302452612 |
| 4930579G24F | 176.177697133553 | -0.850658351469659 | 0.328725265996223 | -2.58774861400351 | 0.0096605454168316 | 0.0892391592827107 |
| Slc24a1 | 476.4101200105 | -0.84873765820399 | 0.308711019885206 | -2.74929498311914 | 0.00597236113707072 | 0.0666435371140802 |
| Lpcat1 | 571.885583491128 | -0.847915292293993 | 0.2389260153784 | -3.5488613115282 | 0.000386900799060767 | 0.0112922227967342 |
| Dock8 | 2812.12093301211 | -0.847453678347236 | 0.217862954670455 | -3.88984754029942 | 0.000100307220483384 | 0.00439140272352454 |
| Bcl2l1 | 1434.03385185298 | -0.846011853670069 | 0.234887795710745 | -3.60177016055742 | 0.000316057740029919 | 0.0100416059118077 |
| Prdm9 | 126.486384706348 | -0.844747735433521 | 0.297208050960418 | -2.84227743058691 | 0.00447924944427826 | 0.0553436153559714 |
| Gdap10 | 184.743163176208 | -0.843903567917936 | 0.289620713118046 | -2.91382325121881 | 0.00357031987412198 | 0.0481767005079158 |
| Rasip1 | 371.5578264993 | -0.843755608521662 | 0.227599736161282 | -3.70719062663482 | 0.000209571194236127 | 0.00746933230739017 |
| Fam53b | 1077.18900039826 | -0.842149318627823 | 0.203011701010536 | -4.14827970228237 | 3.34982978695081e-05 | 0.00193005736947632 |
| B4galt3 | 100.227694387254 | -0.841391155606758 | 0.290905245305808 | -2.89232033173641 | 0.00382407832230774 | 0.0501459327170544 |
| Dpysl5 | 487.284445383765 | -0.839809623488057 | 0.163815469764049 | -5.12655871083284 | 2.95086065516995e-07 | 4.82552507139557e-05 |
| Rbm38 | 152.78379433635 | -0.838994574779491 | 0.332025308525859 | -2.52689946590065 | 0.0115074438305589 | 0.0993499808973721 |
| Isoc1 | 206.846179608693 | -0.837915565979181 | 0.195349387578294 | -4.28931759841506 | 1.79222932987634e-05 | 0.00117232883224852 |
| Pprc1 | 324.102664163841 | -0.83766010213004 | 0.1618645499729 | -5.17506830414865 | 2.27827618298641e-07 | 3.95850486793888e-05 |
| B3gntl1 | 56.776894367744 | -0.836703744594289 | 0.314287238106809 | -2.66222627948367 | 0.00776256753912081 | 0.0776955454860696 |
| Tmem150c | 585.457223856136 | -0.834030907796396 | 0.323564996928423 | -2.57763020015696 | 0.00994803963388539 | 0.0911220763828711 |
| Phf11c | 85.3410220204571 | -0.833990014890022 | 0.26173951153397 | -3.18633594905973 | 0.00144087177323249 | 0.026223394629043 |
| Pisd-ps1 | 342.021485687997 | -0.833417967860979 | 0.173214971087985 | -4.81146613728694 | 1.49827122343717e-06 | 0.000187199730389004 |
| Gtpbp1 | 1398.94385537014 | -0.830959384922029 | 0.174252836971145 | -4.76869931856334 | 1.85419163495449e-06 | 0.000219890846249786 |
| Foxp4 | 1351.92680847926 | -0.830076107626307 | 0.235258382759416 | -3.52835932088836 | 0.00041814413816031 | 0.0117122489076641 |
| Lsr | 267.938973289027 | -0.827845974549261 | 0.247955590365227 | -3.33868646933865 | 0.000841755132234046 | 0.018626875186688 |
| Gm5512 | 68.8064848570149 | -0.827484227717424 | 0.279506201439184 | -2.96052188987825 | 0.00307118295985213 | 0.043229815839944 |
| Mum1 | 186.34109430364 | -0.826119059118522 | 0.218303907044999 | -3.78426144681063 | 0.00015416569691585 | 0.00599413478917572 |
| Lime1 | 226.846949834009 | -0.825858796604004 | 0.261513740432534 | -3.15799389828643 | 0.00158858882944136 | 0.0281740156034895 |
| Tmem39a | 1768.96383588953 | -0.825753141619697 | 0.148043718671186 | -5.57776546706276 | 2.43627818989448e-08 | 6.45033654086347e-06 |
| 5530601H04F | 99.6619918915372 | -0.823818031610191 | 0.264337908633852 | -3.11653381789937 | 0.0018299069083624 | 0.0310665111770838 |
| Dscam | 175.754607444648 | -0.820096803362257 | 0.253253981569119 | -3.23823853935514 | 0.00120270204223063 | 0.0230985262687471 |
| Plk4 | 260.466716238355 | -0.818205388912195 | 0.203574285937931 | -4.01919812781101 | 5.8396549369964e-05 | 0.00300634087497222 |
| Arhgef40 | 1919.64072027741 | -0.817713915141471 | 0.199306754839506 | -4.10279077495364 | 4.08196355831719e-05 | 0.00225828033674065 |
| Mir3064 | 75.5531059034685 | -0.817616580198607 | 0.301539190544509 | -2.71147700145438 | 0.00669841876806238 | 0.0714155481312116 |
| Chst3 | 607.914512757042 | -0.816754966827857 | 0.259103059085444 | -3.15223976787753 | 0.00162023170726027 | 0.0284628382065311 |
| Zmym6 | 266.191219809019 | -0.814391343392143 | 0.151952185224845 | -5.35952373562171 | 8.34416312410741e-08 | 1.68703807163772e-05 |
| Nostrin | 1038.6532374894 | -0.813032058607083 | 0.216273472067194 | -3.75927778305831 | 0.000170404555082598 | 0.00646511886180764 |
| Sp110 | 610.257448878014 | -0.810653186533734 | 0.255667739791069 | -3.17072927228206 | 0.00152056784926075 | 0.0272530472967428 |
| N4bp2l1 | 539.452077224815 | -0.809386545052846 | 0.1395159345079 | -5.80139141745858 | 6.57668787097933e-09 | 2.35912158468678e-06 |
| Xrcc3 | 164.553426502829 | -0.80887751791248 | 0.225665152950251 | -3.58441481698683 | 0.000337834797962282 | 0.0104564092437121 |
| Gabbr1 | 100.155176260319 | -0.808326338844219 | 0.250039882509049 | -3.23278962833045 | 0.00122587800660483 | 0.0233017384150443 |
| Spint2 | 216.455824910492 | -0.807310413239157 | 0.314342089958017 | -2.56825426511283 | 0.0102212152716567 | 0.0924064340006689 |
| Bahcc1 | 526.232421604405 | -0.803485685559134 | 0.224846450745559 | -3.57348618532732 | 0.000352259836008813 | 0.0107025392798306 |
| Ccdc120 | 57.1356207397335 | -0.798897005165323 | 0.31305083300111 | -2.55197214301132 | 0.0107115082711089 | 0.0951166669810377 |
| Gnb4 | 302.604596029173 | -0.798748555527149 | 0.237502859078925 | -3.36311132684814 | 0.000770692771640155 | 0.0176379509592121 |
| Efnb2 | 203.671054733733 | -0.797653962226089 | 0.313220174179082 | -2.54662383838033 | 0.010877061037937 | 0.0960706264828115 |
| Psmg4 | 232.989274918393 | -0.794177699067837 | 0.297035122114623 | -2.67368280698257 | 0.00750233741911112 | 0.0758687928152643 |
| Cnp | 146.144026991615 | -0.794092115153715 | 0.287996191082652 | -2.75730075515415 | 0.00582807192425191 | 0.0656009820830332 |
| Ppp4r1l-ps | 109.763502025896 | -0.793925893807981 | 0.293212070006532 | -2.70768489779529 | 0.00677543176393282 | 0.0717743839738763 |
| Mcm5 | 161.124391910344 | -0.79293970483906 | 0.276193934596779 | -2.8709526369458 | 0.00409236836553157 | 0.051920407984837 |
| Rsad1 | 49.2066816884308 | -0.792832502144262 | 0.289076558412589 | -2.74263851243407 | 0.00609477391114282 | 0.0675038704102671 |
| Ccdc28b | 183.386027934109 | -0.789285563915721 | 0.263812497505289 | -2.99184296187446 | 0.00277298866932027 | 0.0404136749709586 |
| Dock9 | 193.250728532578 | -0.788323492775622 | 0.190872814609197 | -4.13009832955876 | 3.62608111005077e-05 | 0.00206779599711613 |
| Plekhn1 | 412.004856387974 | -0.787948804599844 | 0.207204934239469 | -3.80275116271702 | 0.000143098070601167 | 0.00569564795853141 |
| Shb | 140.543152674438 | -0.787872075234098 | 0.271126556591997 | -2.90591997013307 | 0.00366175090853317 | 0.0488819569062291 |
| Acsf2 | 447.029941099304 | -0.784890691546106 | 0.272230323817043 | -2.88318612174008 | 0.00393674760162087 | 0.0512007407368702 |
| Pira2 | 268.360936651234 | -0.783712548583665 | 0.222279376806242 | -3.52579964837141 | 0.00042220624405045 | 0.0117667504607544 |

| | | | | | | |
|---|---|---|---|---|---|---|
| Deptor | 524.466151595417 | -0.783036989327256 | 0.285642064618638 | -2.74132239722007 | 0.00611924323797539 | 0.0676300314890438 |
| Zfc3h1 | 8717.89047117876 | -0.782116621308875 | 0.162324439854136 | -4.81823083456615 | 1.44836747882003e-06 | 0.000183020981414532 |
| Snord32a | 66.5258454212701 | -0.781332929400795 | 0.306964089168502 | -2.54535614089991 | 0.0109166335218947 | 0.0961147781183443 |
| Tcaf2 | 3222.03506708055 | -0.781081547882842 | 0.266446122142905 | -2.93148026175408 | 0.00337350803277219 | 0.0461419548885938 |
| 9430076C15F | 217.051058597388 | -0.780985593038918 | 0.227981259653799 | -3.42565697823094 | 0.000613314409735237 | 0.015022150300123 |
| Sfi1 | 272.705443779016 | -0.777163928948402 | 0.217839956067172 | -3.56759128572704 | 0.000360277884074599 | 0.0107740972421658 |
| Lfng | 386.678687132135 | -0.774895490721384 | 0.257605348268942 | -3.00807221561404 | 0.00262910654977946 | 0.0388255841082969 |
| Setd6 | 79.9749591866123 | -0.774664449426153 | 0.251696865838324 | -3.0777675631597 | 0.00208557518316698 | 0.0338915392973821 |
| Cryaa | 162.143268893492 | -0.774502652866341 | 0.269624367148831 | -2.87252469447177 | 0.00407206252657583 | 0.051920407984837 |
| Fabp5 | 3089.77822543706 | -0.773042550367493 | 0.284488187460288 | -2.71730983725081 | 0.00658149637208992 | 0.0705069746220037 |
| Npepl1 | 151.032765564656 | -0.772008507391153 | 0.255310715465144 | -3.02379986670224 | 0.00249621509815163 | 0.0374350192739342 |
| Pot1b | 364.688252068619 | -0.771812812863437 | 0.235106442531906 | -3.28282289737209 | 0.00102773210148371 | 0.0210080532509169 |
| Trex1 | 460.487286283694 | -0.77039675328997 | 0.220168481837135 | -3.49912370227385 | 0.000466789965455844 | 0.0125987971258956 |
| Atp6v0a1 | 2861.22395062264 | -0.769072832631286 | 0.188727493080836 | -4.07504396988877 | 4.60056536329081e-05 | 0.00250775915881342 |
| Rasa4 | 288.335683511649 | -0.768649448996145 | 0.187157108533644 | -4.10697437579813 | 4.00875681678001e-05 | 0.00225138261629261 |
| Alms1 | 220.790390288098 | -0.767563278218721 | 0.257042520529966 | -2.9861334873163 | 0.00282529285486806 | 0.0408197047166315 |
| E130310I04R | 94.2061604311557 | -0.764495952140601 | 0.277186119785678 | -2.75806000939627 | 0.00581455226385454 | 0.065575883543674 |
| Nav1 | 2030.18375797644 | -0.764224553273444 | 0.198317985070996 | -3.85353125184214 | 0.000116426394822199 | 0.00492266734001084 |
| Mapre2 | 439.586770803381 | -0.763608925553823 | 0.192755365407592 | -3.96154433335294 | 7.44665516544629e-05 | 0.00353885391680772 |
| Acsf3 | 117.629902668384 | -0.761635890542608 | 0.28842414195641 | -2.6406800948643 | 0.00827398045354085 | 0.0809169030378888 |
| Uri1 | 977.184198335952 | -0.760698247165373 | 0.132523128489332 | -5.74011688251542 | 9.46112284717387e-09 | 2.88157041654001e-06 |
| Tspan33 | 606.033451474531 | -0.758311913893274 | 0.248221176453082 | -3.05498477095732 | 0.00225071950442463 | 0.0354504261886713 |
| Cecr2 | 108.628940562348 | -0.758237523062404 | 0.29129520378564 | -2.60298663763918 | 0.00924155555559054 | 0.0867224453824192 |
| Phykpl | 166.729374739954 | -0.757388162614148 | 0.294157971589919 | -2.57476674359861 | 0.010030770772056 | 0.0913098865498548 |
| Krba1 | 177.461082036983 | -0.754847893236112 | 0.280350557252628 | -2.69251433146219 | 0.00709154876705928 | 0.0734246017594965 |
| Zc3h12d | 619.357309541488 | -0.751626917830703 | 0.265856411135813 | -2.82719124439972 | 0.00469582744556708 | 0.0572992073259378 |
| Gm20199 | 437.389078455017 | -0.750943974838278 | 0.20180147566007 | -3.72120160361575 | 0.000198277062412094 | 0.00722898666689262 |
| Crip1 | 5050.86284974805 | -0.749257313046467 | 0.223925824465307 | -3.3460067182316 | 0.000819844092904616 | 0.0183065588616452 |
| Mvk | 185.902577033963 | -0.745958262280753 | 0.276636332272299 | -2.6965303369714 | 0.0070066015079506 | 0.0729262278799348 |
| Mfhas1 | 1406.69166638657 | -0.745455184899582 | 0.231899037509803 | -3.21456782617336 | 0.00130641047101581 | 0.0244567078075687 |
| Fam84b | 673.876907654177 | -0.741820651387967 | 0.2088698897661 | -3.55159210462879 | 0.000382907951013387 | 0.0112050958296549 |
| Apbb2 | 774.730394519487 | -0.736717283139793 | 0.171618503103581 | -4.29276138538012 | 1.76464619628107e-05 | 0.00116641153695421 |
| Apol7a | 537.924737339915 | -0.73584334135588 | 0.209086573124382 | -3.51932374403659 | 0.00043264840629815 | 0.011938090019939 |
| Mbp | 1174.82446122191 | -0.734745137617753 | 0.167517169743107 | -4.38608853495144 | 1.15407162682598e-05 | 0.000833329642227592 |
| Mob3c | 141.565766356384 | -0.734519436403394 | 0.175704564863056 | -4.18042318351761 | 2.90967093325419e-05 | 0.00174149941575699 |
| Polr1b | 78.2921401246456 | -0.732996871298728 | 0.272690511785796 | -2.68801751296178 | 0.00718776267905169 | 0.0740073342509767 |
| Cx3cl1 | 590.234016457499 | -0.732309801238834 | 0.2893113038265 | -2.53121738263639 | 0.011366737787876 | 0.0985944806561477 |
| Zfp865 | 230.023090788598 | -0.727579862978567 | 0.216629570805041 | -3.35863594372056 | 0.000783281837966014 | 0.0178834957587322 |
| Uchl4 | 87.3874986145496 | -0.727475099356223 | 0.250125461594461 | -2.90844080694076 | 0.00363235933829713 | 0.0487063903277746 |
| Socs1 | 470.714977828952 | -0.726582967302295 | 0.202962146000674 | -3.57989399313842 | 0.000343733622827863 | 0.0105445953058485 |
| Tbc1d8 | 4504.41238323917 | -0.725925959644808 | 0.207330578617185 | -3.50129712889655 | 0.00046299932619561 | 0.0125268917452438 |
| Gm8221 | 5992.10218634785 | -0.723692632070313 | 0.176305965688268 | -4.10475408047109 | 4.04745207465311e-05 | 0.00225828033674065 |
| Samd1 | 434.644954010596 | -0.72360053408618 | 0.188957042067145 | -3.82944465138829 | 0.000128432764277024 | 0.00528952718059445 |
| Arc | 1575.49518420187 | -0.716835186283332 | 0.199419219074566 | -3.59461434865661 | 0.00032487243793824 | 0.0102339419543151 |
| Tbc1d32 | 129.956502723329 | -0.716657492264495 | 0.258358967686669 | -2.77388278286372 | 0.00553916325378146 | 0.0634351136787331 |
| Hspbap1 | 174.637432440859 | -0.71509330767117 | 0.181187471087047 | -3.94670394912472 | 7.92344303469022e-05 | 0.00371766609897702 |
| Rgs10 | 1320.12931868386 | -0.714279113233453 | 0.219299118731877 | -3.25709978847091 | 0.00112556856089499 | 0.0221527830038094 |
| Dgka | 2106.41076765203 | -0.713792080831625 | 0.18114071922612 | -3.9405390675334 | 8.12987084944354e-05 | 0.00377987476875325 |
| Znf41-ps | 154.238598550773 | -0.712755015793461 | 0.239918361823318 | -2.97082311823366 | 0.00297002787452172 | 0.0421797062128755 |
| Naip5 | 150.745426826493 | -0.712550553326907 | 0.237291151248483 | -3.00285345482921 | 0.00267461188566295 | 0.0392370503543166 |
| Vrk2 | 291.765946559112 | -0.71196246639942 | 0.149314715224854 | -4.76820027635769 | 1.85878952765106e-06 | 0.000219890846249786 |
| Relt | 339.249582346406 | -0.711745607717911 | 0.214342328465161 | -3.32060220122877 | 0.000898234650378055 | 0.0195467109827866 |
| Perm1 | 233.703249796598 | -0.711616046357372 | 0.239820169262018 | -2.96729023479209 | 0.00300437208958748 | 0.0424505942010328 |
| Polr3c | 991.825299179531 | -0.709278311339118 | 0.188514209027791 | -3.76246604962576 | 0.000168246153194565 | 0.00640718227234098 |
| Ksr1 | 349.711440130623 | -0.705333888086533 | 0.220656267403556 | -3.19652777773385 | 0.00139092433088482 | 0.0255654191065111 |
| Rpusd2 | 65.8712026519442 | -0.705220411295714 | 0.270580931177827 | -2.60631969971394 | 0.00915209936966622 | 0.086477215432091 |
| Fkbp5 | 158.623270064156 | -0.705131386011459 | 0.224785338280488 | -3.13691004673798 | 0.00170738502007408 | 0.0296195342015971 |
| Galnt12 | 570.772950956949 | -0.704510820747692 | 0.222484082834346 | -3.16656729673666 | 0.00154249650643091 | 0.0274880787684482 |
| Pdcd4 | 769.148452867595 | -0.703932792296599 | 0.1993632442649 | -3.53090558338458 | 0.000414139548996724 | 0.0117122489076641 |
| Zfp933 | 211.805474291471 | -0.703617258092933 | 0.214120195907595 | -3.28608543958452 | 0.00101590190382574 | 0.0208984458871468 |
| Igsf8 | 395.53255211575 | -0.699831240180692 | 0.179182407760416 | -3.90569168551654 | 9.39562448740706e-05 | 0.00414600572618915 |
| Vav2 | 234.894105303153 | -0.698048237819102 | 0.239472069239187 | -2.91494636529776 | 0.00355749674927528 | 0.0480672707800013 |
| Rars2 | 125.776526541749 | -0.697560581545792 | 0.212161931655587 | -3.28786873357741 | 0.00100948896107199 | 0.0208652736935325 |
| Pan2 | 254.509720077646 | -0.696850818535939 | 0.230635438831553 | -3.02143860486632 | 0.00251576652848562 | 0.037601241662312 |
| Dclre1c | 2153.91962464982 | -0.695653723981851 | 0.15189534974793 | -4.57982239177358 | 4.65370943907711e-06 | 0.000431243741354479 |
| Samd9l | 821.8472423129 | -0.695307610951646 | 0.210693763453024 | -3.30008634122039 | 0.00096655086973176 | 0.0202793314555041 |
| Uhrf1 | 300.106858558041 | -0.691333755351452 | 0.271636521466858 | -2.54506924038858 | 0.0109256071525641 | 0.0961176831776207 |
| Epsti1 | 498.761940699157 | -0.688399551816825 | 0.210749625143178 | -3.26643310207145 | 0.00108911537541601 | 0.0215881692952336 |
| Tarbp2 | 145.697285163748 | -0.688071602830277 | 0.192691942032831 | -3.5708374495133 | 0.00035584168739644 | 0.0107526075104577 |

| | | | | | | |
|---|---|---|---|---|---|---|
| Fut4 | 547.821634396159 | -0.687803941338679 | 0.207539348410977 | -3.3140893358529 | 0.000919421266003331 | 0.0197969130460683 |
| Dtx3l | 1984.21476195233 | -0.687329872898575 | 0.120460070541236 | -5.70587307321296 | 1.15748231406772e-08 | 3.30030854677771e-06 |
| Iffo2 | 768.434721734533 | -0.687211125730558 | 0.206784901532714 | -3.32331384272676 | 0.000889547728855202 | 0.0194080185250643 |
| Adora2a | 1296.02959502949 | -0.6848336490438 | 0.225339888301734 | -3.03911417638938 | 0.00237274910212011 | 0.0363930620904491 |
| Ube2l6 | 988.300286001129 | -0.68367883848925 | 0.16812977278575 | -4.06637579508581 | 4.77499154685687e-05 | 0.00256511623193471 |
| Napsa | 333.680937091687 | -0.679193228605504 | 0.268325252213816 | -2.53123111970202 | 0.0113662925923715 | 0.0985944806561477 |
| Htra2 | 256.791031971511 | -0.678960395379265 | 0.230583424280717 | -2.9445325373982 | 0.0032344293557207 | 0.0446238888779333 |
| Gpr18 | 137.762068771123 | -0.67854399363418 | 0.214020385029317 | -3.17046431601004 | 0.00152195524921558 | 0.0272530472967428 |
| Mid2 | 85.0633341413411 | -0.675917995436612 | 0.226368774428507 | -2.98591533723254 | 0.00282730906797259 | 0.0408197047166315 |
| Mylk | 3392.68213892923 | -0.674410426908249 | 0.233959423943139 | -2.88259568920873 | 0.00394413321625151 | 0.051233712314029 |
| Ino80c | 414.293101094208 | -0.673666717829324 | 0.171544843276073 | -3.92705898331883 | 8.59908987705203e-05 | 0.00388706826962677 |
| Orai1 | 493.657583879987 | -0.667980469587764 | 0.23819081431798 | -2.80439223275849 | 0.00504115369072803 | 0.0598677099666391 |
| Ccdc167 | 286.61816433464 | -0.667947306859292 | 0.245719357337073 | -2.71833409503434 | 0.00656115505133576 | 0.0704927963003417 |
| 3110052M02 | 209.272277607634 | -0.667715156075466 | 0.238663981828622 | -2.79772067389261 | 0.00514645981111677 | 0.0606235520123077 |
| Gcnt7 | 115.486055086586 | -0.667470854320753 | 0.243818430261307 | -2.73757342135869 | 0.00618943023109863 | 0.0676300314890438 |
| Ms4a6d | 400.655499900641 | -0.665862050619596 | 0.247247285090525 | -2.69310156581013 | 0.00707907003970295 | 0.0733637081467818 |
| Apol7c | 35337.1110159831 | -0.665771394123089 | 0.150067167869275 | -4.43648936390304 | 9.1437850786182e-06 | 0.000711041189330311 |
| Nxf1 | 2458.516253488 | -0.663763674743314 | 0.202696363342117 | -3.27466987467848 | 0.00105785515390956 | 0.0213817049328046 |
| Akr1e1 | 112.703461164546 | -0.662708092780072 | 0.236754832545644 | -2.79913227389902 | 0.00512401424832989 | 0.0604231584744734 |
| Psd | 408.063005267611 | -0.661047397834954 | 0.179907782680806 | -3.67436799000403 | 0.000238438881284832 | 0.00823428683194824 |
| Dock6 | 257.63966514992 | -0.659901327927007 | 0.154192067392198 | -4.27973591046355 | 1.8711520798552e-05 | 0.00120972157720871 |
| Cacnb3 | 18080.6525295947 | -0.659038076479191 | 0.201339161340863 | -3.27327317790628 | 0.00106309677038863 | 0.0214160074034812 |
| Ptch1 | 147.752601935371 | -0.657427881228548 | 0.232878662664651 | -2.82304902349622 | 0.00475692972217718 | 0.0578055574446157 |
| Rtel1 | 303.93774445399 | -0.651193416811932 | 0.223772825367326 | -2.91006477548375 | 0.00361353855570334 | 0.0485505231880367 |
| Rogdi | 5163.86301662107 | -0.650254803773538 | 0.249104659601899 | -2.61036788638449 | 0.00904448994186068 | 0.0862463100474476 |
| Adam23 | 5330.07665097477 | -0.649897251930954 | 0.171591531210206 | -3.78746694168028 | 0.000152190926256322 | 0.00593811614024666 |
| Nrp2 | 13528.6723220373 | -0.647279740707422 | 0.212401250965116 | -3.04743845794829 | 0.00230800781428507 | 0.0358951704823076 |
| Pus7 | 97.658105023576 | -0.645556076669629 | 0.217039959519356 | -2.97436508050978 | 0.00293595535870078 | 0.0418561840881444 |
| Parp11 | 198.674299911718 | -0.64234961949335 | 0.193168644940529 | -3.32533066994963 | 0.000883137262691816 | 0.0193316660652224 |
| Fam60a | 1904.86211198118 | -0.641920505406085 | 0.208394923846558 | -3.08030778081108 | 0.00206786783895862 | 0.0337660651530394 |
| Cep128 | 168.419627304751 | -0.641365426920603 | 0.175833020843362 | -3.64758237016217 | 0.000264719459495272 | 0.00881957726478636 |
| Arpin | 499.478005589988 | -0.641262696628928 | 0.186958026739711 | -3.42998216129931 | 0.000603620931896971 | 0.0148852235505246 |
| Zdhhc13 | 135.656520674753 | -0.638611308812486 | 0.23249050122718 | -2.74682752818559 | 0.0060174769355547 | 0.0669410986184691 |
| Tiam1 | 178.860674639206 | -0.637823542777661 | 0.238621203380403 | -2.67295417901678 | 0.00751865194985803 | 0.0758687928152643 |
| Peak1 | 1672.05057069316 | -0.634955423546198 | 0.221595397341937 | -2.86538182273894 | 0.00416506701894611 | 0.0523931507360642 |
| Chst12 | 268.773852639769 | -0.633625818387444 | 0.221289052611427 | -2.86334010160033 | 0.00419200336326038 | 0.0526724038411925 |
| Rcsd1 | 2942.23922448063 | -0.629418121593844 | 0.19425780888727 | -3.24011747686859 | 0.00119480463345013 | 0.0229865528096287 |
| Ikzf1 | 1429.3583787865 | -0.628272881896995 | 0.18744402895933 | -3.35178925349131 | 0.000802911141492271 | 0.0181759477386831 |
| Ncoa3 | 7080.89750493972 | -0.628140889243112 | 0.139640154724216 | -4.49828267867268 | 6.85045733488103e-06 | 0.000564274707880571 |
| Inpp5f | 395.452755597559 | -0.627280655529978 | 0.225933477322556 | -2.77639534859385 | 0.00549653267931595 | 0.0631419869772659 |
| Sdhaf1 | 303.730180676363 | -0.626222466454921 | 0.234389120094905 | -2.67172156370297 | 0.00754632357843585 | 0.0760789829485101 |
| Zdhhc3 | 615.055570317928 | -0.625609149840163 | 0.211310094640315 | -2.96062121833343 | 0.00307019276271298 | 0.043229815839944 |
| Plag1 | 136.119154244084 | -0.624767462489504 | 0.185530367805759 | -3.36746738487365 | 0.000758619990921525 | 0.0174294510310896 |
| Dock10 | 9840.9958176227 | -0.623661204247212 | 0.167832823529215 | -3.71596682420499 | 0.00020242814432037 | 0.00730844469104712 |
| Fhod1 | 600.021406224425 | -0.623124193161422 | 0.24576657673596 | -2.53543098267132 | 0.0112309056796928 | 0.0977211824398932 |
| Ptprs | 238.994696819349 | -0.621444570975482 | 0.223708365390679 | -2.77792280986097 | 0.00547076127910958 | 0.062910925981074 |
| H2-Q4 | 2343.18736569623 | -0.620150069611276 | 0.111944381246443 | -5.5398052381569 | 3.02808221822471e-08 | 7.48272761481306e-06 |
| Pdlim4 | 515.065440503415 | -0.619089838855377 | 0.230178260863634 | -2.68961037646448 | 0.00715354858342944 | 0.0737233181165295 |
| Zfand6 | 2603.05224508315 | -0.617124450795391 | 0.150381936982309 | -4.10371393785138 | 4.06570132977034e-05 | 0.00225828033674065 |
| Tmtc2 | 1404.48740400806 | -0.615494077750526 | 0.193470405117932 | -3.18133451664272 | 0.00146598248224682 | 0.0265933526958966 |
| Tango2 | 145.857948644335 | -0.613924565785844 | 0.178977771741978 | -3.43017213707913 | 0.000603198450292034 | 0.0148852235505246 |
| Lgals1 | 356.475906641182 | -0.612966612879488 | 0.137433898996625 | -4.46008311890024 | 8.19278729764777e-06 | 0.000655579437835637 |
| Ntmt1 | 706.571993560931 | -0.612862509405323 | 0.218754836041922 | -2.80159524924914 | 0.00508506275278244 | 0.0602192734940796 |
| Akap5 | 2430.00280307694 | -0.610476103743911 | 0.19843152080198 | -3.07650771045151 | 0.00209440885959394 | 0.0339007664027432 |
| Lonrf1 | 344.323574820874 | -0.609098638196479 | 0.172959771820578 | -3.52162026918223 | 0.000428918056422385 | 0.0118645989736739 |
| Igsf3 | 2749.68466895602 | -0.609094935299934 | 0.200330562481258 | -3.04044938403703 | 0.00236225393660656 | 0.0363826368075691 |
| Vsig10 | 868.454706290783 | -0.608934876086035 | 0.153150250961046 | -3.97606188866721 | 7.00658982781191e-05 | 0.00340232658887635 |
| Adam22 | 157.734074643601 | -0.605087330641054 | 0.217807363521182 | -2.77808482164658 | 0.00546803421629545 | 0.062910925981074 |
| Rabgap1l | 12977.5293705681 | -0.603299338826484 | 0.176789798921878 | -3.41252347423664 | 0.000643643832499032 | 0.0155591418439106 |
| Dcaf15 | 198.997801738666 | -0.602660080152516 | 0.199285161857214 | -3.02410914358148 | 0.00249366458345594 | 0.0374350192739342 |
| Agap1 | 1246.26883564092 | -0.602360683473878 | 0.191139026513538 | -3.15142697156728 | 0.00162474786978952 | 0.0284971550663399 |
| Kdm6a | 1701.04805641788 | -0.60060937304832 | 0.185142374222774 | -3.24404056915481 | 0.00117846960116998 | 0.0227510103559205 |
| Gphn | 494.363760046832 | -0.600569534052885 | 0.172138442418698 | -3.48887514964322 | 0.000485057647143894 | 0.012771936267561 |
| Pde4a | 217.377474328978 | -0.600319650851752 | 0.162729428460986 | -3.68906630183168 | 0.000225078562709762 | 0.00790189332218843 |
| Nol4l | 486.641996255291 | -0.599039671751405 | 0.213480273083842 | -2.80606569917652 | 0.00501504659546077 | 0.0597916146783812 |
| Tmppe | 334.945477039309 | -0.598442130439613 | 0.205025901505488 | -2.91886110996368 | 0.00351312712393536 | 0.0476996014873763 |
| Coro7 | 148.710217848545 | -0.597781337965867 | 0.228344292377324 | -2.61789481025465 | 0.00884740737029472 | 0.0849594492993177 |
| Psmb10 | 428.476858333482 | -0.59728879296651 | 0.16891616811122 | -3.53600723746726 | 0.000406223564284668 | 0.0117026063078899 |
| Tle3 | 2712.79738936022 | -0.596994191322409 | 0.211881908155575 | -2.81757983264934 | 0.00483870839914142 | 0.0582317536080187 |

| | | | | | | |
|---|---|---|---|---|---|---|
| Psme1 | 4273.9814372554 | -0.592341852664745 | 0.171309220145559 | -3.45773480354088 | 0.000544737318526061 | 0.013925239039103 |
| Sbno2 | 5116.09805123236 | -0.589730244290962 | 0.170971182133673 | -3.44929617337433 | 0.000562049869447356 | 0.0142693939457868 |
| D1Ertd622e | 248.803422548072 | -0.588304593334052 | 0.197171514602164 | -2.98372000905447 | 0.00284767225657671 | 0.0410182843175297 |
| Eml6 | 327.394376030227 | -0.588252637371685 | 0.189129228665033 | -3.11032113610288 | 0.00186884041503821 | 0.0314763302452612 |
| Atxn7l1 | 872.976521870228 | -0.586799482129822 | 0.166131257880861 | -3.5321437375177 | 0.000412205234277208 | 0.0117122489076641 |
| Tmem106a | 367.383888125391 | -0.585395811439859 | 0.166679152286382 | -3.51211176328791 | 0.000444561004593002 | 0.0122062182001832 |
| Cd80 | 2691.01464545885 | -0.583701999544091 | 0.211303391096068 | -2.76238822536792 | 0.00573802049414305 | 0.065108967239664 |
| Elk3 | 316.759200271841 | -0.580800608120992 | 0.208390862559662 | -2.78707329576271 | 0.00531864477165691 | 0.0618654078042101 |
| Ctsc | 1106.86569734841 | -0.580131249116454 | 0.149878857990036 | -3.87066766384769 | 0.000108537676856939 | 0.00469626057061931 |
| Ndnl2 | 481.596817179986 | -0.579939830007757 | 0.166318636357527 | -3.48692030375411 | 0.000488616939644884 | 0.0128079448976291 |
| Limd2 | 170.909069543663 | -0.575445629119125 | 0.190920735789171 | -3.01405516137637 | 0.00257780932061453 | 0.0381694269577012 |
| Lilra6 | 285.30235874404 | -0.573219231015008 | 0.155955473326663 | -3.67553134742728 | 0.000237354892829685 | 0.00822238756469188 |
| Ttyh3 | 4116.70745993918 | -0.571526786725045 | 0.218225026951633 | -2.61897910935632 | 0.00881933482828357 | 0.0848365080367762 |
| Hivep1 | 8111.10571489845 | -0.567909341929431 | 0.159221946106053 | -3.56677804673461 | 0.000361397326558259 | 0.0107740972421658 |
| Aida | 611.701994615864 | -0.565610480225891 | 0.171982197436889 | -3.28877342338557 | 0.00100624993284152 | 0.0208370563374259 |
| Tmem110 | 237.378891021024 | -0.565209110479155 | 0.157708348261403 | -3.58388834015506 | 0.000338516846019456 | 0.0104564092437121 |
| Fut11 | 254.889504661414 | -0.564672829710052 | 0.22199360463726 | -2.54364458216142 | 0.0109702645642001 | 0.0963580900109831 |
| Ndufb4 | 468.611180582947 | -0.561880421253712 | 0.217715983420873 | -2.58079545849201 | 0.00985729661428739 | 0.0904398831277853 |
| Poglut1 | 1464.91093672497 | -0.561412076803881 | 0.189143531986069 | -2.96818014821268 | 0.00299568700621453 | 0.0423817296553506 |
| Gatsl2 | 543.2944861509 | -0.559346662537699 | 0.20169470713537 | -2.77323421363897 | 0.00555021583656866 | 0.0634962964018966 |
| Zc3h7b | 1622.54932147522 | -0.558927402792976 | 0.194279987434712 | -2.87691702152701 | 0.00401581180171572 | 0.0516001615010989 |
| Pias3 | 355.161589093898 | -0.557775462905911 | 0.207324298589985 | -2.69035258625905 | 0.00713765619069005 | 0.0736659524473031 |
| Mrpl13 | 241.830949273392 | -0.556775941878493 | 0.1999472383366 | -2.78461431380809 | 0.0053591426749434 | 0.0621414666792186 |
| Trim24 | 1032.94338657221 | -0.556195006314912 | 0.133108409635947 | -4.17851139410432 | 2.93423299090071e-05 | 0.00174484870902759 |
| Fam206a | 184.164495152008 | -0.552396691651297 | 0.163874338639401 | -3.37085535317903 | 0.000749351921161975 | 0.017287953035936 |
| Lrrk1 | 5872.85593876005 | -0.551210032011208 | 0.1564719635672 | -3.52273991739409 | 0.000427110270667141 | 0.0118440553860813 |
| Dhx57 | 672.409547126452 | -0.550824631973956 | 0.17803449781162 | -3.09392077796512 | 0.00197530224537957 | 0.0324613320030173 |
| Sipa1l3 | 3965.89201847434 | -0.55057937550914 | 0.189436270137042 | -2.90640950178568 | 0.0036560263747637 | 0.0488641986627071 |
| Unc13d | 317.745470590493 | -0.549926566336756 | 0.167644642819989 | -3.28031100240553 | 0.00103692713135122 | 0.0211571187167441 |
| Ggta1 | 3618.059868281 | -0.547738784575586 | 0.191654968919427 | -2.85794199682795 | 0.00426398326701356 | 0.0534560247228757 |
| Tdrd7 | 1996.37096510766 | -0.54760602743844 | 0.138957005088476 | -3.94083066981596 | 8.119993234348e-05 | 0.00377987476875325 |
| Znfx1 | 5351.63940984228 | -0.545644053117235 | 0.134474034153527 | -4.05761645028275 | 4.95760990871996e-05 | 0.00265041452812337 |
| Zfp608 | 491.825652879314 | -0.544702583094445 | 0.182443184667907 | -2.98560115625006 | 0.00283021513817652 | 0.0408197047166315 |
| Mapkapk3 | 506.533478162197 | -0.543727162712696 | 0.203976614565609 | -2.66563480264943 | 0.00768431173333453 | 0.0770517529800788 |
| Cd40 | 6069.4102235475 | -0.543565957368229 | 0.190549152948599 | -2.85262856830887 | 0.00433592687696125 | 0.0540676720051698 |
| H2-M3 | 234.707128770954 | -0.5431940689899 | 0.187727120483543 | -2.89353007488078 | 0.00380937807496896 | 0.0500712579121215 |
| Ctsh | 1461.61771308929 | -0.540180508670733 | 0.154496351139211 | -3.49639654715208 | 0.000471587314335762 | 0.0126974598920428 |
| Rsrp1 | 1189.13407897907 | -0.540059835803324 | 0.0972977742705178 | -5.55058776886089 | 2.84710726061731e-08 | 7.36275179954988e-06 |
| Slc25a24 | 826.947129069192 | -0.53837316748783 | 0.150403433970502 | -3.57952709772185 | 0.000344216555397753 | 0.0105445953058485 |
| B2m | 11440.3251940381 | -0.536559979442355 | 0.117593008306421 | -4.56285613549575 | 5.04623943223165e-06 | 0.000459952315462425 |
| Israa | 267.057952856454 | -0.534166106493384 | 0.160005690242909 | -3.33841943797406 | 0.000842564588030943 | 0.018626875186688 |
| Aldh18a1 | 272.747037324987 | -0.533287052207084 | 0.21019455200566 | -2.5371116763898 | 0.0111771292601245 | 0.0974057032700508 |
| Zc3h7a | 1622.57322092267 | -0.533135058539441 | 0.172917147968747 | -3.08318211815406 | 0.00204799779949075 | 0.0334907875446134 |
| Tns3 | 235.36504848126 | -0.532919936733935 | 0.197231874233012 | -2.70199702155818 | 0.00689243798039917 | 0.072390210553334 |
| Idh1 | 780.355114914275 | -0.532522996513956 | 0.169067204565652 | -3.14977110955405 | 0.00163398424500577 | 0.0285690327114216 |
| Nmi | 481.008094740055 | -0.531908144708926 | 0.150733550364655 | -3.52879729444527 | 0.00041745275780277 | 0.0117122489076641 |
| Erap1 | 775.538964500225 | -0.531603535958733 | 0.131419118732186 | -4.04510044723454 | 5.23006945365347e-05 | 0.00275523986216331 |
| Arhgap30 | 5671.37432889686 | -0.53134485061184 | 0.170931788838167 | -3.1085198032702 | 0.00188027044527465 | 0.0314763302452612 |
| Psmb9 | 405.231308268734 | -0.530834590543313 | 0.16047993210772 | -3.30779421184573 | 0.000940338805714928 | 0.0200317385431992 |
| Rai1 | 572.355795190442 | -0.529495788827225 | 0.147633529473884 | -3.58655510515917 | 0.000335075281509508 | 0.0104079249452115 |
| Lmf2 | 384.142909627582 | -0.528488940971346 | 0.146610603610031 | -3.6047115826429 | 0.000312499774063338 | 0.00995701285840778 |
| Nans | 3065.93141761116 | -0.528430742898994 | 0.198734151838083 | -2.65898305858134 | 0.00783769040085854 | 0.0780959832056872 |
| Pik3r5 | 12953.6433296483 | -0.527818901221081 | 0.155504648198975 | -3.39423230967164 | 0.000688212854511718 | 0.0163175414545209 |
| Pirb | 2041.64578704946 | -0.527556867549752 | 0.156036376082537 | -3.3809864135187 | 0.000722261140595307 | 0.0168729913517223 |
| Icam1 | 5834.8430001468 | -0.525811881413094 | 0.127619455569234 | -4.12015455690328 | 3.7861833328957e-05 | 0.00214807952356123 |
| Zmym1 | 616.342700733231 | -0.525266915325076 | 0.181037265825467 | -2.90142978535409 | 0.00371464015964985 | 0.0494692198506663 |
| Gcnt1 | 568.41625004029 | -0.525038471961072 | 0.174812265157549 | -3.00344184367089 | 0.0026694456263673 | 0.0392129925564126 |
| Prkcd | 3948.69169975819 | -0.523065712566917 | 0.123075471729041 | -4.24995903097964 | 2.138096186187e-05 | 0.00135088804490906 |
| Bcor | 2316.89594657341 | -0.52286179558294 | 0.19435943895312 | -2.69017958890618 | 0.00714135762034108 | 0.0736659524473031 |
| Ankrd33b | 4170.32126773262 | -0.520509839101422 | 0.143492849757033 | -3.62742701105153 | 0.000286259659929653 | 0.00932154447170718 |
| Ncf1 | 20592.2379980697 | -0.519381333020851 | 0.175664027332999 | -2.95667440230254 | 0.00310976309733576 | 0.0436295747719396 |
| Cep68 | 340.97184483669 | -0.519201861810894 | 0.162102513974844 | -3.20292294721268 | 0.00136040374485027 | 0.0252549075838649 |
| Peli1 | 2250.90858422393 | -0.518836728108491 | 0.171177159392466 | -3.03099274430024 | 0.00243751105241808 | 0.0368777182352231 |
| Fam49a | 10725.4192439034 | -0.517528549580878 | 0.145605838670461 | -3.55431179344506 | 0.000378969642969716 | 0.0111485249466223 |
| Nup85 | 180.532524898668 | -0.515315461817809 | 0.20124201418885 | -2.56067533360218 | 0.0104468933715471 | 0.0936440502971347 |
| Gusb | 723.16837790922 | -0.513713276628339 | 0.173819853159706 | -2.95543499370201 | 0.00312228487151992 | 0.0437277175960976 |
| Leng8 | 1043.33108425352 | -0.511205257636846 | 0.148127720730053 | -3.4511113457856 | 0.000558283193550205 | 0.0142061993415979 |
| Ganc | 212.164611970756 | -0.510630962112497 | 0.167031531891014 | -3.05709321067401 | 0.00223494765738847 | 0.0353754936070496 |
| Deaf1 | 214.711936727486 | -0.506359338649306 | 0.186721952511488 | -2.71183613837882 | 0.00669116608320264 | 0.0714066860318746 |

| | | | | | | |
|---|---|---|---|---|---|---|
| Zfp946 | 258.402440283343 | -0.505750721526792 | 0.177809346522683 | -2.84434272673216 | 0.00445031500050278 | 0.055108577734511 |
| Psmb8 | 5071.86860374196 | -0.504367601494598 | 0.140816081985495 | -3.58174715830079 | 0.000341304032397389 | 0.0105132987264791 |
| Limk2 | 283.5296419177 | -0.504207441912335 | 0.194848569115138 | -2.58768870719494 | 0.00966222551945897 | 0.0892391592827107 |
| Acat1 | 329.024569711072 | -0.503182404303376 | 0.183362412591689 | -2.74419602791692 | 0.00606593033398509 | 0.0673185082973196 |
| Tap1 | 1655.4310398359 | -0.503087015230124 | 0.0950420414615085 | -5.29331028136502 | 1.20121852417596e-07 | 2.26229454578433e-05 |
| Herc6 | 1498.46176982149 | -0.50123760585744 | 0.13961821090991 | -3.59005893708857 | 0.000330603201759258 | 0.0103557960663745 |
| Fzr1 | 697.153952141789 | -0.499322689563835 | 0.138163318878774 | -3.61400329418798 | 0.00030150522726304 | 0.0096899945871821 |
| Mdm4 | 1397.1190506797 | -0.497721446915437 | 0.119047239560777 | -4.18087348141607 | 2.9039141543919e-05 | 0.00174149941575699 |
| Nabp1 | 2690.61802740333 | -0.49741587600575 | 0.154247134890037 | -3.22479815498913 | 0.00126061513681463 | 0.0237593903752182 |
| Aco1 | 1810.35970653686 | -0.496892542489661 | 0.163168385040696 | -3.0452746245281 | 0.00232467935028129 | 0.0360033904946071 |
| B3gnt5 | 1196.10086591776 | -0.496750697018112 | 0.184370846970373 | -2.69430175746786 | 0.00705362727928254 | 0.0732365409389559 |
| Snapc3 | 316.806552814812 | -0.495164391461286 | 0.17939322089622 | -2.76021796691939 | 0.00577628084342888 | 0.0653430752583206 |
| Nckap1l | 6074.86391892339 | -0.492477916147463 | 0.127486820397353 | -3.86297120449392 | 0.000112016216256672 | 0.0047977101325527 |
| Rgs19 | 459.808757830111 | -0.491697347945651 | 0.183563549142347 | -2.67862192817135 | 0.0073925811841824 | 0.0752108899982692 |
| Nsmaf | 521.769860398934 | -0.490668703607657 | 0.146420685957829 | -3.35108868257164 | 0.000804945199630756 | 0.0181759477386831 |
| Zfp677 | 217.691931736908 | -0.490662820820433 | 0.15952456720945 | -3.07578217827861 | 0.00209951161357759 | 0.0339181961806522 |
| Rcl1 | 650.463519664076 | -0.49043220635806 | 0.179085206281515 | -2.73854114776583 | 0.0061712437328524 | 0.0676300314890438 |
| Sh3bgrl3 | 2801.27449768562 | -0.490169513117316 | 0.182738490526502 | -2.68235505122676 | 0.00731058248285097 | 0.0747695705714894 |
| Rap2a | 2814.74242643012 | -0.488340841049214 | 0.184435951405757 | -2.64775298593965 | 0.00810287070116158 | 0.0798347090978684 |
| Zbtb40 | 359.380871342499 | -0.487528129312888 | 0.148028766558383 | -3.29346883479302 | 0.00098959311604113 | 0.0206459201695635 |
| Abtb2 | 2224.7312907756 | -0.486177164698939 | 0.192852874135329 | -2.52097443130548 | 0.0117030355548296 | 0.0999521930642901 |
| Psme2 | 5520.99861581839 | -0.485786749516785 | 0.149320407333431 | -3.25331786988786 | 0.00114065820710342 | 0.0223311958855458 |
| Gm13157 | 1071.38552785094 | -0.482702870675122 | 0.175944892867276 | -2.74348895730239 | 0.00607900927670899 | 0.0673963939750787 |
| Abhd16a | 247.997670618827 | -0.482240631737451 | 0.169834174593185 | -2.8394793503286 | 0.00451872196603946 | 0.0557075257897548 |
| Rbl1 | 662.109626238797 | -0.478597008150206 | 0.166741293326483 | -2.87029684490395 | 0.00410086618251316 | 0.051920407984837 |
| Vrk1 | 373.93569164207 | -0.47807382631065 | 0.146326234200312 | -3.2671778162226 | 0.00108625429199419 | 0.0215699066553133 |
| Daglb | 200.790246410219 | -0.477730342470305 | 0.182036470396941 | -2.62436610327912 | 0.00868104178772031 | 0.083868970181972 |
| Mtmr4 | 2623.16373958589 | -0.476633267778917 | 0.161778347622119 | -2.94621174455456 | 0.00321692100318261 | 0.0444755336749512 |
| Tbc1d22b | 231.60094090954 | -0.475727050457694 | 0.1791104159234 | -2.65605463537726 | 0.0079060804274612 | 0.078496084244079 |
| Traf5 | 854.183437703201 | -0.473000349930525 | 0.168040843419426 | -2.81479395309821 | 0.0048808516900787 | 0.0584860676655982 |
| Nsun4 | 236.4096997035 | -0.472566121823578 | 0.174200521484513 | -2.71277099400412 | 0.00667231998939867 | 0.0713424983481858 |
| Trim3 | 485.278921468903 | -0.467723324589156 | 0.139659440615594 | -3.34902762410845 | 0.000810957098127564 | 0.0182178645074313 |
| Fbxl5 | 917.558349605124 | -0.467297225108636 | 0.127396224778992 | -3.66806179633114 | 0.000244396122432474 | 0.00839360906580522 |
| 4931406H21I | 707.439218997849 | -0.466298396671628 | 0.146878066904539 | -3.17473130263004 | 0.00149975307454387 | 0.0270596591052187 |
| Prdm1 | 2779.2046449553 | -0.466231038607138 | 0.148527598249098 | -3.13901957685477 | 0.0016951412258348 | 0.029499171253964 |
| Sbf2 | 2921.93986636798 | -0.463858456918627 | 0.168911035077022 | -2.74617023516026 | 0.00602954678589544 | 0.0669815786804768 |
| Atxn1 | 2531.76948878119 | -0.46287317862817 | 0.131044420650591 | -3.53218531800255 | 0.000412140421565996 | 0.0117122489076641 |
| Synpo2 | 1580.27344311163 | -0.462042845549158 | 0.173251781441186 | -2.66688654919262 | 0.0076557510478189 | 0.0769728315115246 |
| Ifi30 | 2872.83536105965 | -0.45742124018061 | 0.111166424313398 | -4.1147427652351 | 3.87611427422256e-05 | 0.00218793861570329 |
| Sirt1 | 922.368194852495 | -0.45637982879641 | 0.158035558670658 | -2.88783000886207 | 0.00387909451019789 | 0.0506531572566505 |
| Lsp1 | 31005.311015419 | -0.456214132989152 | 0.174949978642331 | -2.60768327341062 | 0.00911572570657131 | 0.0864220425055964 |
| Cdk16 | 317.86630435403 | -0.456055852379926 | 0.148287328449143 | -3.07548768427868 | 0.0021015860763012 | 0.0339181961806522 |
| Arfrp1 | 560.061910139539 | -0.455976012524037 | 0.163857394972686 | -2.7827612699448 | 0.00538984491595756 | 0.0623744707406563 |
| Jrkl | 279.778586158579 | -0.454927245736214 | 0.175982095122089 | -2.58507688194419 | 0.00973572856712021 | 0.0895461552244638 |
| Stxbp6 | 954.852258809652 | -0.450842435863041 | 0.172627769469262 | -2.61164491234023 | 0.00901077904900491 | 0.0861563740541141 |
| Cebpg | 893.236066656794 | -0.447572522187733 | 0.107830397573255 | -4.15070826279457 | 3.31448020231031e-05 | 0.00191963645050472 |
| Gfod1 | 2489.31044962359 | -0.446856602851164 | 0.157022414276637 | -2.84581411456269 | 0.00442980454020764 | 0.054977038490077 |
| Rnf115 | 1951.81040360277 | -0.446106922303195 | 0.129443825184202 | -3.44633605865998 | 0.000568243230613974 | 0.0143610561918804 |
| Irf5 | 7132.34491203561 | -0.445445136332817 | 0.167484224893556 | -2.65962443099293 | 0.00782278280575701 | 0.0780422841010338 |
| Tank | 3999.57324907682 | -0.444656991623119 | 0.131507529931933 | -3.38122837417196 | 0.000721625396910672 | 0.0168729913517223 |
| Iscu | 4464.03614689249 | -0.44435303777957 | 0.162589547483142 | -2.73297419580827 | 0.00627652494272276 | 0.0682257647732914 |
| Bre | 420.166833460197 | -0.442118283801334 | 0.160350464878096 | -2.75719988799188 | 0.00582987014626887 | 0.0656009820830332 |
| Nuak2 | 2442.31830643773 | -0.440625840608158 | 0.156520073372835 | -2.81513949689108 | 0.00487560651176732 | 0.0584860676655982 |
| Atxn7 | 938.235037254146 | -0.439761235437423 | 0.153299609745582 | -2.86863897544982 | 0.00412242047594025 | 0.0520332754738428 |
| Heatr6 | 508.024165647069 | -0.438774513886569 | 0.119661652540185 | -3.66679303329211 | 0.000245611427683646 | 0.00840236814130427 |
| Rai14 | 2623.51633014018 | -0.438735453573294 | 0.17098547738122 | -2.56592232447387 | 0.0102901868732974 | 0.0927284262812535 |
| Wipf1 | 4636.84044626328 | -0.437497626828709 | 0.155783599965461 | -2.80836767750718 | 0.00497933406663023 | 0.0594738934703847 |
| Wrn | 790.341566074061 | -0.437264698966698 | 0.156839539780951 | -2.78797489190163 | 0.00530386543150217 | 0.0617580980086954 |
| Phldb1 | 2380.98266775085 | -0.435393111094577 | 0.166805034412515 | -2.610191668543 | 0.00904915056594093 | 0.0862463100474476 |
| Gsto1 | 1772.03960789124 | -0.434851113806497 | 0.168358982524318 | -2.58288038622285 | 0.00979792859906498 | 0.0899694186801012 |
| E030024N20I | 1024.98490054802 | -0.434129745422668 | 0.130418070827153 | -3.32875454045041 | 0.000872352500466552 | 0.019166945958035 |
| AW554918 | 198.330300204127 | -0.433219390357622 | 0.167990824577348 | -2.57882769161689 | 0.00991362227920274 | 0.0908816815702675 |
| Atic | 309.299495907514 | -0.432501241473187 | 0.156078389299146 | -2.77105141471083 | 0.00558756032471473 | 0.0638578322824541 |
| Ilf2 | 384.711828461968 | -0.431379784729078 | 0.141457517842784 | -3.04953594059605 | 0.00229195207728788 | 0.0358170719032573 |
| Gsap | 839.681729969075 | -0.430686392244703 | 0.14904192996391 | -2.88969951173468 | 0.00385610227341474 | 0.0505063101064451 |
| Sav1 | 2981.53546052713 | -0.426635642960796 | 0.143161166743343 | -2.98010733403466 | 0.00288147399204386 | 0.0412912252468141 |
| Ermard | 274.517500130172 | -0.425996355681971 | 0.140387607344106 | -3.03442991686442 | 0.00240990739832594 | 0.0366596036516888 |
| Tyk2 | 2042.88887348464 | -0.423737511021541 | 0.152674439659351 | -2.77543190574001 | 0.0055128442358928 | 0.0632640122839298 |
| Nedd4l | 1301.13882229436 | -0.423272238594487 | 0.118749446060323 | -3.56441442581104 | 0.000364669397584509 | 0.0107913177489103 |

| | | | | | | |
|---|---|---|---|---|---|---|
| Asah1 | 762.233655540028 | -0.422079788393229 | 0.143650109348889 | -2.93824898781041 | 0.00330071820830244 | 0.0453695753724637 |
| Mcmbp | 960.484489511333 | -0.420147551581854 | 0.153426354106771 | -2.73843143850938 | 0.00617330307956999 | 0.0676300314890438 |
| Zbtb46 | 1913.70310175867 | -0.419227368356833 | 0.141797733194737 | -2.9565167151233 | 0.0031113536685385 | 0.0436295747719396 |
| Map3k8 | 514.897122623639 | -0.416822648162637 | 0.161058776400719 | -2.58801573858707 | 0.00965305700497566 | 0.0892391592827107 |
| Zfp280b | 292.630006384287 | -0.416361026607445 | 0.144905712704516 | -2.8733237554027 | 0.00406177636720641 | 0.051920407984837 |
| Snrk | 378.837199504752 | -0.416289351677925 | 0.155714401720088 | -2.67341586314057 | 0.00750831080518885 | 0.0758687928152643 |
| Ptms | 6395.68555130979 | -0.413982835077724 | 0.148912062840252 | -2.78004902478472 | 0.00543506925690471 | 0.0626949897684443 |
| Arhgap26 | 5380.68868741738 | -0.412297325330672 | 0.144495132646481 | -2.85336480045587 | 0.00432589307836053 | 0.0540493607094034 |
| Cd47 | 2085.21557024152 | -0.406877152428238 | 0.142747980847863 | -2.85031809214786 | 0.00436755254496757 | 0.054325709507874 |
| H2-K1 | 10162.1281379053 | -0.404953810943051 | 0.0946095804869843 | -4.28026219816884 | 1.8667325597701e-05 | 0.00120972157720871 |
| Psme2b | 2821.76616309071 | -0.404393872948675 | 0.116021391148887 | -3.48551132635298 | 0.000491197428057483 | 0.0128218671361484 |
| Cers6 | 2428.04150800968 | -0.404237109606763 | 0.152158930708256 | -2.65667685573996 | 0.00789150461362004 | 0.0784213863301652 |
| Il10ra | 2174.06909271899 | -0.40419016869622 | 0.146892488583154 | -2.75160542649131 | 0.00593039274924357 | 0.066544871212501 |
| Suclg1 | 382.358526535995 | -0.402495469586498 | 0.158717652448371 | -2.53592126255411 | 0.0112151947218647 | 0.0976608968732462 |
| Rbck1 | 694.12589833885 | -0.400133089863544 | 0.156444557856127 | -2.5576670441392 | 0.0105376933514607 | 0.0942293870864052 |
| Nfkb2 | 7989.55318171625 | -0.398306439876962 | 0.157889501937983 | -2.52269109084537 | 0.0116460655657673 | 0.0996186531471791 |
| S100a10 | 3366.16214261712 | -0.394785327124025 | 0.137520798415055 | -2.87073178511162 | 0.00409522839865072 | 0.051920407984837 |
| Csrp2bp | 381.606881258924 | -0.393802072499839 | 0.143960149529689 | -2.73549363338654 | 0.00622867912153203 | 0.0678383073765291 |
| Brip1os | 367.642134440241 | -0.393379548806376 | 0.128128689058405 | -3.07019100638 | 0.00213921913129369 | 0.034227506100699 |
| Mapkap1 | 1420.14522544257 | -0.392899045440656 | 0.146845944201144 | -2.67558663317578 | 0.00745985904678844 | 0.0756186258890497 |
| H2-Q5 | 912.774612863973 | -0.39171914896961 | 0.119732151906186 | -3.27162873742162 | 0.00106929893015359 | 0.0214245118978521 |
| Nhlrc2 | 3463.4472508059 | -0.391552027582896 | 0.127521729473603 | -3.07047300251639 | 0.00213720006992071 | 0.034227506100699 |
| Colgalt1 | 385.93022137884 | -0.391403885047242 | 0.130921827983644 | -2.98959990916214 | 0.00279343075900999 | 0.0405521540994662 |
| Slc30a7 | 870.797978042275 | -0.391181989481643 | 0.144509502696617 | -2.70696377872734 | 0.00679016654141348 | 0.0717743839738763 |
| 1810026B05F | 660.412763500677 | -0.390878851393232 | 0.132474972307796 | -2.95058639819945 | 0.00317171323798414 | 0.0441419914973512 |
| Ldb1 | 315.987454620945 | -0.389911714206612 | 0.144543799922143 | -2.69753330420698 | 0.0069855296856775 | 0.0729262278799348 |
| Nrde2 | 578.914868594892 | -0.389103493047356 | 0.148740282924596 | -2.61599269139895 | 0.00889684619113377 | 0.0852135483595241 |
| Zfp592 | 891.030246925137 | -0.382825861578682 | 0.123710413041803 | -3.09453223997661 | 0.00197123509433116 | 0.0324613320030173 |
| Dap3 | 324.614699105191 | -0.382402308962236 | 0.14181588836866 | -2.69647014422075 | 0.00700786794033135 | 0.0729262278799348 |
| Epb4.1 | 3612.63092782094 | -0.381872965055209 | 0.14133382895175 | -2.70192188160117 | 0.00689399577122129 | 0.072390210553334 |
| Mlxip | 1912.91105920774 | -0.379820712346151 | 0.134277391046872 | -2.8286274359737 | 0.0046748085088605 | 0.0571879764780295 |
| Dph5 | 526.002818143516 | -0.376866922814617 | 0.129092283927076 | -2.91936056400946 | 0.00350750269016684 | 0.0476814546633927 |
| Zfp598 | 1063.70617797496 | -0.376234261939094 | 0.125807201638067 | -2.99056220184818 | 0.00278464409387789 | 0.0405128746155246 |
| Zfp942 | 264.690277402477 | -0.372161158368547 | 0.138175138903441 | -2.69340173146936 | 0.00707269913828741 | 0.0733637081467818 |
| Atr | 561.232680526761 | -0.369437600163548 | 0.129881807666737 | -2.84441375432259 | 0.00444932293559205 | 0.055108577734511 |
| AW549877 | 516.998590690285 | -0.368932598509568 | 0.128248293745861 | -2.8767057068272 | 0.00401850178596687 | 0.0516001615010989 |
| Ccdc71 | 421.857792541765 | -0.366004163549023 | 0.136594393322512 | -2.67949624172972 | 0.00737330309592422 | 0.075152273534993 |
| Nisch | 1369.63223137627 | -0.365796104055981 | 0.121083312877986 | -3.02102821075427 | 0.00251917887778806 | 0.0376017035181251 |
| Zfp687 | 433.334337802641 | -0.364061165314185 | 0.140415524901466 | -2.59274154741548 | 0.00952142934095905 | 0.0887496179978748 |
| Fam13b | 1893.46288544602 | -0.361228594361668 | 0.10064933849951 | -3.58898130625494 | 0.00033197264204855 | 0.0103694825269098 |
| Cc2d1a | 449.321904204952 | -0.360619831584904 | 0.137085021147844 | -2.63062899626343 | 0.00852270186932906 | 0.0826984684004705 |
| Ccnl2 | 1020.57121643025 | -0.360281533534472 | 0.127748539183888 | -2.82023994823036 | 0.00479877515903139 | 0.057939608869087 |
| Plcl2 | 2453.7665461716 | -0.355311382937836 | 0.0975013939417041 | -3.6441672121147 | 0.000268258983216519 | 0.00890459669661999 |
| Zfp568 | 1051.19144289488 | -0.355032733450478 | 0.12577914733628 | -2.82266767559864 | 0.00476259103681985 | 0.0578055574446157 |
| Tex10 | 510.099556251833 | -0.352181214259591 | 0.124830426599712 | -2.82127702237945 | 0.00478328773329259 | 0.0578783020611683 |
| Pcif1 | 652.254983710809 | -0.349909024446615 | 0.127245778596664 | -2.74986744790753 | 0.00596193765101033 | 0.0666435371140802 |
| Vwa5a | 15080.9962676003 | -0.348478580137545 | 0.101216861867588 | -3.44289057877952 | 0.000575532126185492 | 0.0144142280251862 |
| Dennd4a | 17501.8630781198 | -0.348256633179117 | 0.12738533795534 | -2.73388318286059 | 0.00625922464829128 | 0.0681042838444218 |
| Tle4 | 602.983401375859 | -0.347070921169545 | 0.135509678518055 | -2.56122606861104 | 0.0104303459840092 | 0.0936440502971347 |
| Fkbp4 | 757.126621053331 | -0.347032192585094 | 0.120166676267524 | -2.88792370201289 | 0.00387793925950146 | 0.0506531572566505 |
| Sart3 | 1426.55068121582 | -0.34434635675214 | 0.121776410825286 | -2.82769342944568 | 0.00468846816406664 | 0.0572920505323308 |
| Cdyl | 1127.385542071 | -0.336088425939212 | 0.123613838977813 | -2.7188576029868 | 0.00655078025696384 | 0.0704927963003417 |
| Trap1 | 548.738410007015 | -0.333527015346577 | 0.116382702890129 | -2.86577822188442 | 0.00415985558050918 | 0.0523868562347249 |
| Zfp398 | 663.504981977364 | -0.332919741247932 | 0.119991180618338 | -2.77453509109862 | 0.00552806699110076 | 0.0633733040629283 |
| Clip2 | 1604.98593852511 | -0.332455855570183 | 0.127894240630273 | -2.59945916197488 | 0.00933707875412711 | 0.0874712011338614 |
| Fam193a | 1702.47369715747 | -0.329093513296981 | 0.119158545486572 | -2.76181210464731 | 0.00574815481027149 | 0.0651447988164746 |
| 2310035C23F | 1157.24404362252 | -0.325934362436199 | 0.100755303988307 | -3.23491021846378 | 0.00121680986408194 | 0.0231782510014922 |
| Nfya | 2429.97987514045 | -0.325095622345193 | 0.099163578506663 | -3.27837727561788 | 0.00104405751880335 | 0.0212247159215599 |
| Fnbp4 | 1308.62543817525 | -0.324667329549916 | 0.119206122955401 | -2.72357930532968 | 0.00645787090209646 | 0.0697199266323424 |
| Pkd1 | 982.280546811896 | -0.321394472454842 | 0.103397002601037 | -3.10835386297378 | 0.00188132661294147 | 0.0314763302452612 |
| Srsf7 | 1607.73233287676 | -0.319969507225049 | 0.102168400932806 | -3.13178540824464 | 0.00173746818553618 | 0.0300476613112945 |
| Zfp217 | 4150.40912769351 | -0.318536499537019 | 0.106883527806419 | -2.98022067641642 | 0.00288040797099905 | 0.0412912252468141 |
| Yeats4 | 614.29561376276 | -0.318434737254406 | 0.124789761286742 | -2.55176974433588 | 0.010717732349572 | 0.0951166669810377 |
| Tet2 | 3358.24734961493 | -0.318167755263436 | 0.103000182867918 | -3.08900184838931 | 0.00200830189582291 | 0.0329385207692489 |
| Gm15800 | 818.50558548122 | -0.312919654443218 | 0.119983063380462 | -2.60803188072437 | 0.00910644725924513 | 0.0864220425055964 |
| Nfat5 | 13319.1350961892 | -0.309019193871618 | 0.12126112087953 | -2.54837817455622 | 0.0108225079412049 | 0.095740881707397 |
| Ppia | 13213.1347794366 | -0.308192208192408 | 0.118364517322922 | -2.60375503709104 | 0.00922086352462568 | 0.0867131310189391 |
| Tbc1d14 | 827.505979947371 | -0.306711848093604 | 0.119462914664805 | -2.56742311163419 | 0.0102457508811965 | 0.0924658952207219 |
| 2610005L07R | 1508.0361861151 | -0.306219830806702 | 0.11331025929891 | -2.7024898954551 | 0.00688222762734572 | 0.072390210553334 |

| | | | | | | |
|---|---|---|---|---|---|---|
| Gpbp1 | 5071.79203660304 | -0.305546983930024 | 0.0923928310796932 | -3.30704211960422 | 0.000942867139235344 | 0.0200471942414857 |
| Tap2 | 1415.48328741042 | -0.305264651794433 | 0.087131292308278 | -3.50350194180962 | 0.000459183307646604 | 0.0125111170194112 |
| Rnf169 | 1148.81407938283 | -0.30376115964096 | 0.100353997154076 | -3.02689646905231 | 0.00247078567464016 | 0.0373303487798893 |
| Fam208a | 2951.15853914647 | -0.30239018463289 | 0.112808374252182 | -2.68056504348605 | 0.00734979792902516 | 0.0749909056219329 |
| Kdm4c | 593.60435190594 | -0.298784047411031 | 0.11817188251021 | -2.52838527291139 | 0.0114588530158904 | 0.0990843277890364 |
| Nup88 | 2705.01925809712 | -0.298426670477577 | 0.110378532520838 | -2.70366586384211 | 0.00685792118954496 | 0.0722844394575734 |
| Prpf39 | 830.902394162559 | -0.298301503210385 | 0.110043544403747 | -2.71075877123626 | 0.00671294445859709 | 0.071448864041667 |
| Kidins220 | 2000.24995093742 | -0.292455890036221 | 0.0975706112575229 | -2.99737683578027 | 0.00272313869577819 | 0.0398438188119124 |
| Zufsp | 769.724930678937 | -0.286547805442391 | 0.111612560416146 | -2.56734371448877 | 0.0102480974220574 | 0.0924658952207219 |
| Tgif1 | 1214.38044402386 | -0.28186202307573 | 0.091001421161977 | -3.09733649735022 | 0.0019526808107227 | 0.032264205966176 |
| Ash1l | 2894.79091909689 | -0.272123466548054 | 0.101299354743863 | -2.68632971291992 | 0.00722417617895046 | 0.0742644073990883 |
| Cd63 | 4510.18381887828 | -0.264697010549527 | 0.100085213723502 | -2.64471644413714 | 0.00817593986568831 | 0.0803148863131219 |
| Jmjd1c | 2756.4005771194 | -0.260639521814113 | 0.0981038951782537 | -2.65677036921454 | 0.0078893160969121 | 0.0784213863301652 |
| Rap1b | 6741.77204447896 | -0.25006971372509 | 0.0867952219359323 | -2.88114608324498 | 0.00396231947752383 | 0.051233712314029 |
| Dhx15 | 4329.29718635712 | -0.238662262215898 | 0.0895087587694374 | -2.6663565163568 | 0.00766783299630921 | 0.0770246638834313 |
| Setd1b | 1895.92530298359 | -0.234785263761991 | 0.0876445564658351 | -2.67883452469194 | 0.00738788940295388 | 0.0752108899982692 |
| Dapp1 | 3148.1056005231 | -0.230119060704199 | 0.0911287278818299 | -2.52520874649543 | 0.0115629583176543 | 0.0995084095357014 |
| Gls | 2971.79639177906 | -0.222463842608235 | 0.0880577888633357 | -2.52633918566245 | 0.0115258142768407 | 0.0993543060143163 |
| H2afz | 5804.04056690441 | 0.198690953762569 | 0.0772604883217799 | 2.57170201843725 | 0.0101199953404303 | 0.0919372431451346 |
| Erbb2ip | 4111.92655748014 | 0.198741847967075 | 0.0718268066060638 | 2.76695926434653 | 0.00565818247838081 | 0.0643343447439617 |
| Prrc2c | 9488.91521567464 | 0.208260602223718 | 0.0824310153218251 | 2.52648352577767 | 0.0115210791718071 | 0.0993543060143163 |
| Ppp6r1 | 4247.11264314495 | 0.215080646222513 | 0.0844795102018598 | 2.5459504406286 | 0.0108980659395492 | 0.0961147781183443 |
| Supt6 | 6487.78761501287 | 0.22286188461856 | 0.0864869265790446 | 2.57682742853482 | 0.0099711718176814 | 0.0912587906276684 |
| Pstpip2 | 2768.80115424162 | 0.229227639781956 | 0.0880336814098066 | 2.60386293190302 | 0.00921796136596541 | 0.0867131310189391 |
| Smu1 | 1363.64632708746 | 0.233149055972624 | 0.0914312168134976 | 2.54999401843469 | 0.0107724767321007 | 0.095450152399171 |
| Tcf25 | 2446.64950631184 | 0.239884085763688 | 0.0928614360360337 | 2.58324764297856 | 0.00978750408621334 | 0.0899479714369359 |
| Ywhaz | 11729.7977764921 | 0.244903084162404 | 0.093250933473064 | 2.62628024236504 | 0.0086323713458328 | 0.0835439246002269 |
| Vim | 29998.2327623664 | 0.252837754002045 | 0.0934027088247187 | 2.70696382560515 | 0.00679016558261418 | 0.0717743839738763 |
| Capns1 | 1887.82575684692 | 0.252849385984557 | 0.0923225440883423 | 2.73876103048686 | 0.00616711818804962 | 0.0676300314890438 |
| Csf2rb2 | 26373.3803196829 | 0.253395632782486 | 0.0943309255486258 | 2.68624134989395 | 0.00722608712282496 | 0.0742644073990883 |
| Acap2 | 2257.99202082654 | 0.25517740759851 | 0.0945491896804427 | 2.69888518834438 | 0.00695721736376942 | 0.0727791694121504 |
| Kras | 1456.50970363713 | 0.25687269453924 | 0.0930178163268076 | 2.76154294610343 | 0.00575289500339731 | 0.0651447988164746 |
| Hectd1 | 4720.99510408186 | 0.260036950063249 | 0.0769893290768051 | 3.37757132295353 | 0.000731289856621696 | 0.0170124334845884 |
| Rps3 | 2280.08416264548 | 0.261346130144437 | 0.10274065687547 | 2.54374595308661 | 0.0109670816341925 | 0.0963580900109831 |
| Phf8 | 1082.31391081964 | 0.262420316485602 | 0.100918314007749 | 2.60032402508678 | 0.00931357730885814 | 0.0873246034354996 |
| Sec61a1 | 1196.48884810068 | 0.266952992755002 | 0.0947609940170634 | 2.817118958323 | 0.00484565745147556 | 0.0582317536080187 |
| Tpr | 8220.37092443616 | 0.267807267620124 | 0.0871558498281386 | 3.07274001857832 | 0.002121031860847 | 0.0341329584553092 |
| Txnrd1 | 12359.3458299637 | 0.269638251894389 | 0.10497380954599 | 2.56862405070913 | 0.0102103160411227 | 0.0923830060026729 |
| Rnps1 | 970.276946544259 | 0.271573114725074 | 0.10688382820414 | 2.54082511160051 | 0.0110591222319148 | 0.0969853621600099 |
| Smarca5 | 4137.09176697308 | 0.273635744458073 | 0.107748823208996 | 2.53957060790645 | 0.0110988639604393 | 0.0970971334928143 |
| Copa | 5080.91150723509 | 0.274828421355317 | 0.0930770835392368 | 2.95269695724256 | 0.003150110416668 | 0.0439974209636195 |
| Atp5f1 | 1305.68134783314 | 0.27597110774179 | 0.107164784434801 | 2.57520331139836 | 0.0100181179733123 | 0.0913098865498548 |
| Ndrg1 | 12521.1799265639 | 0.278280995543782 | 0.104937593829079 | 2.65187132074939 | 0.00800470421585396 | 0.0791220541158187 |
| Ctnnd1 | 2909.2130590841 | 0.281059892490221 | 0.0798793730851312 | 3.51855406014119 | 0.000433905400522893 | 0.0119431387470658 |
| Golga7 | 1700.59558307953 | 0.282698148144853 | 0.104543668918121 | 2.70411542918264 | 0.00684864938901733 | 0.0722552003850785 |
| Aamp | 2206.77162360884 | 0.282947970558596 | 0.0990218308899723 | 2.85743020519377 | 0.0042708654877226 | 0.0534820092606703 |
| Golga2 | 1976.13219989811 | 0.284836554444188 | 0.104073712383417 | 2.73687320189775 | 0.00620261954373219 | 0.0676870749031422 |
| Map1lc3b | 1418.37934067585 | 0.285166355779976 | 0.0877621128148816 | 3.24931051262955 | 0.00115685122785492 | 0.0224898350589978 |
| Nucks1 | 1814.25852998098 | 0.286594143728717 | 0.1114038212389 | 2.57257013755507 | 0.0100946499783066 | 0.0917845525419206 |
| Rbbp7 | 1430.47056817213 | 0.287265663110888 | 0.102863116111616 | 2.7926984323436 | 0.00522703970972638 | 0.0612574214906234 |
| Bap1 | 812.668184684941 | 0.292287512225579 | 0.0986492373389539 | 2.96289682627037 | 0.00304758700608456 | 0.043006557750838 |
| Ube2b | 837.972061386482 | 0.292310808724961 | 0.110719604910494 | 2.64009981756406 | 0.00828816120145282 | 0.0809169030378888 |
| Atp6v1g1 | 1445.09624267954 | 0.293107392295754 | 0.105814639055882 | 2.77000795835972 | 0.00560549230047539 | 0.0639313583397809 |
| Ppp1r12a | 9995.5099423878 | 0.293214838659062 | 0.105084975582347 | 2.79026413656338 | 0.00526650569742708 | 0.0615163270539802 |
| Rnf10 | 2314.82188254349 | 0.294541555791975 | 0.0988293325664336 | 2.98030501818863 | 0.00287961494352248 | 0.0412912252468141 |
| Cct8 | 2478.26858751553 | 0.2967338771131 | 0.114438938149806 | 2.59294504047794 | 0.00951579757874773 | 0.0887496179978748 |
| Bcl10 | 713.544968298126 | 0.299868119493639 | 0.118495942217785 | 2.53061930966807 | 0.0113861353551037 | 0.098685756156472 |
| Copb2 | 2831.66901430642 | 0.300981042147075 | 0.0847355073765938 | 3.55200613609843 | 0.000382305944961543 | 0.0112050958296549 |
| Gpalpp1 | 2009.8676438234 | 0.303725312515362 | 0.0997363842264116 | 3.04528096613042 | 0.0023246303298601 | 0.0360033904946071 |
| Capza1 | 4402.89339081247 | 0.304806687322419 | 0.104630870127578 | 2.91316211889248 | 0.00357788798120986 | 0.0481767005079158 |
| Rpl5 | 5295.81151385162 | 0.308676206760916 | 0.116003433691428 | 2.66092301700313 | 0.00779267717573811 | 0.0779267717573811 |
| Rab2a | 2000.53123365463 | 0.309815092786624 | 0.0983337208665226 | 3.15064954378331 | 0.00162907835060611 | 0.0285281122184881 |
| Sec31a | 1579.57681828544 | 0.31182734393302 | 0.105805388739293 | 2.94717828315315 | 0.0032068825360318 | 0.0444645059858773 |
| Mapk1 | 2537.94582630728 | 0.312338941757773 | 0.0894249052937382 | 3.49275116067295 | 0.000478071779320947 | 0.0127485807818919 |
| Apaf1 | 3967.41815167333 | 0.316093980709645 | 0.0803774131836253 | 3.93262196666515 | 8.40243226739688e-05 | 0.00382930519727268 |
| Adipor1 | 1307.77472908012 | 0.316663695649826 | 0.103419150013224 | 3.06194448135897 | 0.00219904235412177 | 0.0350334541229715 |
| Bnip2 | 2642.26484943277 | 0.318362634616194 | 0.119432351576479 | 2.66563146763739 | 0.007684387954578 | 0.0770517529800788 |
| Vapb | 683.965739033423 | 0.318447633598839 | 0.125688174941353 | 2.53363241010882 | 0.0112887082085661 | 0.098147330163608 |
| Trps1 | 1379.14751831577 | 0.321081367002957 | 0.0987987654851735 | 3.24985201410376 | 0.00115465077020937 | 0.022486368764848 |

| | | | | | | |
|---|---|---|---|---|---|---|
| Skap2 | 3298.26313393821 | 0.325210698383454 | 0.0952840880774736 | 3.41306407969221 | 0.000642368358567612 | 0.0155591418439106 |
| Vps26a | 2010.80608334403 | 0.325512617048373 | 0.112417876207091 | 2.89555921203073 | 0.00378483618384847 | 0.0499257157347509 |
| Pabpc1 | 12303.7285395659 | 0.325827076263779 | 0.115615916897251 | 2.81818528977585 | 0.00482959303421907 | 0.0581853461977422 |
| Atp5d | 1083.15918966833 | 0.326382774366222 | 0.113337184156604 | 2.87975016138782 | 0.00397990417131385 | 0.0513416872215894 |
| Prkar1a | 4570.51707508316 | 0.327008702742589 | 0.103072612268801 | 3.17260517168024 | 0.00151077828453319 | 0.027140314255265 |
| Ppp4r2 | 5237.69384958403 | 0.328676205308505 | 0.0909835144379197 | 3.61248086907842 | 0.000303281488009715 | 0.00971899177714131 |
| Ube2m | 1014.19160486819 | 0.329359756133664 | 0.101817622804452 | 3.23480107924168 | 0.00121727505259636 | 0.0231782510014922 |
| Wbp2 | 1083.8147865346 | 0.329526907770489 | 0.126395857173793 | 2.60710212453715 | 0.00913121218596801 | 0.0864220425055964 |
| Csde1 | 10442.4867637051 | 0.3318633579529 | 0.0855053081412981 | 3.88120182438841 | 0.000103941535787968 | 0.00451496046078988 |
| Arl8b | 1640.32702849287 | 0.331895825284736 | 0.125705809038705 | 2.6402584560157 | 0.00828428225555683 | 0.0809169030378888 |
| Stt3b | 884.497519215428 | 0.334127638449676 | 0.122036692648313 | 2.73792767731398 | 0.00618276710115591 | 0.0676300314890438 |
| Midn | 1548.72530119703 | 0.335321001388838 | 0.121627891153562 | 2.75694167027426 | 0.00583447583454316 | 0.0656009820830332 |
| Cul5 | 1008.35401035051 | 0.336108300554431 | 0.0949444679114966 | 3.54005144215182 | 0.000400049045900629 | 0.0116150010193603 |
| Csf2rb | 51584.2004096577 | 0.336110477181865 | 0.108380339383464 | 3.10121262854383 | 0.0019272984252942 | 0.0319397294922079 |
| Prpf6 | 1577.49482311598 | 0.336901326672472 | 0.0914969198999354 | 3.68210566039732 | 0.00023131549815729 | 0.00806341172259894 |
| Dhx8 | 1384.1171141998 | 0.337172719409754 | 0.121032159060724 | 2.78581099458523 | 0.00533939943265394 | 0.0620419244421232 |
| Tmem50a | 717.151469880392 | 0.337626848154434 | 0.127771216314375 | 2.64243276297629 | 0.00823128067344881 | 0.0806447939107936 |
| Wapal | 5644.53068201002 | 0.338806167725841 | 0.12593781912502 | 2.69026548244022 | 0.00713951963774189 | 0.0736659524473031 |
| Ssfa2 | 7973.40441267185 | 0.338995317996825 | 0.121065720445628 | 2.80009334392117 | 0.00510878313537446 | 0.0604046129010099 |
| Pafah1b2 | 1604.50627385046 | 0.342937861078728 | 0.132042240295814 | 2.59718299470264 | 0.0093991834722537 | 0.0879048950474862 |
| Ik | 3805.59629414769 | 0.343150296018041 | 0.11000300107319 | 3.11946303891953 | 0.00181180990310821 | 0.0309008069364467 |
| Ppig | 3092.45198861546 | 0.344183922123655 | 0.126796055978298 | 2.71446867544969 | 0.00663821779099758 | 0.0710461807852676 |
| Eif2ak1 | 1589.66110879053 | 0.345072212810891 | 0.131045386279597 | 2.63322672096711 | 0.00845778682437542 | 0.0822651503642702 |
| Samd4b | 1931.35138569983 | 0.34518289541948 | 0.102445198444355 | 3.36943947262667 | 0.000753212316645957 | 0.0173410371865487 |
| Vps4b | 2348.79068700937 | 0.347386498793369 | 0.120569636710353 | 2.88121046286224 | 0.00396151017978063 | 0.051233712314029 |
| Rab5b | 2362.59929276785 | 0.347689678892387 | 0.120127767208258 | 2.8943323177698 | 0.0037996579217974 | 0.050002598923535 |
| Chmp5 | 1234.69272390401 | 0.34877122798993 | 0.13237667271577 | 2.63468797662552 | 0.00842146586735442 | 0.0820742335188266 |
| Smap1 | 840.119287851087 | 0.349588809452092 | 0.11501732174009 | 3.03944487806867 | 0.00237014570972861 | 0.0363930620904491 |
| Akirin1 | 948.635973899987 | 0.349673875091199 | 0.108325084007088 | 3.22800465188948 | 0.00124656946935567 | 0.0235345543280732 |
| Lrp10 | 1563.74330483926 | 0.350350007834232 | 0.113843964135568 | 3.07745790911697 | 0.00208774320312111 | 0.0338915392973821 |
| Pde8a | 998.899562238906 | 0.350504845799381 | 0.13391837897743 | 2.61730203483463 | 0.00886278807205432 | 0.0849605201390034 |
| Derl1 | 802.984514312376 | 0.35057310925728 | 0.121009868836557 | 2.89706213739298 | 0.00376675141091411 | 0.0497461706524524 |
| Map7d1 | 3072.0300202338 | 0.351149829038552 | 0.106897722732091 | 3.28491402869834 | 0.00102013493845697 | 0.0208984458871468 |
| Tjp1 | 2792.09505887596 | 0.3518235610223 | 0.113115125493027 | 3.11031402289332 | 0.00186888542481646 | 0.0314763302452612 |
| Sucla2 | 702.960222611536 | 0.352155028397217 | 0.136168700608378 | 2.58616720893898 | 0.00970498375792017 | 0.0894854223781694 |
| Fam135a | 944.135949283916 | 0.352696303447478 | 0.107127291161635 | 3.29231048057888 | 0.000993678450274252 | 0.0206536530225228 |
| Nsf | 927.306980953303 | 0.353207042427928 | 0.121664749034783 | 2.90311733867095 | 0.00369468178911074 | 0.0492624238548099 |
| Cab39 | 2622.66529412093 | 0.354770988713775 | 0.105646489699515 | 3.35809537754481 | 0.000784815281498322 | 0.0178834957587322 |
| Sdc4 | 7011.73567832705 | 0.355380364942323 | 0.135090674346151 | 2.63068022024755 | 0.00852141752970329 | 0.0826984684004705 |
| Myo6 | 781.394136371591 | 0.356094611264965 | 0.132765788750136 | 2.68212628130528 | 0.00731558388325364 | 0.0747695705714894 |
| Glg1 | 3046.42055697833 | 0.357419200480939 | 0.110275215669568 | 3.24115621366745 | 0.00119045928422077 | 0.0229426468640121 |
| Mia3 | 2223.07262606502 | 0.358513021031103 | 0.13243498745909 | 2.70708691041217 | 0.00678764852882671 | 0.0717743839738763 |
| Pld3 | 550.953933132735 | 0.359569725770175 | 0.140426901971088 | 2.56054730769611 | 0.0104507433829806 | 0.0936440502971347 |
| Abcc1 | 6861.05321844646 | 0.362676328467329 | 0.080086193298991 | 4.52857494566293 | 5.93828189363073e-06 | 0.000507951497362874 |
| Syap1 | 1229.71184243069 | 0.363597965524922 | 0.121389990366472 | 2.99528786868863 | 0.00274185942311387 | 0.0400308284812173 |
| Ghitm | 3845.06991561741 | 0.364628097074026 | 0.142286989659598 | 2.56262429858379 | 0.0103884394738788 | 0.0933867800723789 |
| Nol11 | 468.382521086665 | 0.365592892531657 | 0.133875051080665 | 2.73085156330863 | 0.00631709167028449 | 0.0685327408522571 |
| Soga1 | 3516.63741757669 | 0.366052444829933 | 0.114725913345865 | 3.19066925818576 | 0.00141943670777018 | 0.0259607503131652 |
| Ftl1 | 62767.6287757189 | 0.366424633014218 | 0.0976059160744037 | 3.75412318998059 | 0.000173949304060843 | 0.00650754181686748 |
| Acsl4 | 2791.99445343052 | 0.367732593896325 | 0.121121027005226 | 3.03607559305504 | 0.00239679266122118 | 0.0365372098580231 |
| Ppfibp2 | 3103.46722244531 | 0.368256877061278 | 0.109901602327115 | 3.35078715199426 | 0.000805822143450608 | 0.0181759477386831 |
| Ppard | 978.733032010291 | 0.369151152678479 | 0.137462130596125 | 2.68547527291772 | 0.00724267337252339 | 0.0743661384140906 |
| Ss18 | 1008.41889937811 | 0.369870482713471 | 0.134660125438286 | 2.74669640704428 | 0.00601988296928679 | 0.0669410986184691 |
| Anxa5 | 3102.53930882774 | 0.370447395718818 | 0.141349294733878 | 2.62079408614148 | 0.0087725230653658 | 0.0844592696855997 |
| Dnajb11 | 1327.22438501785 | 0.37064020951418 | 0.123947709068747 | 2.9902949582441 | 0.00278708175187737 | 0.0405128746155246 |
| Clptm1l | 982.114561507421 | 0.3712928938237 | 0.126824169823531 | 2.92761935158208 | 0.00341567936050065 | 0.0465747672253917 |
| Zfp91 | 1061.01583064438 | 0.372322635082337 | 0.118083046924163 | 3.15305748607126 | 0.0016156998584386 | 0.0284609738973186 |
| Osgin2 | 3327.22368839218 | 0.373483485081147 | 0.128482981068811 | 2.90687126010194 | 0.00365063407918196 | 0.0488508435144445 |
| Myo1e | 853.531823538282 | 0.373639403137252 | 0.146787754154533 | 2.54543987875105 | 0.0109140156172761 | 0.0961147781183443 |
| Ppfibp1 | 2964.07777896962 | 0.373861155695915 | 0.147245608683939 | 2.53903093638877 | 0.0111159993375213 | 0.097101266797515 |
| Gnas | 7364.12940223868 | 0.373943116909786 | 0.140907791763836 | 2.65381432941992 | 0.00795876059635917 | 0.0788782690120446 |
| Gpkow | 900.455560795351 | 0.376627816294006 | 0.133354266791553 | 2.82426520992174 | 0.00473891544388391 | 0.057655076297581 |
| Arhgef12 | 2960.39155378583 | 0.376636311510224 | 0.0979679007080109 | 3.8444869063059 | 0.000120804934780246 | 0.00505019125848247 |
| Slc6a6 | 18095.4121076449 | 0.377246165207012 | 0.147455824173852 | 2.55836734371533 | 0.0105164935675696 | 0.0941573337128614 |
| Ppp3ca | 2385.702525592 | 0.377277392949428 | 0.100965381030254 | 3.73670053140667 | 0.000186450761955578 | 0.00686533931439082 |
| Arcn1 | 2787.48728559682 | 0.379033094703993 | 0.138460917347883 | 2.73747351934462 | 0.00619131043667685 | 0.0676300314890438 |
| Ano6 | 2145.84833547448 | 0.379429053896487 | 0.134579983004531 | 2.81935727309248 | 0.00481199255708765 | 0.0580361792134649 |
| Chmp2b | 705.01794632115 | 0.379670522797854 | 0.150421454980224 | 2.52404500972132 | 0.0116013073761571 | 0.0995084095357014 |
| Rps24 | 2181.61189282238 | 0.379787738588947 | 0.116980417520949 | 3.24659243519055 | 0.00116795513769261 | 0.0226497579532223 |

| | | | | | | |
|---|---|---|---|---|---|---|
| Osbpl9 | 3068.04223833001 | 0.382282606638531 | 0.0815252020730842 | 4.68913411948159 | 2.74363526620163e-06 | 0.000285775408058701 |
| Tnip3 | 1231.67719248163 | 0.38246084595645 | 0.141003828817006 | 2.71241461430672 | 0.00667949874137593 | 0.0713506493795392 |
| Hacd3 | 2846.90408834833 | 0.382774476591425 | 0.0830943976647925 | 4.60650160959778 | 4.09499893044506e-06 | 0.000395968592230861 |
| Spty2d1 | 2080.03490442181 | 0.383516465300156 | 0.133102155132569 | 2.88136931305264 | 0.00395951396155149 | 0.051233712314029 |
| Rad54l2 | 1820.0646073252 | 0.384307949340738 | 0.126503537194174 | 3.0379225582511 | 0.0023821516400246 | 0.0364367623618618 |
| Larp4b | 3571.97125091969 | 0.384448443097908 | 0.0959215319908854 | 4.00794727855722 | 6.12487883841705e-05 | 0.0030818394879275 |
| Taf1 | 2359.75355674814 | 0.38556439670099 | 0.0908409877054459 | 4.24438798432258 | 2.1919082507352e-05 | 0.00137706326260878 |
| Mtfr1l | 497.7671810942 | 0.38656544713654 | 0.129637296865685 | 2.98189993530221 | 0.00286465613175139 | 0.0412095422834093 |
| Copb1 | 1919.31879709789 | 0.386570158357216 | 0.151164171751571 | 2.55728691447151 | 0.0105492167181067 | 0.0942293870864052 |
| Grpel1 | 541.663521384579 | 0.386930017939081 | 0.143517289754309 | 2.69605159490872 | 0.0070166797432904 | 0.0729262278799348 |
| Gramd3 | 1163.06498180849 | 0.38723498583897 | 0.128109474925074 | 3.0226881037913 | 0.00250540319855007 | 0.0374967477360387 |
| Smox | 9244.50057248333 | 0.389023011508542 | 0.121666688973517 | 3.19744882342626 | 0.00138649011178736 | 0.0255261093428401 |
| Dot1l | 1958.45741751681 | 0.391045351967476 | 0.122711646477918 | 3.18670120719019 | 0.00143905354471649 | 0.026223394629043 |
| Uqcc1 | 238.000604354435 | 0.391874314649924 | 0.150529088029679 | 2.60331288642803 | 0.00923276502935467 | 0.0867131310189391 |
| Hnrnph2 | 1923.23898126195 | 0.393840213904944 | 0.106840110221823 | 3.68625802694555 | 0.000227575618420904 | 0.00795799017874357 |
| Fam118b | 227.703345664264 | 0.3945582284264 | 0.150151799445272 | 2.62772893754236 | 0.00859569796558071 | 0.0833340552548017 |
| mars-06 | 1541.7484274454 | 0.395374948650177 | 0.134193816586132 | 2.9462978154169 | 0.00321602591141015 | 0.0444755336749512 |
| Gtpbp4 | 3720.57141236758 | 0.396550097303194 | 0.121191176137216 | 3.27210371202445 | 0.00106750409034071 | 0.0214245118978521 |
| Mpp6 | 814.516789046649 | 0.397216074444751 | 0.157357042000447 | 2.52429805107561 | 0.011592959220498 | 0.0995084095357014 |
| Paf1 | 5694.20368312116 | 0.397269072593378 | 0.156051715793111 | 2.5457526729156 | 0.0109042416365889 | 0.0961147781183443 |
| Stxbp5 | 2348.28585036555 | 0.397859592189019 | 0.102991313053184 | 3.86304029334558 | 0.000111984527859232 | 0.0047977101325527 |
| Aldh9a1 | 877.868160991396 | 0.398818994981712 | 0.151803313922233 | 2.6272087524125 | 0.0086088502730402 | 0.0833888632719574 |
| Tmem50b | 1045.41439321035 | 0.399284680005883 | 0.155441808838966 | 2.56870839954991 | 0.0102078313542943 | 0.0923830060026729 |
| Atxn3 | 661.253188366808 | 0.399390927531589 | 0.142914035261938 | 2.79462354274422 | 0.00519601830511698 | 0.0610134356419227 |
| Cdc34 | 389.656373186674 | 0.401777820699788 | 0.143303061576705 | 2.8036932098951 | 0.00505209520765313 | 0.0598926425470179 |
| Slc38a1 | 1254.81174994639 | 0.40347473554783 | 0.134157509714907 | 3.00747037124675 | 0.00263431802818094 | 0.0388509502299365 |
| Gab2 | 3995.68999127472 | 0.406511246441349 | 0.124140286035767 | 3.27461180751771 | 0.00105807259498444 | 0.0213817049328046 |
| Pitpna | 2540.61019577609 | 0.406674814722544 | 0.143645493829208 | 2.83110039780344 | 0.00463881576453346 | 0.0568101666317314 |
| Slc39a6 | 694.885388959707 | 0.407233417224216 | 0.132970019250728 | 3.06259576044986 | 0.00219426255791943 | 0.0350074600345252 |
| sept-11 | 1409.08188868348 | 0.40747869055737 | 0.157377643384572 | 2.58917773702866 | 0.00962054241116567 | 0.0891503596768019 |
| Gtf2f2 | 675.632839705869 | 0.407643678464782 | 0.157683452825016 | 2.58520263960196 | 0.00973217805602477 | 0.0895461552244638 |
| Gpx1 | 3731.76630926393 | 0.408276986130702 | 0.139964370826574 | 2.91700654759194 | 0.00353408353964621 | 0.0479256206839828 |
| Cltc | 16034.9532745332 | 0.40874620998764 | 0.126358579870621 | 3.2348116796355 | 0.00121722986290814 | 0.0231782510014922 |
| Tmem167 | 851.409664556709 | 0.409458678982454 | 0.15550988287752 | 2.63300744239481 | 0.0084632492820796 | 0.0822651503642702 |
| Idi1 | 5766.82971999075 | 0.409733197316454 | 0.151939591843396 | 2.69668486235481 | 0.00700335129422111 | 0.0729262278799348 |
| Rplp2 | 4486.49218552002 | 0.410266702707506 | 0.130131010678169 | 3.15272048199295 | 0.00161756614236559 | 0.0284609738973186 |
| Rusc2 | 1338.47852934656 | 0.410528670835314 | 0.121289419898869 | 3.38470306130257 | 0.000712552921974953 | 0.0167164314184841 |
| Ccdc43 | 329.809703100215 | 0.410620622981279 | 0.160830347734148 | 2.55312898819341 | 0.0106759950991629 | 0.0949736524021528 |
| Cux1 | 1509.94733490137 | 0.412167821371096 | 0.125768246156238 | 3.27720099443124 | 0.00104841708085928 | 0.0212744487940788 |
| Calu | 1379.84908305938 | 0.412206701676498 | 0.127313844932071 | 3.23772093990588 | 0.00120488603344522 | 0.0231005736067428 |
| Chmp7 | 528.550383822749 | 0.41240442638043 | 0.155828226171078 | 2.64653225229983 | 0.00813217500220338 | 0.0800263593137183 |
| Entpd7 | 692.549141724779 | 0.412462917747523 | 0.139939082220457 | 2.9474462116147 | 0.00320410488966369 | 0.0444645059858773 |
| Ppp3r1 | 4610.4001966427 | 0.41269193232108 | 0.123863946596878 | 3.33181642971711 | 0.000862811490991039 | 0.0189989381778621 |
| Rheb | 1109.32560137337 | 0.413673399683062 | 0.124902845570487 | 3.3119613712052 | 0.000926443372687345 | 0.0198881280005469 |
| Usp9x | 6705.04296330823 | 0.414823638248277 | 0.105406575141668 | 3.93546263779796 | 8.30365821916795e-05 | 0.00379986334967686 |
| Naga | 557.00497248746 | 0.414965898516641 | 0.136292742411798 | 3.04466614416529 | 0.00232938729333945 | 0.036026128931759 |
| Rrn3 | 628.316460143742 | 0.415825690038246 | 0.157320819601981 | 2.64317012262126 | 0.00821337558742146 | 0.0805694474606014 |
| Smarca5-ps | 1207.31903373711 | 0.416500159902005 | 0.141300002533958 | 2.94763023660887 | 0.00320219835086056 | 0.0444645059858773 |
| Htatsf1 | 953.576694862224 | 0.417515054140867 | 0.143566045896838 | 2.90817408484511 | 0.00363545899029253 | 0.0487063903277746 |
| Rc3h2 | 2052.74851736394 | 0.418140390074079 | 0.13355733630329 | 3.13079312337093 | 0.00174334922198182 | 0.0301025517832885 |
| Trp53inp2 | 1400.47226197882 | 0.418540639530753 | 0.12792671971803 | 3.27172181427992 | 0.00106894699041644 | 0.0214245118978521 |
| Slc37a2 | 1002.88174599686 | 0.419345617543623 | 0.137903394836201 | 3.04086507835222 | 0.00235899513123163 | 0.0363826368075691 |
| Lamp2 | 2965.87410935638 | 0.419835204680977 | 0.165259860239062 | 2.54045479690985 | 0.0110708403494269 | 0.0970116191376098 |
| 2700029M09 | 527.020912321797 | 0.421957632827561 | 0.149601589816293 | 2.82054243772218 | 0.0047942531698915 | 0.057939608869087 |
| Nucb1 | 1438.59277860198 | 0.42209791386242 | 0.150121707826666 | 2.8117047159481 | 0.00492797203927582 | 0.058987135712322 |
| Sertad2 | 1058.44166173273 | 0.424237333311014 | 0.135833647553493 | 3.12321240688133 | 0.00178888596402467 | 0.0306513787403028 |
| Akirin2 | 754.698756743181 | 0.424798080669815 | 0.151487916801331 | 2.80417137973398 | 0.00504460829485079 | 0.0598677099666391 |
| Vamp3 | 973.976047813271 | 0.424977709190493 | 0.118273128746346 | 3.59318903368085 | 0.000326655430339367 | 0.010261040636649 |
| Rab31 | 941.632683390086 | 0.425084876127141 | 0.146661645409433 | 2.89840520294477 | 0.00375065672746366 | 0.0497107304045243 |
| Psme4 | 3820.80677898666 | 0.42571188395865 | 0.148474650728228 | 2.86723613674555 | 0.00414073930489602 | 0.0522052393088932 |
| Golgb1 | 4665.30888660136 | 0.426532606825343 | 0.0912731232804474 | 4.67314573551704 | 2.96621132653774e-06 | 0.000305409906954626 |
| Dad1 | 383.958092268023 | 0.427094748111714 | 0.138926664837947 | 3.07424603196157 | 0.00211035316139124 | 0.0340103292096676 |
| Clcn4-2 | 575.011165329644 | 0.428175693624148 | 0.14998946362355 | 2.85470514581479 | 0.00430768010870787 | 0.0538823428670771 |
| Cops2 | 1790.75703366943 | 0.428693608878885 | 0.145659787593355 | 2.94311570792407 | 0.00324926949978525 | 0.0447730815831622 |
| Pi4k2a | 1416.38788250182 | 0.429397720403449 | 0.116818948202156 | 3.67575403658294 | 0.000237147924407709 | 0.00822238756469188 |
| Tecpr1 | 1063.33412888958 | 0.429723470181831 | 0.16064415224002 | 2.67500225927786 | 0.00747287465881554 | 0.0756815721366382 |
| Flywch1 | 624.268103745389 | 0.431230237675149 | 0.16978933466083 | 2.53979579186509 | 0.0110917209741044 | 0.0970971334928143 |
| Atg5 | 571.084089225324 | 0.433204501918915 | 0.171410880645814 | 2.52728706769814 | 0.0114947503691776 | 0.0993175012472841 |
| Acbd3 | 3543.72834775838 | 0.433584201442191 | 0.170827556853862 | 2.53813968558427 | 0.0111443493369881 | 0.0972724997074632 |

| | | | | | | |
|---|---|---|---|---|---|---|
| Rpl23 | 2269.44828035576 | 0.433591558622764 | 0.130782163853046 | 3.31537226368247 | 0.000915211574758222 | 0.0197969130460683 |
| Hsd17b11 | 2148.81747957502 | 0.433597459763665 | 0.171976783916883 | 2.52125577585648 | 0.011693681798018 | 0.0999490711713758 |
| Hif1a | 3671.50052543267 | 0.434249470921336 | 0.11553775961099 | 3.75850693646329 | 0.000170930300842756 | 0.00646511886180764 |
| Gns | 1351.65937640683 | 0.434998952203936 | 0.167906834575392 | 2.59071617485955 | 0.00957764466112413 | 0.0889610028954468 |
| Hspe1 | 1759.54787202984 | 0.435139937722867 | 0.119556868077707 | 3.6396063623885 | 0.000273055134686693 | 0.00903682469558341 |
| Spcs2 | 475.756619204729 | 0.43530830341825 | 0.152138314084958 | 2.86126677580482 | 0.00421951829169405 | 0.0529582882659569 |
| Edem2 | 285.772576966311 | 0.436096266269441 | 0.162900324319951 | 2.67707426667186 | 0.00742681707070487 | 0.0753523775786844 |
| Dst | 5902.24224304625 | 0.436870506808065 | 0.155693763529248 | 2.80596021899102 | 0.0050166885337167 | 0.0597916146783812 |
| Picalm | 9122.57771583161 | 0.437821619875105 | 0.14711152704391 | 2.97612042151138 | 0.00291920209040038 | 0.0417243280787304 |
| 2010107E04F | 952.272939392593 | 0.440745680688449 | 0.170295360132975 | 2.58812501024275 | 0.0096499952399533 | 0.0892391592827107 |
| Pdha1 | 554.48028104287 | 0.440929951466656 | 0.124999847554548 | 3.5274439136755 | 0.000419592644831844 | 0.0117232919862565 |
| Crlf3 | 1878.22246579243 | 0.441042608149585 | 0.13309320462109 | 3.3137875776995 | 0.00092041403280731 | 0.0197969130460683 |
| Txnip | 1021.24378785699 | 0.443414165072833 | 0.152908520225269 | 2.89986564790231 | 0.0037332263828318 | 0.049538755819916 |
| Hspd1 | 3954.06056404199 | 0.443466359581796 | 0.127883112211963 | 3.46774763228129 | 0.000524839927656318 | 0.0135726046407866 |
| Srp14 | 1166.88826307029 | 0.443803307138763 | 0.155595847310895 | 2.85228246645942 | 0.00434065104060157 | 0.0540676720051698 |
| Mfsd1 | 2621.89766288794 | 0.44471926625825 | 0.145428191344611 | 3.0579990175662 | 0.00222820306569699 | 0.0353461028395871 |
| Ybx1 | 24116.1893219797 | 0.445662896873124 | 0.146183604166687 | 3.04865172406718 | 0.00229870803198451 | 0.0358170719032573 |
| Alg10b | 368.425930737906 | 0.445998100428666 | 0.145919285258357 | 3.05647125147992 | 0.00223958956063209 | 0.0353754936070496 |
| Atp2a2 | 1929.44786235681 | 0.446482905469352 | 0.175827691515147 | 2.53932074988818 | 0.011106794406732 | 0.0970971334928143 |
| Mfsd5 | 255.272499254001 | 0.44755830553768 | 0.147424314813199 | 3.03585135264003 | 0.00239857582701051 | 0.0365372098580231 |
| Ei24 | 536.182372864204 | 0.449060355469009 | 0.148500745991948 | 3.0239602667946 | 0.00249489202841027 | 0.0374350192739342 |
| Srp68 | 1209.99468350922 | 0.449085391193231 | 0.168859769207968 | 2.659516788988 | 0.00782528298315222 | 0.0780422841010338 |
| Stx11 | 5691.60645119266 | 0.450586283151094 | 0.127030614661484 | 3.54706843190386 | 0.000389543393479131 | 0.0113395877892354 |
| Uso1 | 2750.78630140363 | 0.450941669861375 | 0.163452380115017 | 2.75885655225123 | 0.00580039902190968 | 0.0654826772828788 |
| Mtss1 | 13163.9053257282 | 0.451471994958114 | 0.167349848675998 | 2.69777354763074 | 0.00698049075277349 | 0.0729262278799348 |
| Morf4l2 | 3605.37781694102 | 0.452402954648569 | 0.160603857901565 | 2.81688721902216 | 0.00484915502167494 | 0.0582317536080187 |
| Atp2b4 | 1818.78287274659 | 0.453231478278209 | 0.178245081322507 | 2.54274325504756 | 0.0109986013107432 | 0.0965307392071543 |
| Fam204a | 329.183282369441 | 0.45394820533318 | 0.176207278746449 | 2.57621710387107 | 0.00998879061377408 | 0.0913064628532722 |
| Ube4b | 2618.09308803957 | 0.454240654673035 | 0.13745294964226 | 3.30469921420572 | 0.000950783784564539 | 0.0201385060654432 |
| Cox5a | 357.496097377137 | 0.454663456417489 | 0.15607543995586 | 2.91310059126582 | 0.0035785930413254 | 0.0481767005079158 |
| Dcun1d2 | 217.791226384022 | 0.455300123952029 | 0.15047833724454 | 3.0256855058953 | 0.00248070179641656 | 0.0373785961736478 |
| Srp9 | 757.179164143964 | 0.457871563200323 | 0.14202862411792 | 3.22379778051058 | 0.00126502696886524 | 0.0238021994818638 |
| Slc9a3r1 | 1256.15358143153 | 0.459068334836408 | 0.169818014871621 | 2.70329584987467 | 0.00686556078817471 | 0.0722964355724458 |
| Ttc39c | 991.505551950308 | 0.462154368002 | 0.135749784191237 | 3.40445747855439 | 0.000662956209902841 | 0.0158198992577674 |
| Myo5a | 4178.32687960793 | 0.464621599164118 | 0.126879913397314 | 3.66190034910565 | 0.000250351247167635 | 0.00846171996505804 |
| Sppl2a | 5104.16118577318 | 0.466606734061493 | 0.149062296956358 | 3.13028004793261 | 0.00174639727177626 | 0.0301084304839566 |
| Rpn1 | 906.955850209177 | 0.467932766012069 | 0.170210458228305 | 2.74914227294087 | 0.00597514446967068 | 0.0666435371140802 |
| Mmp14 | 9626.14109005074 | 0.468013955657025 | 0.158867071121999 | 2.94594689983063 | 0.00321967667341149 | 0.0444755336749512 |
| Ufm1 | 886.061708468345 | 0.468090724184674 | 0.14487985506001 | 3.23088895961961 | 0.00123405871309011 | 0.0233777391645008 |
| Yipf5 | 378.686818107605 | 0.471963057368965 | 0.155650140277488 | 3.03220451024049 | 0.00242774660417299 | 0.0367800302975526 |
| Tctex1d2 | 357.006859776345 | 0.472308632785964 | 0.17117347221288 | 2.75923965717408 | 0.00579360295283755 | 0.0654724236133675 |
| Ciapin1 | 260.341859202964 | 0.47294496108613 | 0.161314095606292 | 2.93182662871825 | 0.00336974806071049 | 0.0461419548885938 |
| Arl4c | 16307.5643757898 | 0.47371611399493 | 0.112465985492304 | 4.21208343057061 | 2.53026048608518e-05 | 0.00158070205647569 |
| Tubb4b | 6658.58716015573 | 0.473876697914801 | 0.151863978514119 | 3.12040223462698 | 0.00180604237826795 | 0.0308497561387705 |
| Slco3a1 | 532.497839612048 | 0.474786548812848 | 0.180958178286183 | 2.62373634233861 | 0.00869710813129159 | 0.0839512521006619 |
| Grina | 538.628823392931 | 0.475771244870489 | 0.179608920016698 | 2.64892882172142 | 0.00807473360933615 | 0.0796726155597321 |
| St5 | 284.412926961102 | 0.476182581314782 | 0.158483641278953 | 3.00461661198607 | 0.00265915802379516 | 0.0391135413023839 |
| Aldoa | 7424.92594546779 | 0.476610708203031 | 0.142667211253147 | 3.34071651093914 | 0.000835624983984627 | 0.0185472052333514 |
| G6pdx | 1956.25926385459 | 0.476635887289891 | 0.136711707458499 | 3.48643065140987 | 0.000489512282508306 | 0.0128079448976291 |
| Aldh2 | 441.921967944424 | 0.476807605986001 | 0.169750269540959 | 2.80887687115544 | 0.00497146564431225 | 0.0594437612524217 |
| Cript | 305.39348002206 | 0.477249451380464 | 0.18227532489616 | 2.61828885315304 | 0.00883719636541828 | 0.084934851843951 |
| Dnajb9 | 696.27518740862 | 0.477261861672473 | 0.185622203217793 | 2.57114641136166 | 0.010136246411505 | 0.0919372431451346 |
| Manf | 476.190167583295 | 0.478095340047973 | 0.189449793151358 | 2.52359916627624 | 0.0116160292969921 | 0.0995084095357014 |
| Zfp451 | 924.480603973539 | 0.480001907111065 | 0.176618663681078 | 2.71773037518736 | 0.00657313781660239 | 0.0705069746220037 |
| Ptprc | 2865.22150844914 | 0.480400807781735 | 0.141094719794974 | 3.40481067243204 | 0.000662099401332332 | 0.0158198992577674 |
| Man1a | 640.44021486541 | 0.481636332319914 | 0.13279301398481 | 3.62697040956544 | 0.000286766142432912 | 0.00932154447170718 |
| Scpep1 | 2088.1099136586 | 0.481998160407185 | 0.166354052226522 | 2.89742362122238 | 0.0037624133916837 | 0.0497461706524524 |
| Fech | 397.486746162176 | 0.483410185097209 | 0.151124222344555 | 3.19876044751489 | 0.00138019801674881 | 0.02549468761835 |
| Hmgcl | 482.04931617622 | 0.483590246207688 | 0.177206592789491 | 2.72896306280298 | 0.00635338199057911 | 0.0687922178532032 |
| Slc30a9 | 943.929295480147 | 0.483838780759276 | 0.120490070573966 | 4.01559048355157 | 5.92971427987228e-05 | 0.00302469829321925 |
| Myom1 | 475.517542468527 | 0.484437767757321 | 0.13050062565514 | 3.71214900561083 | 0.000205506937852046 | 0.00737173273843467 |
| Flnb | 1392.92624507221 | 0.485330023320069 | 0.160388482976639 | 3.02596554511185 | 0.00247840542950337 | 0.0373785961736478 |
| Bag1 | 1481.46912276074 | 0.487391051662607 | 0.127078674316977 | 3.83534888353407 | 0.000125386181011292 | 0.00520259079419989 |
| Sec22b | 538.99843713573 | 0.489804976368115 | 0.159102025439596 | 3.07855902534736 | 0.00208004320016742 | 0.0338915392973821 |
| Fnip2 | 869.643222935777 | 0.492357546179903 | 0.182421274363388 | 2.69901385075906 | 0.00695452818345627 | 0.0727791694121504 |
| Ulk1 | 697.7575819984 | 0.495091303005573 | 0.18282444682633 | 2.70801477373469 | 0.00676870092391961 | 0.0717743839738763 |
| Klhl25 | 213.522247648717 | 0.49611162463991 | 0.173940734290692 | 2.8521877101588 | 0.00434194524286121 | 0.0540676720051698 |
| Senp2 | 1161.69307251957 | 0.498119703557394 | 0.112961672670411 | 4.40963462900166 | 1.0354517032897e-05 | 0.000772766640307479 |
| Ptk2b | 2150.02031643189 | 0.498332436926327 | 0.173561340651384 | 2.87121795127913 | 0.00408893493961525 | 0.051920407984837 |

| | | | | | | |
|---|---|---|---|---|---|---|
| Tbcel | 570.375650454622 | 0.498729950223233 | 0.168841844223902 | 2.9538290849386 | 0.00313857776967957 | 0.0439006098098576 |
| Galnt10 | 632.964050914281 | 0.501493322473298 | 0.161025170525872 | 3.11437845919078 | 0.00184332882981706 | 0.0312456662342024 |
| Sec23a | 392.736118756185 | 0.504493142020324 | 0.193678567619425 | 2.60479591635377 | 0.00919289989821432 | 0.0866313956509688 |
| Nek7 | 1736.3381389747 | 0.508075914649608 | 0.189782986450337 | 2.67714152966269 | 0.0074253261923126 | 0.0753523775786844 |
| Ubl3 | 4853.10408606561 | 0.509127457105074 | 0.129869593878892 | 3.92029759929683 | 8.84396869611059e-05 | 0.00398157619031375 |
| Hsp90b1 | 8181.79531749912 | 0.509166991766512 | 0.195388180333913 | 2.60592524530583 | 0.00916264566957488 | 0.086477215432091 |
| Scamp1 | 326.704292495306 | 0.510381782182595 | 0.198123554798876 | 2.57607825935036 | 0.00999280263421153 | 0.0913064628532722 |
| Lrrc8d | 1017.70001376381 | 0.510991488355447 | 0.147665582842202 | 3.46046437172501 | 0.000539244586468897 | 0.0138165894044565 |
| Sf3b5 | 186.614107522195 | 0.514071208423086 | 0.197447676564258 | 2.60358195836144 | 0.00922552070598637 | 0.0867131310189391 |
| Golt1b | 539.102899754994 | 0.514583037900008 | 0.200242267805899 | 2.56980228769088 | 0.0101756571138475 | 0.0922194841939565 |
| Cwc27 | 647.45627508076 | 0.515322697895817 | 0.192247091418533 | 2.68052272777396 | 0.00735072725970386 | 0.0749909056219329 |
| Itgav | 4157.96906992839 | 0.51699571490228 | 0.14648617092039 | 3.52931414381258 | 0.000416638238283604 | 0.0117122489076641 |
| Mettl6 | 295.007617100235 | 0.518622221587244 | 0.205487322271326 | 2.52386481002684 | 0.0116072556443121 | 0.0995084095357014 |
| Gdap2 | 404.435802428122 | 0.520540159584499 | 0.138694136533549 | 3.75315188222527 | 0.000174624977870522 | 0.00650754181686748 |
| 1110007C09F | 425.385529593464 | 0.520954593670184 | 0.145700895367647 | 3.57550715358105 | 0.00034954961701629 | 0.0106492924417018 |
| Etfb | 359.045762622696 | 0.521922262526121 | 0.152413646085167 | 3.42438013873428 | 0.000616203600566215 | 0.0150455914278742 |
| Cyth3 | 851.044111534238 | 0.522786050169945 | 0.144001767148437 | 3.63041412978674 | 0.000282966836156002 | 0.00930944147353473 |
| Timm17a | 234.091718559647 | 0.522851239067769 | 0.157094258469325 | 3.32826447103962 | 0.000873888633158609 | 0.019166945958035 |
| Psmd13 | 1382.84256324683 | 0.522949790704398 | 0.19431477420454 | 2.69125079575231 | 0.00711846588351805 | 0.0736347354648565 |
| Srp54a | 8093.59133940683 | 0.523262711696343 | 0.137242707328725 | 3.81268135758223 | 0.000137467290858381 | 0.00555867736125528 |
| Eprs | 2052.12211177391 | 0.523922786077487 | 0.149950740329657 | 3.49396598460052 | 0.0004759016501155 | 0.0127212171857797 |
| Arl1 | 777.089175818124 | 0.525281679516512 | 0.191741142432072 | 2.73953556786908 | 0.00615260572164036 | 0.0676300314890438 |
| Sar1b | 274.788552422635 | 0.52539416689207 | 0.189024592168737 | 2.77950165565265 | 0.00544423760075936 | 0.0627356705911338 |
| Mdh1 | 970.341585301416 | 0.525505245539565 | 0.13353002061655 | 3.93548389428192 | 8.30292325542022e-05 | 0.00379986334967686 |
| Tax1bp1 | 11935.805081993 | 0.527336252152492 | 0.162261213063977 | 3.24992179088771 | 0.0011543675052151 | 0.022486368764848 |
| Fam107b | 1989.84088071712 | 0.527503516249222 | 0.108710519018911 | 4.8523686668947 | 1.21995570621966e-06 | 0.000157743109920496 |
| Zdhhc20 | 640.05736581278 | 0.52792195630646 | 0.183215084984195 | 2.88143280533916 | 0.00395871633036211 | 0.051233712314029 |
| Afg3l2 | 737.963856374703 | 0.531018255169523 | 0.166616862377279 | 3.18706190713824 | 0.00143726008221242 | 0.026223394629043 |
| Srp54b | 8115.0473847633 | 0.531930330569095 | 0.140092303958289 | 3.7969989466907 | 0.00014645841146019 | 0.00577993136630165 |
| Lclat1 | 237.149830136261 | 0.534913924740802 | 0.183497268883732 | 2.91510564704772 | 0.00355568155047534 | 0.0480672707800013 |
| Dennd1b | 2071.50574670653 | 0.536443778405963 | 0.175968964994903 | 3.04851357409246 | 0.00229976522832066 | 0.0358170719032573 |
| Ift20 | 295.716986073591 | 0.537864804344905 | 0.173222480012047 | 3.10505197886267 | 0.00190245588933037 | 0.0317154979168519 |
| Atp6v0b | 1174.28370497018 | 0.539026495509349 | 0.145766048340047 | 3.69788782537271 | 0.000217400915871773 | 0.00769903880412138 |
| Atp6v1e1 | 1141.19257286309 | 0.539718851481203 | 0.166256559925048 | 3.24630108865792 | 0.00116915117492353 | 0.0226497579532223 |
| Slc23a2 | 510.23894329194 | 0.540838502827726 | 0.14815150606741 | 3.65057715026968 | 0.000261651699078343 | 0.00879023230740536 |
| Rab10 | 4048.40527792188 | 0.541770267288444 | 0.142699267898468 | 3.79658757376331 | 0.000146701553223047 | 0.00577993136630165 |
| Lif | 184.004732908019 | 0.543148745750258 | 0.215146196919723 | 2.52455657374656 | 0.0115844357343751 | 0.0995084095357014 |
| Dhx40 | 691.51499362403 | 0.543351038804555 | 0.179948437008121 | 3.01948184623596 | 0.00253207467190729 | 0.0376426074219372 |
| Gm6377 | 666.70318435436 | 0.543685601058011 | 0.157827575581373 | 3.44480740488658 | 0.000571466416148331 | 0.0143771641347725 |
| Dnaja2 | 2597.03990929316 | 0.544802494954962 | 0.179022601990213 | 3.04320509755939 | 0.00234072739952194 | 0.0361512342815056 |
| Furin | 3604.37576449751 | 0.545531866491113 | 0.164785970993624 | 3.31054799872632 | 0.000931134813003058 | 0.0199503258585626 |
| Pdia6 | 728.693293994826 | 0.548731326726428 | 0.211182961508225 | 2.59836931354453 | 0.00936676919789442 | 0.087675482727766 |
| Ept1 | 211.355904298227 | 0.549597424747624 | 0.190325090492663 | 2.88767720180756 | 0.00388097931498797 | 0.0506531572566505 |
| Trove2 | 249.188800958069 | 0.549680517066582 | 0.209665522999379 | 2.62170198134201 | 0.00874919010017274 | 0.0843643467169321 |
| Wdyhv1 | 113.235924904802 | 0.549714766082802 | 0.208089939847436 | 2.64171716559596 | 0.00824869070223709 | 0.0807442258880955 |
| Tnfaip2 | 7482.91943330902 | 0.551089001558103 | 0.143240905598557 | 3.84728789067118 | 0.000119432598765519 | 0.00501166225763233 |
| Hyou1 | 1270.45249954275 | 0.551730751663587 | 0.211223348688583 | 2.61207274237959 | 0.00899951032121362 | 0.0861226805265882 |
| Ubc | 298271.748054085 | 0.552237971483748 | 0.213268589877543 | 2.5894013356624 | 0.00961429697403072 | 0.0891503596768019 |
| Sub1 | 5624.35142996375 | 0.553068410474125 | 0.202999970214064 | 2.72447532820283 | 0.00644037420034878 | 0.0696663045796482 |
| Vmp1 | 616.181163763721 | 0.554477591798853 | 0.186494707829204 | 2.97315456429282 | 0.00294755979609296 | 0.0419678168150495 |
| Acaa1a | 378.859222231822 | 0.554487321821363 | 0.122592379505469 | 4.52301622709451 | 6.09645642500385e-06 | 0.000517500728595747 |
| Rars | 742.58502964618 | 0.557277803981512 | 0.168578347186452 | 3.30574960119374 | 0.00094722695759673 | 0.0201014575734268 |
| Cyb5a | 1461.84346824328 | 0.557647156971484 | 0.166645722777162 | 3.34630344948707 | 0.00081896717453707 | 0.0183065588616452 |
| St7 | 608.902463762304 | 0.560079121652173 | 0.206659637487055 | 2.71015244419587 | 0.00672522902186104 | 0.0714957425651002 |
| Ttc1 | 697.778006195321 | 0.561021886605008 | 0.178577301911739 | 3.14161923491425 | 0.00168016381147549 | 0.0293303321563697 |
| Psmb6 | 2032.19993012136 | 0.564349145490523 | 0.220684887279117 | 2.5572623139198 | 0.0105499628527495 | 0.0942293870864052 |
| Vapa | 2275.26118215329 | 0.564582493510028 | 0.118026447538314 | 4.78352526307083 | 1.72247231381445e-06 | 0.000212821023662408 |
| Dynlt1a | 291.443412525922 | 0.566098444901516 | 0.217124566559596 | 2.60725192856578 | 0.00912721795687336 | 0.0864220425055964 |
| Eif4e | 861.773247025738 | 0.566624799926642 | 0.167179222235423 | 3.38932549362334 | 0.000700647886803066 | 0.0165229739282886 |
| Psmd5 | 1219.32255725853 | 0.568172522701625 | 0.204203267645015 | 2.7823870266823 | 0.00539606482486613 | 0.0623744707406563 |
| Atp1a1 | 1691.28379602451 | 0.56924029004589 | 0.222301009006548 | 2.56067344268835 | 0.0104469502261674 | 0.0936440502971347 |
| Dusp4 | 401.451414030514 | 0.570023353546325 | 0.203187101318926 | 2.80541112032307 | 0.00502524383260188 | 0.059829455480228 |
| Atp6v1d | 1186.9262240876 | 0.574473833448126 | 0.188980170151168 | 3.03986303424638 | 0.00236685759090728 | 0.0363930620904491 |
| Sat1 | 3142.91034639789 | 0.576184831781132 | 0.139308354737322 | 4.13603931270009 | 3.5335168177557e-05 | 0.0020253972687342 |
| Gem | 1508.31427802431 | 0.578385074078308 | 0.228347428182914 | 2.53291696202071 | 0.0113117746195282 | 0.0982710420071511 |
| Tmem64 | 244.405040299858 | 0.578426861074312 | 0.186022208874924 | 3.10945055739677 | 0.00187435652062263 | 0.0314763302452612 |
| Dcaf6 | 657.307697155635 | 0.578865665923684 | 0.188148591593407 | 3.07664097308061 | 0.00209347284513082 | 0.0339007664027432 |
| App | 5484.88235474132 | 0.579058549672624 | 0.16911563714209 | 3.42403907443581 | 0.000616977490207792 | 0.0150455914278742 |
| Slc4a7 | 1294.45434589218 | 0.580256148938666 | 0.178091187472459 | 3.25819686630143 | 0.00112122593928559 | 0.0221064405050634 |

| | | | | | | |
|---|---|---|---|---|---|---|
| Prdx5 | 450.283466578094 | 0.580317511923803 | 0.177959428731204 | 3.26095400542297 | 0.00111038050101314 | 0.0219314940875064 |
| Dedd2 | 230.72821127051 | 0.580476400605355 | 0.222462393406099 | 2.60932372306951 | 0.00907213736525914 | 0.0862887252724732 |
| Bckdha | 205.22574475292 | 0.581456232448101 | 0.20050697143002 | 2.89993025330313 | 0.00373245702369604 | 0.049538755819916 |
| Tpt1 | 12005.1349383855 | 0.581550718260206 | 0.127146205474541 | 4.57387395942895 | 4.78787931346543e-06 | 0.000440010065832525 |
| Arrb1 | 1971.22632518959 | 0.585074485729197 | 0.15284561593464 | 3.82787875302479 | 0.000129252395965789 | 0.0053011215731587 |
| Zmym3 | 335.627815482824 | 0.586052668672618 | 0.201397488984065 | 2.90993036521418 | 0.00361509291364158 | 0.0485505231880367 |
| Atp1b3 | 1097.07785230229 | 0.586404021506195 | 0.189519661454118 | 3.09415929200655 | 0.00197371484931078 | 0.0324613320030173 |
| 9530082P21F | 156.013357446213 | 0.590374943619911 | 0.221261798299413 | 2.66821904258869 | 0.00762545257405686 | 0.0768070947676742 |
| Mtmr10 | 229.543059269412 | 0.590748231863917 | 0.193736215538096 | 3.04924007224532 | 0.00229421066489456 | 0.0358170719032573 |
| Nsa2 | 1972.32941174232 | 0.591602707601606 | 0.147043751747953 | 4.02331075322173 | 5.73856983294167e-05 | 0.00298191105337904 |
| Gpc1 | 3478.42504525839 | 0.591930526275148 | 0.102406663304449 | 5.78019542063755 | 7.46139045904503e-09 | 2.51426248195699e-06 |
| Pitpnc1 | 1358.20885634984 | 0.593612547885805 | 0.169466180962397 | 3.50283781999857 | 0.000460329648302184 | 0.0125111170194112 |
| Sdcbp | 1462.58598077065 | 0.595945297857571 | 0.229759060269714 | 2.59378366693349 | 0.00949261954842608 | 0.0887041423348723 |
| Acox1 | 1101.26092035473 | 0.596250333706018 | 0.16711367532059 | 3.56793262168505 | 0.000359808993928699 | 0.0107740972421658 |
| Lamc1 | 1417.75439742721 | 0.597000686202405 | 0.202237030333788 | 2.95198503071899 | 0.0031573823693269 | 0.043997608956034 |
| Pex5 | 553.76587562944 | 0.597394163066202 | 0.154660284665568 | 3.86262164432184 | 0.000112176675761124 | 0.0047977101325527 |
| Lims1 | 2189.71263253962 | 0.597598408407451 | 0.156417516469953 | 3.82053379886164 | 0.000133163154292186 | 0.00540428567784345 |
| Wbp5 | 201.164678235767 | 0.598139607830832 | 0.229059222403437 | 2.6112880396378 | 0.00902018844950724 | 0.0861722470434025 |
| Lmbrd1 | 718.852464339661 | 0.598985238589558 | 0.22040818890985 | 2.71761789592378 | 0.00657537250361256 | 0.0705069746220037 |
| Lrp1 | 11412.5398436772 | 0.59900416891532 | 0.227222189242185 | 2.63620454900586 | 0.00838391754237122 | 0.0817799676062877 |
| Laptm5 | 4663.26059832817 | 0.60007643695404 | 0.231420258293118 | 2.5930160193408 | 0.00951383390562663 | 0.0887496179978748 |
| Fam114a1 | 257.39556624302 | 0.600222878345873 | 0.167342820635005 | 3.58678595274208 | 0.00033477890848182 | 0.0104079249452115 |
| Ndufb3 | 133.548790717452 | 0.601345637365225 | 0.217233364473529 | 2.76820109481158 | 0.00563666639323618 | 0.0641593366645213 |
| Il1rn | 773.21963073614 | 0.602056821603496 | 0.179379273103248 | 3.35633438126913 | 0.000789830109353484 | 0.0179609628139279 |
| Slc25a22 | 134.732283236705 | 0.602532269256039 | 0.208961097277511 | 2.88346623896143 | 0.00393324806031875 | 0.0512007407368702 |
| Pdia3 | 2718.8054616132 | 0.602946061369066 | 0.227729413519066 | 2.64764244570711 | 0.0081055203751343 | 0.0798347090978684 |
| Als2 | 762.3074963106 | 0.604620509392409 | 0.238243427543557 | 2.53782660712379 | 0.0111543233526317 | 0.0972831966127569 |
| Atp7a | 878.894182814589 | 0.605418520999025 | 0.161223513480742 | 3.7551502750952 | 0.00017323750469028 | 0.00650754181686748 |
| Cdk2ap2 | 209.775820798385 | 0.605521466077081 | 0.182942791355028 | 3.30989519506114 | 0.000933309102041161 | 0.019958456182111 |
| Psmc1 | 2520.200482409 | 0.606366201168879 | 0.204274787887671 | 2.96838492620202 | 0.00299369172606936 | 0.0423817296553506 |
| Ern1 | 536.458096269271 | 0.608523032452516 | 0.179351972051314 | 3.39289847495187 | 0.00069157264867628 | 0.0163623145814473 |
| Hspa13 | 1143.58926219924 | 0.609441981740701 | 0.186523902724778 | 3.26736666367073 | 0.00108552987329997 | 0.0215699066553133 |
| Cipc | 171.966453023493 | 0.611470989011559 | 0.231179787945257 | 2.64500194608862 | 0.00816904472130648 | 0.0803148863131219 |
| Anxa11 | 2225.65409262315 | 0.611913048256761 | 0.139320025224493 | 4.39213994736762 | 1.12240409982032e-05 | 0.000816879143176314 |
| Ubap1 | 1296.2776468303 | 0.612082852651094 | 0.236176683401165 | 2.59163116289267 | 0.00955221212403118 | 0.0888132097150725 |
| Lpcat3 | 189.13876534598 | 0.612442102942747 | 0.189190980188215 | 3.23716332741372 | 0.00120724295713734 | 0.0231059237235236 |
| Slc36a4 | 187.897351145073 | 0.613308963257823 | 0.185921409484781 | 3.29875383882579 | 0.000971150320355044 | 0.0202992322600528 |
| Ranbp2 | 33658.2220696741 | 0.615005908262304 | 0.144556422142379 | 4.25443504444625 | 2.09577485058762e-05 | 0.00133171521934482 |
| Prdx3 | 327.462580309747 | 0.615614660350086 | 0.197109754602734 | 3.1232074819982 | 0.00178891589950148 | 0.0306513787403028 |
| 4933434E20F | 297.339975603387 | 0.61638274615934 | 0.174968931347282 | 3.52281254399348 | 0.000426993253695556 | 0.0118440553860813 |
| Thra | 93.6765971215437 | 0.617647497928488 | 0.238305969731951 | 2.59182553682236 | 0.00954681716765263 | 0.0888132097150725 |
| Btg3 | 305.181949911864 | 0.619494672601291 | 0.210961912922492 | 2.93652377350643 | 0.00331913378331569 | 0.0455101944148834 |
| Hspa4l | 1524.62931598698 | 0.620201109084174 | 0.128512690927244 | 4.82599115005144 | 1.39308694587042e-06 | 0.000178058929173323 |
| Anp32a | 4336.8272922858 | 0.62064957577749 | 0.165389861079007 | 3.75264584980216 | 0.00017497796791757 | 0.00650754181686748 |
| Zdhhc18 | 1457.67304266697 | 0.621873875243873 | 0.162492667482151 | 3.82708884579165 | 0.000129667721933378 | 0.0053011215731587 |
| Gsr | 2273.5269993752 | 0.622698335690509 | 0.190849705660118 | 3.26276812184068 | 0.0011032975192222 | 0.021830370842973 |
| Secisbp2l | 9082.9992129626 | 0.623960698824989 | 0.132556554731468 | 4.70712821473827 | 2.51230904973177e-06 | 0.000271231811971041 |
| Fopnl | 231.639702950541 | 0.62399078972842 | 0.182592152903592 | 3.41740200663435 | 0.000632218492716131 | 0.0153499337096144 |
| Nxt2 | 338.048123335378 | 0.625959712071521 | 0.201621756995809 | 3.10462383325293 | 0.00190521156550873 | 0.0317154979168519 |
| Odc1 | 1170.08077174141 | 0.626602832491415 | 0.221795115044053 | 2.82514262032759 | 0.00472595751829294 | 0.0575604026324398 |
| Slmo2 | 933.874458349729 | 0.630985533543024 | 0.139267600711453 | 4.53074175414536 | 5.87769515833386e-06 | 0.000506666643535405 |
| Mrpl51 | 273.442401960534 | 0.631618080170034 | 0.193222429556914 | 3.26886522241969 | 0.00107979722425311 | 0.0215561980434915 |
| Pkm | 14763.3732083646 | 0.632396383326379 | 0.105213724390813 | 6.01058832379475 | 1.84851265712845e-09 | 1.08186635511939e-06 |
| Vaultrc5 | 10068.007004361 | 0.633934831567045 | 0.19394427661759 | 3.26864418287015 | 0.00108064103390287 | 0.0215561980434915 |
| Atp13a3 | 11003.7848959502 | 0.634641476701056 | 0.182926223411978 | 3.46938489661892 | 0.000521651538077178 | 0.0135215969776648 |
| Rin2 | 2156.88899506148 | 0.634939280353756 | 0.212439599538425 | 2.98879908328443 | 0.00280076237231303 | 0.040605577027537 |
| Emc1 | 897.750121385283 | 0.635654598749211 | 0.200931752708829 | 3.16353483299546 | 0.00155865698496513 | 0.0277316250764995 |
| Hcfc2 | 83.0899506504523 | 0.636243524043457 | 0.231472138035516 | 2.74868297084566 | 0.00598352286624552 | 0.0666701145016534 |
| Serp1 | 2161.51477535566 | 0.63656119890489 | 0.192953829111428 | 3.29903377318978 | 0.000970182381471635 | 0.0202992322600528 |
| Arrdc3 | 1152.40688791324 | 0.638499482508927 | 0.24157697594238 | 2.64304774914154 | 0.00821634473204335 | 0.0805694474606014 |
| Bola3 | 255.009628091383 | 0.642991439400035 | 0.250067013443089 | 2.57127651722993 | 0.0101324388348259 | 0.0919372431451346 |
| Mrfap1 | 5973.42177581953 | 0.648199923582745 | 0.159027253927923 | 4.07603041348204 | 4.58110297278928e-05 | 0.00250775915881342 |
| Psmd2 | 5968.67645259417 | 0.654322285699741 | 0.256163004715812 | 2.55431999802489 | 0.0106395425495412 | 0.0947353295731959 |
| Rcbtb2 | 405.083057540597 | 0.66052573564267 | 0.134050166839642 | 4.92745179819751 | 8.33089666034156e-07 | 0.000117265279573415 |
| Alox5 | 245.055157058767 | 0.662971719803835 | 0.239851292509754 | 2.76409483920907 | 0.00570809460252287 | 0.064835558713028 |
| Serpinb6a | 1850.70726594525 | 0.664522378607187 | 0.21639053536168 | 3.07094012913499 | 0.00213385933883782 | 0.034227506100699 |
| Calr | 4577.81958270621 | 0.665039935073719 | 0.190586036468999 | 3.48944732465695 | 0.000484020441057245 | 0.012771936267561 |
| Tmem263 | 138.266085176032 | 0.669025746784261 | 0.252439395946738 | 2.65024301882508 | 0.00804338915059726 | 0.0794338253593619 |
| Ncf4 | 1101.23120228891 | 0.671660560455381 | 0.170548255194905 | 3.93824351757689 | 8.208026930736e-05 | 0.00378727217716947 |

| | | | | | | |
|---|---|---|---|---|---|---|
| Plcd3 | 75.2323416329468 | 0.673098408144592 | 0.264097094902126 | 2.54867782015566 | 0.0108132144746492 | 0.0957348287882957 |
| Cstb | 910.245897746997 | 0.673848512509518 | 0.264488220899031 | 2.54774488715987 | 0.0108421726244537 | 0.0958386006231515 |
| Agpat5 | 99.2614588344296 | 0.675401259585642 | 0.225128344935444 | 3.00007206902052 | 0.00269915733436383 | 0.0395449664797442 |
| Gnl3l | 585.34884421571 | 0.678727102446814 | 0.169077319194086 | 4.01430011832452 | 5.9622444371905e-05 | 0.0030274044813497 |
| Ift80 | 110.882343468475 | 0.680545167270036 | 0.236263199581226 | 2.88045353011513 | 0.00397103486367744 | 0.0512867685064961 |
| Plek | 3065.35315612647 | 0.68329867449229 | 0.139168499943222 | 4.90986591628898 | 9.11386877470736e-07 | 0.000126682775968432 |
| Rnf11 | 1304.06949279744 | 0.683712191005204 | 0.195621325451408 | 3.49508004522256 | 0.000473919602392863 | 0.0127045454962423 |
| Ppa1 | 372.523234558499 | 0.683890566659915 | 0.22416994736436 | 3.05076828852682 | 0.00228256652301854 | 0.0358170719032573 |
| Bace1 | 330.687950417168 | 0.685383114691132 | 0.209701785389399 | 3.26837043098337 | 0.00108168691621117 | 0.0215561980434915 |
| Fbxo32 | 378.779997391685 | 0.685490646168199 | 0.241041508912505 | 2.84386971049465 | 0.00445692688253764 | 0.0551290622178182 |
| Nfe2l2 | 1291.91341396027 | 0.695398095778234 | 0.185129635247156 | 3.75627648620303 | 0.000172460157258162 | 0.00650087101257884 |
| Pgam1 | 1321.70385390371 | 0.695910537988616 | 0.244894051533234 | 2.84168003931356 | 0.00448765051769668 | 0.05538587542373 |
| Ccpg1 | 1231.03200716239 | 0.698203807963511 | 0.231366706672061 | 3.01773672628338 | 0.00254670047403827 | 0.0378094916839861 |
| Ikbkg | 1363.02689537012 | 0.698287925633453 | 0.182313995494265 | 3.83013889712827 | 0.000128070947597378 | 0.00528952718059445 |
| Pcx | 210.990603631381 | 0.699849012456048 | 0.259151406538992 | 2.70054105359737 | 0.00692267927074078 | 0.0725543765227498 |
| Creld2 | 198.5291131096 | 0.700588107373448 | 0.252899124351591 | 2.7702274935498 | 0.00560171524363816 | 0.0639313583397809 |
| Htatip2 | 432.137348497195 | 0.701685891280527 | 0.247302062016869 | 2.83736369020919 | 0.00454877650678889 | 0.0559389492098291 |
| Sqle | 2065.04450425902 | 0.7043386407547 | 0.160659529496752 | 4.38404521014698 | 1.16495580109651e-05 | 0.00083576183923827 |
| Gpr65 | 453.852883552664 | 0.708184423974093 | 0.227580993937312 | 3.111790715569 | 0.00185956278154425 | 0.0314260457914469 |
| Gngt2 | 543.63546142811 | 0.714772357152204 | 0.226224129665558 | 3.15957611687533 | 0.00157998823318441 | 0.0280662446533717 |
| Adgre1 | 751.373665721986 | 0.715974344192312 | 0.264748848040422 | 2.70435301037834 | 0.00684375408321628 | 0.0722552003850785 |
| Gbe1 | 327.816823923776 | 0.716645323064559 | 0.170868438752259 | 4.19413513869357 | 2.73914654310969e-05 | 0.00168283478228618 |
| Ugcg | 769.328448553705 | 0.721454608699489 | 0.224777909343667 | 3.2096330587204 | 0.00132904517502522 | 0.0248386257920679 |
| Mob4 | 617.487372127711 | 0.722030425486918 | 0.125232553199467 | 5.76551708833158 | 8.14078526256709e-09 | 2.66251565058077e-06 |
| Adgrl2 | 1245.91161689886 | 0.722224783061643 | 0.253581186570505 | 2.848100810747 | 0.00439809904111282 | 0.0546445378068989 |
| Sqstm1 | 23024.946329146 | 0.725756417441071 | 0.22080240840491 | 3.28690444404107 | 0.00101295199593822 | 0.0208980077826215 |
| Rdh11 | 151.317047019185 | 0.727219295884033 | 0.244451754815779 | 2.97489906109315 | 0.00293084969608889 | 0.041837032888971 |
| Edem3 | 988.384108176041 | 0.731037740416711 | 0.189649223765481 | 3.85468353575077 | 0.000115879424770576 | 0.00491824123453743 |
| Abca1 | 5191.89317779285 | 0.731560676516952 | 0.174177041013011 | 4.20009820044138 | 2.66799239482736e-05 | 0.00165743438159107 |
| Fos | 873.905953197261 | 0.732117103199913 | 0.225117444159312 | 3.25215625085813 | 0.00114533040695148 | 0.0223832585681906 |
| Eepd1 | 188.678881963924 | 0.733638715467716 | 0.240028116583112 | 3.05646990823963 | 0.00223959959526645 | 0.0353754936070496 |
| Mpp7 | 254.68158035713 | 0.734433459588358 | 0.206055051399491 | 3.56425845714633 | 0.000364886283596249 | 0.0107913177489103 |
| Sash1 | 198.545207022057 | 0.73865959362065 | 0.271710154569492 | 2.7185571874964 | 0.00655673203619549 | 0.0704927963003417 |
| Sesn1 | 847.812846204525 | 0.738984315861104 | 0.124967215959866 | 5.91342545470831 | 3.35065207823181e-09 | 1.48609840579005e-06 |
| Crtc1 | 172.102780247105 | 0.73913354237446 | 0.238910094261504 | 3.09377276275914 | 0.00197628792860096 | 0.0324613320030173 |
| Taf13 | 379.037099789568 | 0.739699967244311 | 0.238406008361334 | 3.10269012231942 | 0.00191770324614575 | 0.0318281493987175 |
| Prdx6 | 743.90895524115 | 0.741177164163858 | 0.204489869847882 | 3.6245177558928 | 0.000289501131569989 | 0.00935829239261127 |
| Plekha8 | 597.147252193292 | 0.741918065614022 | 0.171424743922257 | 4.32795201345316 | 1.50502248401182e-05 | 0.00101925977186225 |
| Top1 | 6801.21466600745 | 0.743450873708592 | 0.155845069426044 | 4.77044847454346 | 1.83816202274055e-06 | 0.000219890846249786 |
| Flrt3 | 3760.60452046789 | 0.743749013864618 | 0.129621295559276 | 5.73786128780437 | 9.58795911978242e-09 | 2.88157041654001e-06 |
| Nrd1 | 5415.11066808286 | 0.743938038724593 | 0.187562364916465 | 3.96635028064357 | 7.29815837940791e-05 | 0.00350338257157202 |
| Sfn | 302.965163298488 | 0.745507438649555 | 0.162671460403384 | 4.58290247595297 | 4.58565940981884e-06 | 0.00042999819888111 |
| Rora | 3744.64244000175 | 0.750184605418011 | 0.133854055479749 | 5.60449664919944 | 2.08861031055735e-08 | 5.80633666334943e-06 |
| Cadm1 | 175.290188496262 | 0.750256330858867 | 0.249381806135596 | 3.00846458081602 | 0.00262571406588631 | 0.0388255841082969 |
| Irgq | 415.193907414097 | 0.754387087523779 | 0.237646498815206 | 3.17440859126812 | 0.00150142173272661 | 0.0270596591052187 |
| Lgals8 | 339.028272033779 | 0.757882350079807 | 0.293138512779512 | 2.58540695623252 | 0.00972641205308339 | 0.0895461552244638 |
| Igf2r | 542.926112341172 | 0.759729482609951 | 0.284224274733543 | 2.67299295009966 | 0.00751778303532415 | 0.0758687928152643 |
| Susd3 | 113.22089688932 | 0.760967844726594 | 0.265238180794076 | 2.86899813008968 | 0.00411774233035329 | 0.0520332754738428 |
| Tuba4a | 558.282137394695 | 0.761921789076213 | 0.297010896357224 | 2.56529911333565 | 0.0103086895410443 | 0.0928199414545855 |
| Wdfy1 | 534.949793303228 | 0.763185667612754 | 0.200334811995571 | 3.80955092133276 | 0.000139219432274324 | 0.00558888118011007 |
| Smim3 | 2032.37876283758 | 0.764044463825746 | 0.128479218731272 | 5.94683304717029 | 2.73379987661041e-09 | 1.38181157399581e-06 |
| Tex264 | 239.054915061018 | 0.766749204905621 | 0.222440809635921 | 3.44698082227176 | 0.000566888821353093 | 0.0143594617162788 |
| Emilin2 | 134.055265510301 | 0.770054042614956 | 0.2951453399743 | 2.609067257108 | 0.00907893961949583 | 0.0862887252724732 |
| Alas1 | 948.92733161424 | 0.771365223115397 | 0.201740973951212 | 3.82354267458797 | 0.00013154781113682 | 0.00535828446828364 |
| Zfp367 | 114.512031638452 | 0.773453525053562 | 0.266975894852437 | 2.89709123545054 | 0.00376640204889643 | 0.0497461706524524 |
| Lrrc20 | 179.306443379534 | 0.775077502299173 | 0.19492141727185 | 3.9763588483363 | 6.99785034707416e-05 | 0.00340232658887635 |
| Alg2 | 123.020831277383 | 0.779300866068465 | 0.223345390223365 | 3.48921849378264 | 0.000484435004103769 | 0.012771936267561 |
| Serinc2 | 741.330632368348 | 0.779674501840281 | 0.187397472589557 | 4.16053904605184 | 3.17497327510578e-05 | 0.0018581948852198 |
| Insig2 | 267.125567299682 | 0.780416529208912 | 0.257234722085847 | 3.03386931157942 | 0.00241438995639112 | 0.0366776179167612 |
| Ddit4 | 1339.00028717823 | 0.784858671538543 | 0.177965201626192 | 4.41018055421365 | 1.03284465522609e-05 | 0.000772766640307479 |
| Ak6 | 124.574542100206 | 0.789009458583234 | 0.253116128041136 | 3.11718366067533 | 0.00182587784049475 | 0.0310665111770838 |
| Glce | 201.959744641099 | 0.792659596916296 | 0.307871134595779 | 2.57464733729267 | 0.0100342339356046 | 0.0913098865498548 |
| Arl4d | 151.474143525892 | 0.793126500752293 | 0.257668911643845 | 3.07808379246219 | 0.00208336325848169 | 0.0338915392973821 |
| Ankrd28 | 550.269733194849 | 0.798837366544933 | 0.308373790651256 | 2.59048398652124 | 0.00958410804575047 | 0.0889610028954468 |
| Pgap1 | 230.730528139052 | 0.800116102140086 | 0.24436707389006 | 3.27423858461413 | 0.00105947117068124 | 0.0213817049328046 |
| Dnajb4 | 878.715975856253 | 0.80052963360051 | 0.278874282505863 | 2.87057532307118 | 0.00409725568103334 | 0.051920407984837 |
| Dnah2 | 1605.15735383824 | 0.804262017730009 | 0.272412014483549 | 2.95237351867452 | 0.00315341227589971 | 0.0439974209636195 |
| Ust | 98.0741031914432 | 0.807080864401422 | 0.248773790182078 | 3.24423591331997 | 0.00117766164288495 | 0.0227510103559205 |
| Runx1 | 501.139728594003 | 0.808533698944417 | 0.286009148322715 | 2.82695047933263 | 0.00469935944975317 | 0.0572992073259378 |

| | | | | | | |
|---|---|---|---|---|---|---|
| Trim23 | 173.091581119377 | 0.809226174228857 | 0.217613783816533 | 3.71863473000913 | 0.000200302448878107 | 0.00726223825838619 |
| Arhgdib | 2115.45042024747 | 0.810387824245149 | 0.177976628943825 | 4.55333843018756 | 5.28012271726733e-06 | 0.000469719716928102 |
| Tsc22d1 | 683.696354847089 | 0.81593002352428 | 0.156752045198062 | 5.20522729061253 | 1.93759301628796e-07 | 3.47957218576127e-05 |
| Mesdc1 | 1124.86956910194 | 0.816754246987169 | 0.162481306058692 | 5.02675825791391 | 4.98840537945711e-07 | 7.29882471310041e-05 |
| Sec24d | 2338.22396675343 | 0.818070680533922 | 0.184677126561369 | 4.4297347254971 | 9.43490721854556e-06 | 0.000723559781173976 |
| Thap11 | 205.246033749286 | 0.818386671173169 | 0.217005561761151 | 3.77127048971183 | 0.000162418499910358 | 0.00624272372995241 |
| Golim4 | 572.738813195565 | 0.827601697693996 | 0.187544353840448 | 4.41283184882267 | 1.02027237901705e-05 | 0.000772766640307479 |
| Pim3 | 286.540970635067 | 0.830772675484893 | 0.24215793806678 | 3.43070593562698 | 0.000602012823823078 | 0.0148852235505246 |
| Rilpl2 | 579.333517564662 | 0.831351320188543 | 0.257360966274645 | 3.23029297030755 | 0.00123663428736772 | 0.0233866892440971 |
| Eaf1 | 1411.35655880944 | 0.832981655342523 | 0.208848837817289 | 3.98844285679606 | 6.65084187038549e-05 | 0.00329852479589297 |
| Serpinb9b | 3473.41017766209 | 0.835038403878287 | 0.207741579879502 | 4.01960168187149 | 5.82966176962827e-05 | 0.00300634087497222 |
| Cks2 | 99.6302232642475 | 0.84366469636803 | 0.327595332431589 | 2.5753257535933 | 0.0100145718535347 | 0.0913098865498548 |
| Slc15a3 | 2441.27751934246 | 0.844547731719436 | 0.159100032498238 | 5.30828132752762 | 1.10663745417251e-07 | 2.12169111903418e-05 |
| Tmem189 | 684.658629149216 | 0.847699242352955 | 0.280736606854815 | 3.01955363730441 | 0.00253147464151658 | 0.0376426074219372 |
| Trp53inp1 | 956.180020817245 | 0.849535418331304 | 0.17836964121509 | 4.76278032822231 | 1.90943668539919e-06 | 0.000221176416058739 |
| Acaa1b | 132.375343798966 | 0.853345862954272 | 0.215116182941586 | 3.96690686532866 | 7.28114283609077e-05 | 0.00350338257157202 |
| Sod2 | 1674.58570611353 | 0.854127499963628 | 0.277019076370182 | 3.08328044102729 | 0.0020473212111283 | 0.0334907875446134 |
| Hs2st1 | 374.898779360437 | 0.856535143209595 | 0.161904448590736 | 5.29037435762344 | 1.22066252470377e-07 | 2.26229454578433e-05 |
| Abr | 1523.82090604971 | 0.856693142937492 | 0.298494723164969 | 2.87004451487079 | 0.00410414016354961 | 0.051920407984837 |
| Tmem101 | 66.0899322146335 | 0.857201794089531 | 0.266628923606833 | 3.21496176218881 | 0.00130461899321633 | 0.0244567078075687 |
| Eif4g3 | 2014.5480823561 | 0.869047102014154 | 0.260805239314469 | 3.33216887934636 | 0.000861719471319916 | 0.0189989381778621 |
| Btd | 108.916745749541 | 0.876912375938351 | 0.275998115857353 | 3.17724044316149 | 0.00148683711843895 | 0.0269277341319889 |
| Sgtb | 119.615458662037 | 0.87708036168103 | 0.287550641896962 | 3.05017702584477 | 0.0022870651734215 | 0.0358170719032573 |
| Ccrl2 | 262.226046759294 | 0.878447489108795 | 0.202975133426555 | 4.3278576753672 | 1.50566708979216e-05 | 0.00101925977186225 |
| Igfbp4 | 320.478361800033 | 0.881111998269986 | 0.20143410752201 | 4.37419466399806 | 1.21881583887487e-05 | 0.000857799501790416 |
| Sesn3 | 173.359193390868 | 0.88835704363342 | 0.344972137887174 | 2.57515592150217 | 0.0100194907598795 | 0.0913098865498548 |
| Ier3 | 3733.32570684323 | 0.892201995606545 | 0.182942606322824 | 4.87695028260475 | 1.0773857647647e-06 | 0.000144343731375704 |
| Wnt9a | 321.019392130096 | 0.895042079212298 | 0.236923808152391 | 3.77776335013406 | 0.00015824313656883 | 0.00610994332862981 |
| Hspa5 | 11365.1489535136 | 0.896299781429207 | 0.255251066197368 | 3.51144382972395 | 0.000445679635938894 | 0.0122067919991145 |
| Tmem26 | 334.714201965728 | 0.896844318379837 | 0.181665095106967 | 4.93680042306289 | 7.94146651215787e-07 | 0.000113216804634866 |
| Arl6ip5 | 1390.25586492178 | 0.896902161444509 | 0.226827851678996 | 3.95410949231136 | 7.68202267218151e-05 | 0.00361966492011265 |
| Itgb3 | 208.272819812524 | 0.903308402336981 | 0.328525158815258 | 2.74958668491184 | 0.00596704775666861 | 0.0666435371140802 |
| Denr | 1609.69120400662 | 0.908389794950004 | 0.162323242045011 | 5.59617823982418 | 2.19128424098471e-08 | 5.9431904292073e-06 |
| Fam63a | 248.573091779615 | 0.908799261694065 | 0.239651043540744 | 3.79217736032749 | 0.000149332197312243 | 0.00584709166940897 |
| Fam129b | 4310.42612929782 | 0.914854172682238 | 0.229178401517482 | 3.9918865243174 | 6.5549739514443e-05 | 0.00326866862511483 |
| Tnfaip8l2 | 92.9422215806647 | 0.916013626726388 | 0.314145717118775 | 2.91588768144833 | 0.00354678158703429 | 0.0480392341629979 |
| Arhgap6 | 131.47944563608 | 0.916317384840396 | 0.308310427979471 | 2.97206095442679 | 0.00295807949829731 | 0.0420637391573735 |
| Errfi1 | 3220.67759448611 | 0.917426970733259 | 0.271427028221357 | 3.3800133197681 | 0.000724823170342259 | 0.0168973451870145 |
| Galc | 245.840220425894 | 0.920173526063376 | 0.319634438618849 | 2.8788309859209 | 0.00399152186015588 | 0.0514318923348011 |
| Sema4b | 277.687479341518 | 0.922344337405684 | 0.255775125426965 | 3.60607520323181 | 0.00031086308153055 | 0.00993332605350493 |
| Ninj1 | 844.070080653652 | 0.926268466885312 | 0.138001002956052 | 6.71204155798995 | 1.9192001291268e-11 | 1.64165426429923e-08 |
| Tfrc | 161.36882196451 | 0.927325198566473 | 0.274900051390578 | 3.37331766173055 | 0.00074268217244985 | 0.0171697001198385 |
| Gprc5c | 462.910120960416 | 0.940516712158481 | 0.335919892561926 | 2.7998244015427 | 0.00511304122652744 | 0.0604046129010099 |
| Cd84 | 125.318623525867 | 0.940940151817959 | 0.358403558005568 | 2.62536498536474 | 0.00865561289417643 | 0.0836960133767321 |
| Ssh2 | 435.263399799895 | 0.941692101598043 | 0.217194589889988 | 4.33570699010054 | 1.45292378402757e-05 | 0.00099731558508559 |
| Gm13889 | 219.070858096442 | 0.951614529353631 | 0.21882729430435 | 4.34870125492712 | 1.36946111149079e-05 | 0.000951775472486097 |
| Ecm1 | 51.4929525696018 | 0.955559545843262 | 0.332160156990116 | 2.87680363142319 | 0.00401725502658439 | 0.0516001615010989 |
| Rab20 | 41.9054777125525 | 0.957161868617276 | 0.374701078265407 | 2.55446787889759 | 0.0106350241787182 | 0.0947353295731959 |
| Abcc5 | 407.097126567267 | 0.959024506413067 | 0.290455019106643 | 3.30180042804133 | 0.000960663955290398 | 0.0201939190601687 |
| Slc2a1 | 1965.35050980568 | 0.959274122626791 | 0.239701128571576 | 4.00195914113248 | 6.28201487297447e-05 | 0.00314666690934577 |
| Klf10 | 212.344510497188 | 0.962981190506405 | 0.292448138055236 | 3.2928272236923 | 0.000991854052755948 | 0.0206536530225228 |
| Atp6v1a | 3601.1014604509 | 0.967724624392012 | 0.281479994743635 | 3.43798721921035 | 0.000586055359085552 | 0.0146119632130747 |
| Slc27a1 | 49.4954256396954 | 0.96865910129353 | 0.377323194741174 | 2.56718673750757 | 0.0102527382020818 | 0.0924658952207219 |
| Satb1 | 1997.56477549336 | 0.969609001071259 | 0.258562561725282 | 3.74999765859936 | 0.000176836221544369 | 0.00655472927857793 |
| Cdk14 | 208.702861303063 | 0.971452711271877 | 0.318608644511351 | 3.04904693581617 | 0.00229568612139265 | 0.0358170719032573 |
| Fgd3 | 1794.15689207481 | 0.975230556174665 | 0.268808969519887 | 3.62796880593865 | 0.000285659764407575 | 0.00932154447170718 |
| Slc7a8 | 1789.42209751288 | 0.988567954547103 | 0.169711451189252 | 5.82499264262793 | 5.71151158719922e-09 | 2.19006927067777e-06 |
| Cat | 1859.71835367592 | 0.991227360248099 | 0.280628587922005 | 3.53216815003749 | 0.000412167180618769 | 0.0117122489076641 |
| 1700071M16 | 319.659756015609 | 0.994201873557699 | 0.278309448255623 | 3.57228933400974 | 0.000353874126221511 | 0.0107222896010442 |
| Utp14b | 620.386620925312 | 0.995684833118614 | 0.30311740298386 | 3.28481579519085 | 0.00102049065797848 | 0.0208984458871468 |
| Ipo13 | 393.702738817512 | 0.996527876390939 | 0.212341703897492 | 4.69303889956545 | 2.69176282267612e-06 | 0.000285070500839604 |
| P3h1 | 570.316018070885 | 0.997155367718107 | 0.21216158586896 | 4.69998074172575 | 2.6018602626188e-06 | 0.000278198905003087 |
| Adgrg1 | 126.961783521535 | 1.0002231009189 | 0.315234090919494 | 3.17295346452341 | 0.001508967087408 | 0.027140314255265 |
| Fermt3 | 815.167497523159 | 1.00408150290331 | 0.263432729504524 | 3.81152905636224 | 0.00013810981725699 | 0.00556442452136858 |
| Slc22a4 | 385.663135676096 | 1.00807993208545 | 0.270508698955271 | 3.72660818664517 | 0.000194073791314111 | 0.00711348969992236 |
| Taldo1 | 4833.54286952939 | 1.009814830302 | 0.275450575994271 | 3.66604726331376 | 0.000246328418531042 | 0.00840236814130427 |
| Acsl3 | 480.400019648201 | 1.01272696548078 | 0.338122545172163 | 2.99514770588612 | 0.00274311972146471 | 0.0400308284812173 |
| Krt16 | 81.0063815132656 | 1.01400432955251 | 0.309169376071684 | 3.27976962801582 | 0.00103891882947633 | 0.0211589329373202 |
| Pag1 | 152.036766706908 | 1.01413798125391 | 0.259601649026066 | 3.90651594494335 | 9.36364671503704e-05 | 0.00414600572618915 |

| | | | | | | |
|---|---|---|---|---|---|---|
| Hand1 | 1783.92548344106 | 1.01608484141566 | 0.26953149440192 | 3.76981860197937 | 0.000163366241494258 | 0.00624272372995241 |
| Hfe | 101.128850448979 | 1.01641213152329 | 0.317601704630912 | 3.20027291007292 | 0.00137297516052598 | 0.0254458063084149 |
| Gabarapl1 | 2163.70192714213 | 1.02190399357912 | 0.278608265726154 | 3.66788828362886 | 0.000244561990766267 | 0.00839360906580522 |
| Zeb2 | 3419.26274777438 | 1.02345179699933 | 0.265975546752771 | 3.84791688369251 | 0.000119126452723147 | 0.00501166225763233 |
| Mitf | 588.736332028801 | 1.02995781318814 | 0.283107965501232 | 3.63803897698405 | 0.000274721856735158 | 0.00906500607387226 |
| Srxn1 | 2224.93460451148 | 1.0339486679423 | 0.228704399800182 | 4.52089539530351 | 6.15786128818442e-06 | 0.000518753163065233 |
| Coq7 | 125.824312204903 | 1.03454629959499 | 0.308878098074611 | 3.34936761798207 | 0.000809962509490264 | 0.0182178645074313 |
| Gas2l3 | 378.200332797524 | 1.03804411837861 | 0.308661625766119 | 3.36304882669528 | 0.000770867281131032 | 0.0176379509592121 |
| Lpar6 | 288.564987041079 | 1.05398276957342 | 0.270303290166622 | 3.8992598607428 | 9.64871782592953e-05 | 0.00424085937645598 |
| Mid1 | 201.893697009456 | 1.05807016108855 | 0.380264496770035 | 2.78245844688576 | 0.00539487732363154 | 0.0623744707406563 |
| Entpd1 | 429.497867103501 | 1.06000929072382 | 0.259960163005691 | 4.07758357460569 | 4.55061749787822e-05 | 0.00250509240477257 |
| Tm4sf19 | 109.752001184221 | 1.06186271759443 | 0.224283486152076 | 4.73446679384338 | 2.19631724148794e-06 | 0.000246697451771171 |
| Whrn | 1105.15094642544 | 1.06215498526774 | 0.209367769472503 | 5.07315422972609 | 3.9127511754356e-07 | 5.80130574277919e-05 |
| Acpp | 1271.88985037839 | 1.06881169832305 | 0.179012936078725 | 5.97058358873583 | 2.36406423208882e-09 | 1.25182829813465e-06 |
| Arhgef10l | 272.115878115691 | 1.07204897881873 | 0.270909618860456 | 3.95722006227817 | 7.58270693092568e-05 | 0.00358807238603802 |
| Def6 | 324.774982127599 | 1.07921012511676 | 0.25991707762264 | 4.15213242233976 | 3.29391558689006e-05 | 0.00191771420556112 |
| Il1a | 211.689000447089 | 1.08682610964607 | 0.348706934181008 | 3.11673214127115 | 0.00182867642510275 | 0.0310665111770838 |
| Atp8a1 | 1302.56248225559 | 1.08881731724309 | 0.237620086672148 | 4.58217708987443 | 4.60159943047232e-06 | 0.00042999819888111 |
| Pgd | 3014.75575790739 | 1.08933225119818 | 0.286343003372156 | 3.80429149086767 | 0.000142210641374555 | 0.00568842565498219 |
| Gyk | 227.887299934413 | 1.08964652140604 | 0.345144639902072 | 3.15707212406717 | 0.00159361924158898 | 0.028218226061257 |
| Fuca2 | 197.883592301498 | 1.09134238653965 | 0.411037248148063 | 2.65509364773318 | 0.00792863941215959 | 0.0786498396638846 |
| Gpnmb | 6538.93524367041 | 1.09231432096266 | 0.253414540811818 | 4.31038533725573 | 1.62970314953036e-05 | 0.00108516760615434 |
| Abi3 | 68.091170107812 | 1.09476939923576 | 0.380819856638611 | 2.87476973732142 | 0.00404322245202053 | 0.0518577089578642 |
| Lilr4b | 220.565105231065 | 1.10482232235479 | 0.227248401112522 | 4.86173859506163 | 1.16359217794479e-06 | 0.00015222523551466 |
| Tgm2 | 721.726562488058 | 1.11572908047004 | 0.284851377877073 | 3.91688145862341 | 8.97018291335208e-05 | 0.00402211427405141 |
| Slc6a8 | 266.044112169507 | 1.11687303755016 | 0.277974480383887 | 4.01789774373441 | 5.87196669174078e-05 | 0.0030090446825879 |
| Sqrdl | 145.889412331616 | 1.12141748796986 | 0.425765249439078 | 2.63388684127524 | 0.00844136156757657 | 0.0821960951238629 |
| Slc16a3 | 680.091367263635 | 1.12589891299373 | 0.316572366400284 | 3.55652935155467 | 0.000375786512211129 | 0.011084206938429 |
| 2610301B20F | 25.3234405019187 | 1.12993315785677 | 0.447782611187185 | 2.52339668764947 | 0.011622720690148 | 0.0995084095357014 |
| Tma16 | 823.287326443903 | 1.13163306614608 | 0.327738941641633 | 3.45284896716199 | 0.000554699482204256 | 0.0141473812892462 |
| Ahr | 1517.1321262823 | 1.13744076870559 | 0.255029734651004 | 4.4600319655358 | 8.19474297294546e-06 | 0.000655579437835637 |
| Blvrb | 1420.204267066 | 1.14901596377561 | 0.455353192182893 | 2.52335106792027 | 0.0116242287757982 | 0.0995084095357014 |
| Cxcl1 | 3917.53765632906 | 1.15025356951841 | 0.166537811066366 | 6.90686134369829 | 4.95493795953859e-12 | 5.0089918281881e-09 |
| Hephl1 | 489.788464011219 | 1.15099540485016 | 0.31740330647463 | 3.62628675055141 | 0.000287526056996004 | 0.00932154447170718 |
| Gss | 198.54249109314 | 1.15371698730864 | 0.285285078255757 | 4.04408458501406 | 5.25279542067106e-05 | 0.00275523986216331 |
| Frk | 122.584242451077 | 1.15378107651292 | 0.406676715616803 | 2.83709647542272 | 0.00455258534486469 | 0.0559389492098291 |
| Clec4a1 | 137.092564310373 | 1.15861128657171 | 0.425357193474828 | 2.72385492556687 | 0.00645248429787826 | 0.0697199266323424 |
| Cxx1c | 36.079205522511 | 1.16024814625307 | 0.382136814982302 | 3.0362113797039 | 0.00239571347213355 | 0.0365372098580231 |
| Sh3bgrl2 | 707.001184897986 | 1.16634373101139 | 0.245055476230101 | 4.7595089444817 | 1.9406452574136e-06 | 0.000222473971777724 |
| Tmem51 | 155.696490937357 | 1.17935782343678 | 0.346329577291823 | 3.40530494871085 | 0.000660902071379699 | 0.0158198992577674 |
| Fli1 | 166.704319492472 | 1.18352057968891 | 0.297050237735461 | 3.98424383939697 | 6.76953352028318e-05 | 0.00333085012148447 |
| Chsy3 | 50.9422653729426 | 1.20255577815976 | 0.452131669532304 | 2.65974683747262 | 0.00781994056675663 | 0.0780422841010338 |
| Epas1 | 212.046591401738 | 1.20268478643967 | 0.387481542403752 | 3.10385051886285 | 0.0019101981455885 | 0.0317509766501406 |
| Acss2 | 36.7808682886633 | 1.20924102080395 | 0.434969753320818 | 2.78005772027109 | 0.00543492372146446 | 0.0626949897684443 |
| Lmo4 | 38.7088504418209 | 1.21564438510808 | 0.441359121357179 | 2.75432029448032 | 0.00588141789339396 | 0.0660619868429705 |
| Cep19 | 78.2293441769459 | 1.21630577672581 | 0.387285826034861 | 3.14058944314767 | 0.00168608213190382 | 0.0293875130200164 |
| Npnt | 135.995269614094 | 1.21631076388623 | 0.326430463925089 | 3.72609452335108 | 0.000194469502587806 | 0.00711348969992236 |
| Cd72 | 85.710229394253 | 1.22454251730084 | 0.317526716591651 | 3.85650231402618 | 0.00011502100972928 | 0.00490051198540074 |
| Sord | 274.078717124866 | 1.23560976919356 | 0.325509995589858 | 3.79591959059354 | 0.000147097174160375 | 0.00577993136630165 |
| Zfp827 | 470.398404480664 | 1.23674571621786 | 0.296594932754656 | 4.1698140448033 | 3.04848275591689e-05 | 0.00179360466908972 |
| Wwtr1 | 978.863509066126 | 1.23692055385786 | 0.158653343388496 | 7.79637243968442 | 6.37121632234883e-15 | 7.87199172272433e-12 |
| Rtn4r | 67.6341305952206 | 1.25210564929384 | 0.367722133777634 | 3.40503204534052 | 0.000661562900632728 | 0.0158198992577674 |
| Itga9 | 74.5201177689111 | 1.26887042526475 | 0.427241943789331 | 2.9699107115064 | 0.00297886317378581 | 0.042251222566962 |
| Prkcb | 689.837056518528 | 1.27575036095299 | 0.213637007776033 | 5.97157942920838 | 2.34967614583762e-09 | 1.25182829813465e-06 |
| Tiparp | 2082.559641231 | 1.28087216031974 | 0.240535301900136 | 5.32509012274432 | 1.00902998734561e-07 | 1.96849358934792e-05 |
| Cd244 | 52.327492080071 | 1.29287168125243 | 0.510994667018537 | 2.53010797313376 | 0.0114027430573859 | 0.0987527280359276 |
| Grb10 | 115.249556146529 | 1.29434610697224 | 0.353282059835671 | 3.66377536287662 | 0.000248524766159135 | 0.0084513620785614 |
| Gpr35 | 55.9769055394037 | 1.29945684117043 | 0.383812465812538 | 3.3856556441424 | 0.000710084284196178 | 0.0166937362373393 |
| Nlrp3 | 163.101749958926 | 1.31078235652701 | 0.330877181428374 | 3.96153748308798 | 7.44686885371408e-05 | 0.00353885391680772 |
| Itgam | 68.3870331269408 | 1.3112220589924 | 0.470772837934713 | 2.78525427410968 | 0.00534857620898587 | 0.0620836820917776 |
| Pla2g7 | 8107.07639030953 | 1.31420833735949 | 0.145862664848627 | 9.00990214818404 | 2.06240427589525e-19 | 4.58678710959105e-16 |
| Esd | 8992.38151459984 | 1.34475958248119 | 0.376988577566445 | 3.56710962215872 | 0.000360940513096126 | 0.0107740972421658 |
| Ptprf | 224.153354552924 | 1.35617126817498 | 0.340635921301891 | 3.98129258650049 | 6.85415160219027e-05 | 0.00335762845005973 |
| Tnfsf4 | 59.3825595500505 | 1.35753762150896 | 0.428482634058785 | 3.16824420315412 | 0.00153362640643285 | 0.0273738774310326 |
| Gclm | 889.128239113679 | 1.36360023651397 | 0.450986519198154 | 3.02359422835616 | 0.00249791225730748 | 0.0374350192739342 |
| Them6 | 31.6105974155892 | 1.3710080872448 | 0.409866926059745 | 3.34500785517138 | 0.000822802400246412 | 0.0183357969754311 |
| Blnk | 99.0227101630571 | 1.3732632608399 | 0.371721447422484 | 3.69433421278784 | 0.000220463677406261 | 0.00778271775478609 |
| Metap1d | 20.7945208649402 | 1.37349782139298 | 0.492382892992109 | 2.78949135102261 | 0.00527909064516729 | 0.0615986232678491 |
| Sgk1 | 2160.56794946985 | 1.37673420082583 | 0.125619362587256 | 10.9595700254372 | 5.97827248145577e-28 | 2.21594633312627e-24 |

| | | | | | | |
|---|---|---|---|---|---|---|
| 4930539E08R | 86.4468399526551 | 1.38581833214945 | 0.321119345097681 | 4.31558656713097 | 1.59179655279823e-05 | 0.00106631190765761 |
| Ttll7 | 101.708558163298 | 1.38630300952189 | 0.548242237180064 | 2.52863226418394 | 0.0114507932592147 | 0.0990843277890364 |
| Lpar1 | 140.723376355736 | 1.39687974623773 | 0.435675598763466 | 3.20623819695745 | 0.00134482616317858 | 0.025049358349323 |
| Stap1 | 952.118623820203 | 1.4085724565066 | 0.237379300255715 | 5.93384703295204 | 2.95917549982218e-09 | 1.43069702426185e-06 |
| Hrsp12 | 43.3418845742503 | 1.41068660690343 | 0.337395789842569 | 4.18110317132785 | 2.90098188292737e-05 | 0.00174149941575699 |
| Adk | 56.258108801507 | 1.41871421393667 | 0.453908664100145 | 3.12554997545422 | 0.00177472911800028 | 0.0305022995242089 |
| Hist1h1b | 30.3847696246095 | 1.4187879257261 | 0.507678023378922 | 2.7946609078785 | 0.00519541784911258 | 0.0610134356419227 |
| Ednrb | 45.5131990324226 | 1.44771103552237 | 0.504219297824207 | 2.87119323232865 | 0.00408925471643901 | 0.051920407984837 |
| Nr1d1 | 89.2545718714182 | 1.48219802306682 | 0.447250375347181 | 3.31402298302434 | 0.00091963947781517 | 0.0197969130460683 |
| Gas7 | 124.061547409518 | 1.49880254945276 | 0.49194955590915 | 3.04665901503436 | 0.00231400047380799 | 0.0359381079172414 |
| Rgs9 | 124.833523066249 | 1.50073401628758 | 0.288325900462566 | 5.20499203810663 | 1.94004924026258e-07 | 3.47957218576127e-05 |
| Slc7a11 | 4080.39062618051 | 1.50538698628856 | 0.249929067223564 | 6.02325693050331 | 1.70941754565363e-09 | 1.05604017264824e-06 |
| Pla2g4a | 432.719577804449 | 1.5091337155952 | 0.2554656170596 | 5.90738484875295 | 3.4758111967735e-09 | 1.48609840579005e-06 |
| Ly9 | 75.2637539858989 | 1.51035624061972 | 0.457382901631337 | 3.30217031557754 | 0.000959397965816525 | 0.0201939190601687 |
| Gm6194 | 62.214166738894 | 1.51181134179287 | 0.339680813423947 | 4.45068217587554 | 8.55979445775049e-06 | 0.000679892245501325 |
| Cxcl2 | 2497.93851834448 | 1.54346116583388 | 0.41208090840948 | 3.74552941991712 | 0.000180013748813107 | 0.00665034181661712 |
| Scd1 | 450.961582109931 | 1.55167818666084 | 0.556476671199754 | 2.78839755009936 | 0.00529694982060138 | 0.0617422243239909 |
| Dpy19l3 | 97.76578100606 | 1.56097809371699 | 0.287113385612663 | 5.43680013520115 | 5.42459278391476e-08 | 1.18897866097857e-05 |
| Rnf144b | 36.4255182826396 | 1.56316907233954 | 0.445689537015848 | 3.50730484454688 | 0.000452670250803743 | 0.0123677965330163 |
| Acss1 | 162.083367311562 | 1.5775047846207 | 0.493271019402865 | 3.19804878569669 | 0.00138360871017187 | 0.0255153049039987 |
| Cd300a | 349.683285696734 | 1.59259174235328 | 0.335776898187682 | 4.74300570095533 | 2.10570341580135e-06 | 0.000238932877384806 |
| Ccr1 | 58.2715278145454 | 1.59436404743167 | 0.513028630458136 | 3.10774867673155 | 0.00188518308843021 | 0.0314763302452612 |
| Cystm1 | 154.230213803715 | 1.63468309695939 | 0.343063850211318 | 4.76495292626277 | 1.88897744658842e-06 | 0.000221109781116455 |
| Synj2 | 65.4363521099128 | 1.63510311667071 | 0.29952423289917 | 5.4590010993239 | 4.78820796378712e-08 | 1.10926817827735e-05 |
| 5031414D18R | 40.5277973228704 | 1.66102628527905 | 0.499837028974008 | 3.32313572023377 | 0.000890115957534426 | 0.0194080185250643 |
| Gm4961 | 17.0735734651224 | 1.66602467074465 | 0.601848257160882 | 2.76818060187437 | 0.00563702085622638 | 0.0641593366645213 |
| Olfr726 | 25.1579566008532 | 1.67364526040128 | 0.618060905311573 | 2.70789698234929 | 0.00677110366756932 | 0.0717743839738763 |
| Nid1 | 28.0002767913925 | 1.67678148311329 | 0.459719317508883 | 3.64740270693735 | 0.000264904568924338 | 0.00881957726478636 |
| Thbs1 | 336.071214606053 | 1.68749967470654 | 0.257797509826227 | 6.54583388274011 | 5.91643166270306e-11 | 4.38604800595054e-08 |
| Dfna5 | 73.3822568437465 | 1.6900752646993 | 0.497664820986661 | 3.3960111171784 | 0.000683755823778305 | 0.0162465058983221 |
| Sema4f | 295.502863581976 | 1.69779880992136 | 0.249601941536208 | 6.80202565521739 | 1.03158221515758e-11 | 9.55932852712692e-09 |
| Vcan | 96.7325395603629 | 1.70068163587025 | 0.39207399600709 | 4.33765476208601 | 1.44011126727522e-05 | 0.000994660701372698 |
| Akr1b8 | 69.3627930609996 | 1.72342218617038 | 0.386198441930935 | 4.46253013749491 | 8.09975321653517e-06 | 0.000655579437835637 |
| Ppbp | 57.8403036756564 | 1.77446382698191 | 0.568340100731329 | 3.12218656522487 | 0.00179513140450182 | 0.0307105557200926 |
| Mfap3l | 36.3242199150145 | 1.79020587744964 | 0.542076473721977 | 3.30249690630886 | 0.000958281450117389 | 0.0201939190601687 |
| Fbxo36 | 29.4974516307941 | 1.80776243014071 | 0.506555192555058 | 3.56873733940497 | 0.000358705817970275 | 0.0107740972421658 |
| Procr | 571.734454116672 | 1.86227414851863 | 0.586432737946307 | 3.17559717938042 | 0.0014952842852063 | 0.0270366849617789 |
| Il10 | 112.826625749554 | 1.87312824432473 | 0.397422416279351 | 4.71319222972087 | 2.43865941932017e-06 | 0.000268493987552874 |
| Itgb8 | 2140.81744933748 | 1.91053038182363 | 0.119657704881088 | 15.9666306797565 | 2.18261893523376e-57 | 2.42707225597994e-53 |
| Slc7a2 | 2659.42293571377 | 1.92926363356412 | 0.231035228911142 | 8.35051711661743 | 6.79647424261475e-17 | 1.07966847968394e-13 |
| Tfec | 39.8223146104817 | 1.97973212664935 | 0.438304404333222 | 4.51679724656442 | 6.27819659908941e-06 | 0.000524913881066724 |
| Slc11a1 | 195.649907089907 | 1.99756174296418 | 0.753263060116182 | 2.65187800747335 | 0.00800454569781921 | 0.0791220541158187 |
| Nudt12 | 26.2925251569177 | 2.01220902390982 | 0.549961681750485 | 3.65881676975224 | 0.000253382413569345 | 0.00853821951179128 |
| Clec4d | 254.248511556916 | 2.02627504605073 | 0.415721100415693 | 4.87412124143949 | 1.09293855655537e-06 | 0.000144684247010663 |
| Mcoln2 | 55.4022118014258 | 2.03545050655257 | 0.565853316988929 | 3.59713453193806 | 0.000321742096081556 | 0.010164125308031 |
| Emb | 912.223701190384 | 2.06570800244585 | 0.324952220719845 | 6.356959179629 | 2.05786649111812e-10 | 1.43021721132709e-07 |
| Mmp19 | 151.406846819163 | 2.10946534058219 | 0.383489571920548 | 5.50071108848623 | 3.78262590204777e-08 | 9.14408696321112e-06 |
| AV051173 | 48.4044671920518 | 2.11547719081816 | 0.47706482231796 | 4.43436005308354 | 9.23461828511041e-06 | 0.000713117745350193 |
| Cxcl3 | 2482.14308886966 | 2.14769916554288 | 0.626936270267231 | 3.42570571746858 | 0.000613204374351371 | 0.015022150300123 |
| F5 | 42.4913559176138 | 2.17380712682047 | 0.632479490166246 | 3.43696066136325 | 0.000588281038370894 | 0.0146346423863184 |
| Crygn | 27.5740015241406 | 2.1773146321303 | 0.466788420286735 | 4.66445725194477 | 3.09432646117167e-06 | 0.000315678075671826 |
| Avil | 190.083420054468 | 2.18297390309523 | 0.726307643865334 | 3.00557748707927 | 0.00265077046726234 | 0.0390418113853738 |
| Abcb4 | 33.6922571155281 | 2.30129375085668 | 0.645513453681514 | 3.56505931477007 | 0.000363773910501172 | 0.0107913177489103 |
| Clec4e | 712.850896865535 | 2.30477563613256 | 0.796271562326658 | 2.89445930908061 | 0.00379812133474808 | 0.050002598923535 |
| Hopx | 151.72683361384 | 2.31307647951609 | 0.448949424039163 | 5.15219834498399 | 2.57450529069115e-07 | 4.3376513382554e-05 |
| Lekr1 | 20.9873568533116 | 2.33151881168312 | 0.647930992056051 | 3.59840606525799 | 0.000320173450310432 | 0.0101433868018576 |
| Areg | 50.8665012546411 | 2.34606281766821 | 0.459523590257636 | 5.1054241118565 | 3.30053425655913e-07 | 5.16928745534331e-05 |
| Sel1l3 | 61.0340398745959 | 2.38767100575583 | 0.509242328668899 | 4.68867348870415 | 2.74981732574469e-06 | 0.000285775408058701 |
| Rnasel | 27.426065511315 | 2.3990423816717 | 0.859160543576081 | 2.79230977214826 | 0.00523332287914499 | 0.0612574214906234 |
| Nqo1 | 210.299157250667 | 2.46265900735939 | 0.30972727160261 | 7.95105640719641 | 1.84927715156802e-15 | 2.57049524067955e-12 |
| Ptgs2 | 2878.33477505545 | 2.5733293100685 | 0.245122645535319 | 10.4981296381191 | 8.81085636372952e-26 | 2.44941806911681e-22 |
| Pdgfc | 49.1988376283799 | 2.5976912583969 | 0.594924556576796 | 4.3664213044828 | 1.26298762153101e-05 | 0.000883297003234269 |
| Tmtc1 | 54.6943478470015 | 2.60946037073992 | 0.483036035197845 | 5.40220642062682 | 6.58261367753892e-08 | 1.40766661719679e-05 |
| Peg10 | 63.3362057923028 | 2.69512140258795 | 0.74304126759784 | 3.62714901596373 | 0.000286567924448796 | 0.00932154447170718 |
| Xdh | 63.2206253616052 | 2.70290847963506 | 0.502301051192031 | 5.38105280333513 | 7.40514672671239e-08 | 1.52491169631559e-05 |
| Hpgd | 16.554556533266 | 2.72868791155978 | 0.898180682449282 | 3.03801669850973 | 0.0023814075812862 6 | 0.0364367623618618 |
| Syt7 | 288.985839278059 | 3.36355260080477 | 0.281167730140285 | 11.9627974345654 | 5.56548549238771e-33 | 3.094409933376757e-29 |
| Igfbp2 | 26.2503742465217 | 3.43159925535528 | 0.581507193242257 | 5.90121548836227 | 3.60833246010173e-09 | 1.48609840579005e-06 |
| Uty | 229.659433377459 | 7.09131165224007 | 2.64211517961707 | 2.68395250401909 | 0.00727574416585448 | 0.0746367851700201 |

| | | | | | | |
|---|---|---|---|---|---|---|
| Ddx3y | 700.100600627655 | 7.71186993999144 | 2.95511210662683 | 2.60967085570039 | 0.00906293760381844 | 0.0862841319815591 |
