## Supplementary Table 3 for "Epidermal maintenance of Langerhans cells relies on autophagy-regulated lipid metabolism"

[https://www.genome.jp/pathway/hsa04216](https://www.genome.jp/pathway/hsa04216) ***39% <=> 16/41 genes with p<0.05*** ***24% <=> 10/41 genes with p<0.05 and FDR<0.1***

| Genes in pathway ***(italics: FDR>0.1)*** | **genes with p<0.05** | **baseMean** | **log2FoldChange** | **lfcSE** | **stat** | **p** | **FDR** |
|---|---|---|---|---|---|---|---|
| ACSL1 | | | | | | | |
| ACSL3 | Acsl3 | 480.400019648201 | 1.01272696548078 | 0.338122545172163 | 2.99514770588612 | 0.00274311972146471 | 0.0400308284812173 |
| ACSL4 | Acsl4 | 2791.99445343052 | 0.367732593896325 | 0.121121027005226 | 3.03607559305504 | 0.00239679266122118 | 0.0365372098580231 |
| ACSL5 | | | | | | | |
| ACSL6 | | | | | | | |
| ALOX15 | | | | | | | |
| ATG5 | Atg5 | 571.084089225324 | 0.433204501918915 | 0.171410880645814 | 2.52728706769814 | 0.0114947503691776 | 0.0993175012472841 |
| ATG7 | | | | | | | |
| CP (ceruloplasmin) | | | | | | | |
| CYBB | | | | | | | |
| ***FTH1*** | ***Fth1*** | ***36706.3133164478*** | ***1.40400183864457*** | ***0.611413193693371*** | ***2.29632244303299*** | ***0.0216574517962314*** | ***0.142373636481794*** |
| FTL | Ftl1 | 62767.6287757189 | 0.366424633014218 | 0.0976059160744037 | 3.75412318998059 | 0.000173949304060843 | 0.00650754181686748 |
| FTMT | | | | | | | |
| ***GCLC*** | ***Gclc*** | ***524.787109399089*** | ***0.490107803613869*** | ***0.217794381850878*** | ***2.25032344475002*** | ***0.0244284207322337*** | ***0.153570632800047*** |
| GCLM | Gclm | 889.128239113679 | 1.36360023651397 | 0.450986519198154 | 3.02359422835616 | 0.00249791225730748 | 0.0374350192739342 |
| GPX4 | | | | | | | |
| GSS | Gss | 198.54249109314 | 1.15371698730864 | 0.285285078255757 | 4.04408458501406 | 0.0000525279542067106 | 0.00275523986216331 |
| ***HMOX1*** | ***Hmox1*** | ***3031.83370684409*** | ***1.8712486673362*** | ***0.826503544120706*** | ***2.26405401482818*** | ***0.0235707928074841*** | ***0.150720231050148*** |
| LPCAT3 | Lpcat3 | 189.13876534598 | 0.612442102942747 | 0.189190980188215 | 3.23716332741372 | 0.00120724295713734 | 0.0231059237235236 |
| MAP1LC3A | | | | | | | |
| MAP1LC3B | Map1lc3b | 1418.37934067585 | 0.285166355779976 | 0.0877621128148816 | 3.24931051262955 | 0.00115685122785492 | 0.0224898350589978 |
| MAP1LC3B2 | | | | | | | |
| MAP1LC3C | | | | | | | |
| NCOA4 | | | | | | | |
| PCBP1 | | | | | | | |
| PCBP2 | | | | | | | |
| PRNP | | | | | | | |
| SAT1 | Sat1 | 3142.91034639789 | 0.576184831781132 | 0.139308354737322 | 4.13603931270009 | 0.000035335168177557 | 0.0020253972687342 |
| ***SAT2*** | ***Sat2*** | ***14.9006849238277*** | ***1.819569761809*** | ***0.758668803460777*** | ***2.39837166561848*** | ***0.0164681460783392*** | ***NA*** |
| SLC11A2 | | | | | | | |
| SLC39A14 | | | | | | | |
| SLC39A8 | | | | | | | |
| ***SLC3A2*** | ***Slc3a2*** | ***4391.73055573213*** | ***0.505206863923555*** | ***0.212969407078203*** | ***2.37220392757182*** | ***0.0176823283009167*** | ***0.127887279760233*** |
| SLC40A1 | | | | | | | |
| SLC7A11 | Slc7a11 | 4080.39062618051 | 1.50538698628856 | 0.249929067223564 | 6.02325693050331 | 1.70941754565363E-09 | 1.05604017264824E-06 |
| STEAP3 | | | | | | | |
| TF | | | | | | | |
| TFRC | Tfrc | 161.36882196451 | 0.927325198566473 | 0.274900051390578 | 3.37331766173055 | 0.00074268217244985 | 0.0171697001198385 |
| TP53 | | | | | | | |
| VDAC2 | | | | | | | |
| VDAC3 | | | | | | | |
| ***41*** | ***16*** | | | | | | ***6*** |

*(13 visible on KEGG pathway map)*

https://www.genome.jp/pathway/hsa04210 *19% <=> 26/137 genes with p<0.05* *9% <=> 13/137 genes with p<0.05 and FDR<0.1*

| Genes in pathway ***(italics: FDR>0.1)*** | genes with p<0.05 | baseMean | log2FoldChange | lfcSE | stat | p | FDR |
|---|---|---|---|---|---|---|---|
| ACTB | | | | | | | |
| ACTG1 | | | | | | | |
| AIFM1 | | | | | | | |
| AKT1 | | | | | | | |
| AKT2 | | | | | | | |
| AKT3 | | | | | | | |
| APAF1 | Apaf1 | 3967.41815167333 | 0.316093980709645 | 0.0803774131836253 | 3.93262196666515 | 0.0000840243226739688 | 0.00382930519727268 |
| ATF4 | | | | | | | |
| ATM | | | | | | | |
| BAD | | | | | | | |
| ***BAK1*** | ***Bak1*** | ***753.653030341207*** | ***-0.306908173789676*** | ***0.155757172260808*** | ***-1.9704272319208*** | ***0.0487894271457412*** | ***0.234054542649112*** |
| BAX | | | | | | | |
| BBC3 | | | | | | | |
| BCL2 | | | | | | | |
| BCL2A1 | | | | | | | |
| BCL2L1 | Bcl2l1 | 1434.03385185298 | -0.846011853670069 | 0.234887795710745 | -3.60177016055742 | 0.000316057740029919 | 0.0100416059118077 |
| ***BCL2L11 (Bim)*** | ***Bcl2l11*** | ***1593.27198141821*** | ***0.386934924530282*** | ***0.169502422699883*** | ***2.28276928652153*** | ***0.0224439620286047*** | ***0.145804558443916*** |
| BID | | | | | | | |
| BIRC2 | | | | | | | |
| BIRC3 | | | | | | | |
| ***BIRC5 (IAP/XIAP-related)*** | ***Birc5*** | ***20.7756125997574*** | ***1.58238004080547*** | ***0.647813301225646*** | ***2.44264827198153*** | ***0.0145799391282185*** | ***0.113774682881256*** |
| CAPN1 | | | | | | | |
| CAPN2 | | | | | | | |
| CASP10 | | | | | | | |
| CASP12 | | | | | | | |
| CASP2 | | | | | | | |
| CASP3 | | | | | | | |
| CASP6 | | | | | | | |
| CASP7 | Casp7 | 136.509460268506 | -1.23633014026956 | 0.271084055567175 | -4.56068925810799 | 5.09859850078706E-06 | 0.000460946466087415 |
| CASP8 | | | | | | | |
| CASP9 | | | | | | | |
| CFLAR | | | | | | | |
| CHUK | | | | | | | |
| CSF2RB (IL3R) | Csf2rb | 51584.2004096577 | 0.336110477181865 | 0.108380339383464 | 3.10121262854383 | 0.0019272984252942 | 0.0319397294922079 |
| CTSB | | | | | | | |
| CTSC | Ctsc | 1106.86569734841 | -0.580131249116454 | 0.149878857990036 | -3.87066766384769 | 0.000108537676856939 | 0.00469626057061931 |
| CTSD | | | | | | | |
| CTSF | | | | | | | |

| | | | | | | | |
|---|---|---|---|---|---|---|---|
| CTSH | Ctsh | 1461.61771308929 | -0.540180508670733 | 0.154496351139211 | -3.49639654715208 | 0.000471587314335762 | 0.0126974598920428 |
| CTSK | | | | | | | |
| CTSL | | | | | | | |
| CTSO | | | | | | | |
| CTSS | | | | | | | |
| CTSV | | | | | | | |
| CTSW | | | | | | | |
| CTSZ | | | | | | | |
| CYCS | | | | | | | |
| DAB2IP | | | | | | | |
| DAXX | | | | | | | |
| DDIT3 | | | | | | | |
| DFFA | | | | | | | |
| DFFB | | | | | | | |
| DIABLO | | | | | | | |
| EIF2AK3 | | | | | | | |
| EIF2S1 | | | | | | | |
| ENDOG | | | | | | | |
| ERN1 (IRE1alpha) | Ern1 | 536.458096269271 | 0.608523032452516 | 0.179351972051314 | 3.39289847495187 | 0.00069157264867628 | 0.0163623145814473 |
| FADD | | | | | | | |
| FAS | | | | | | | |
| FASLG | | | | | | | |
| FOS | Fos | 873.905953197261 | 0.732117103199913 | 0.225117444159312 | 3.25215625085813 | 0.00114533040695148 | 0.0223832585681906 |
| GADD45A | | | | | | | |
| GADD45B | | | | | | | |
| ***GADD45G*** | ***Gadd45g*** | ***402.92603104958*** | ***0.433344440183135*** | ***0.207100248737512*** | ***2.09243804787688*** | ***0.0363993482239758*** | ***0.197540630673797*** |
| genes | | | | | | | |
| GZMB | | | | | | | |
| HRAS | | | | | | | |
| HRK | | | | | | | |
| HTRA2 | Htra2 | 256.791031971511 | -0.678960395379265 | 0.230583424280717 | -2.9445325373982 | 0.0032344293557207 | 0.0446238888779333 |
| IKBKB | | | | | | | |
| IKBKG | Ikbkg | 1363.02689537012 | 0.698287925633453 | 0.182313995494265 | 3.83013889712827 | 0.000128070947597378 | 0.00528952718059445 |
| IL3 | | | | | | | |
| IL3RA | | | | | | | |
| ITPR1 | | | | | | | |
| ITPR2 | | | | | | | |
| ITPR3 | | | | | | | |
| JUN | | | | | | | |
| KRAS | Kras | 1456.50970363713 | 0.25687269453924 | 0.0930178163268076 | 2.76154294610343 | 0.00575289500339731 | 0.0651447988164746 |
| LMNA | | | | | | | |

| | | | | | | | |
|---|---|---|---|---|---|---|---|
| LMNB1 | | | | | | | |
| LMNB2 | | | | | | | |
| MAP2K1 | | | | | | | |
| ***MAP2K2 (MEK2)*** | ***Map2k2*** | ***1801.31557956592*** | ***0.22891928268153*** | ***0.0949344776133296*** | ***2.41133978336009*** | ***0.0158940333433823*** | ***0.119824847985363*** |
| MAP3K14 | | | | | | | |
| MAP3K5 | | | | | | | |
| MAPK1 (ERK2) | Mapk1 | 2537.94582630728 | 0.312338941757773 | 0.0894249052937382 | 3.49275116067295 | 0.000478071779320947 | 0.0127485807818919 |
| ***MAPK10*** | ***Mapk10*** | ***7.85099094605177*** | ***1.88205559568669*** | ***0.792105791106304*** | ***2.37601544745443*** | ***0.0175007301366277*** | ***NA*** |
| MAPK3 | | | | | | | |
| MAPK8 | | | | | | | |
| MAPK9 | | | | | | | |
| MCL1 | | | | | | | |
| ***NFKB1*** | ***Nfkb1*** | ***21172.4988184562*** | ***-0.401518170311485*** | ***0.172306046700454*** | ***-2.33026163620077*** | ***0.0197923270820356*** | ***0.135550102038925*** |
| ***NFKBIA*** | ***Nfkbia*** | ***9255.91835054725*** | ***-0.265668097595959*** | ***0.133892963487207*** | ***-1.98418266857871*** | ***0.0472354799170597*** | ***0.229881793645285*** |
| NGF | | | | | | | |
| NRAS | | | | | | | |
| NTRK1 | | | | | | | |
| PARP1 | | | | | | | |
| PARP2 | | | | | | | |
| ***PARP3*** | ***Parp3*** | ***1327.87191546372*** | ***-0.30383904808312*** | ***0.137266210382332*** | ***-2.21350212289556*** | ***0.0268630470404066*** | ***0.163501413841993*** |
| PARP4 | | | | | | | |
| PDPK1 | | | | | | | |
| PIDD1 | | | | | | | |
| PIK3CA | | | | | | | |
| PIK3CB | | | | | | | |
| ***PIK3CD*** | ***Pik3cd*** | ***172.166939921816*** | ***0.513715621808792*** | ***0.240246320749821*** | ***2.13828715547218*** | ***0.0324934443971402*** | ***0.183414772434619*** |
| PIK3R1 | | | | | | | |
| PIK3R2 | | | | | | | |
| PIK3R3 | | | | | | | |
| ***PMAIP1 (Noxa)*** | ***Pmaip1*** | ***3522.57039607843*** | ***0.53203288077495*** | ***0.229905184803062*** | ***2.31414041936763*** | ***0.0206600208980078*** | ***0.138003180351218*** |
| PRF1 | | | | | | | |
| PTPN13 | | | | | | | |
| ***RAF1*** | ***Raf1*** | ***837.690878801616*** | ***0.284428300485756*** | ***0.129513712069702*** | ***2.19612499665426*** | ***0.0280829978867221*** | ***0.167605666492612*** |
| RELA | | | | | | | |
| RIPK1 | | | | | | | |
| SEPTIN4 | | | | | | | |
| SPTA1 | | | | | | | |
| SPTAN1 | | | | | | | |
| TNF | | | | | | | |
| TNFRSF10A | | | | | | | |
| TNFRSF10B | | | | | | | |

| | | | | | | | |
|---|---|---|---|---|---|---|---|
| ***TNFRSF1A*** | ***Tnfrsf1a*** | ***613.17232495532*** | ***0.662382176748632*** | ***0.264461336764383*** | ***2.50464655761296*** | ***0.0122573811753929*** | ***0.102791914532707*** |
| TNFSF10 | | | | | | | |
| TP53 | | | | | | | |
| TP53AIP1 | | | | | | | |
| TRADD | | | | | | | |
| TRAF1 | | | | | | | |
| TRAF2 | | | | | | | |
| TUBA1A | | | | | | | |
| TUBA1B | | | | | | | |
| TUBA1C | | | | | | | |
| TUBA3C | | | | | | | |
| TUBA3D | | | | | | | |
| TUBA3E | | | | | | | |
| TUBA4A | Tuba4a | 558.282137394695 | 0.761921789076213 | 0.297010896357224 | 2.56529911333565 | 0.0103086895410443 | 0.0928199414545855 |
| TUBA8 | | | | | | | |
| TUBAL3 | | | | | | | |
| XIAP | | | | | | | |
| ***137*** | ***26*** | | | | | | ***13*** |
| | *(16 visible on KEGG pathway map)* | | | | | | |
