## Supplementary figures and images for "Epidermal maintenance of Langerhans cells relies on autophagy-regulated lipid metabolism"

### Graphical abstract

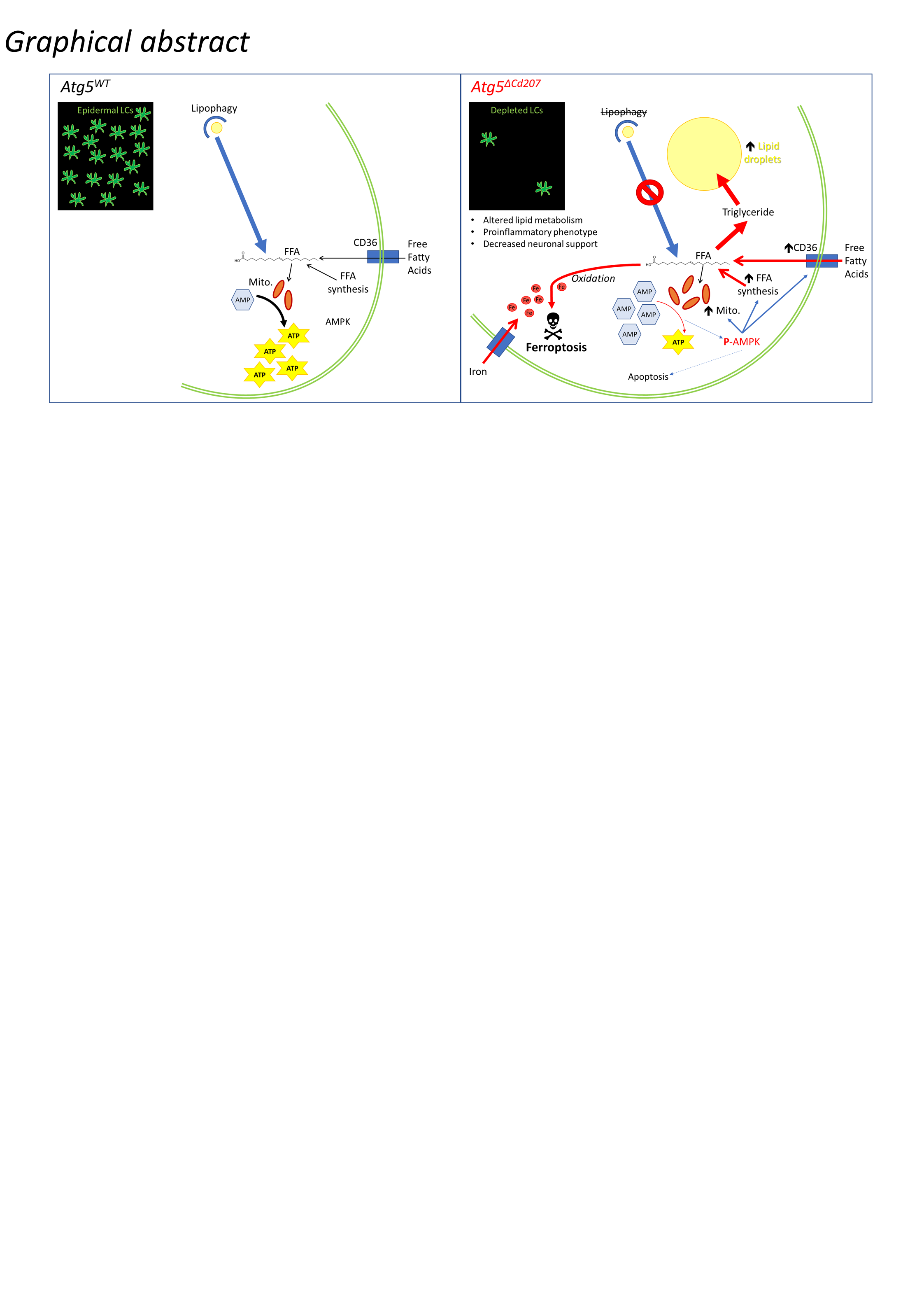

### Supplementary Figure S1

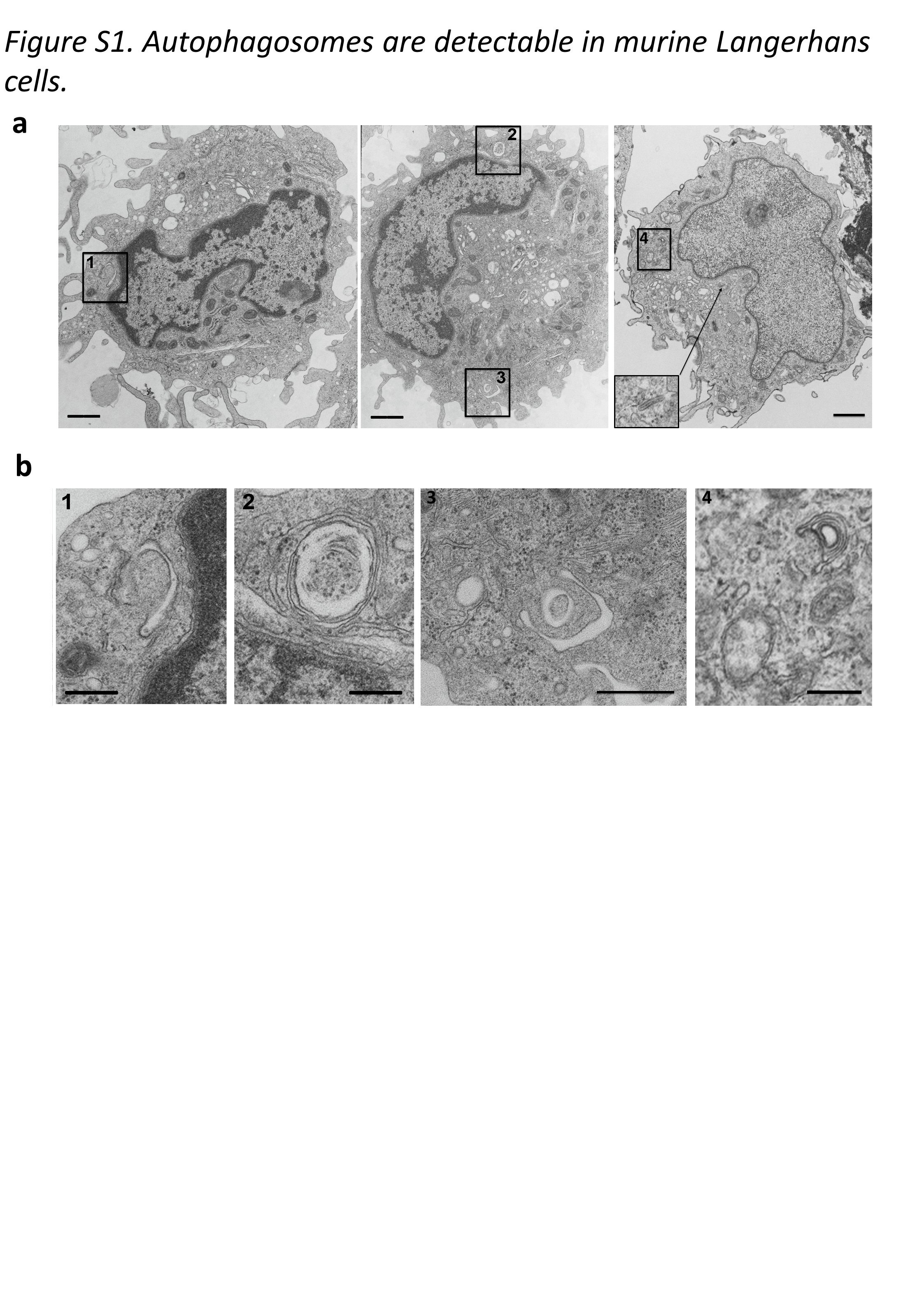

### Supplementary Figure S2

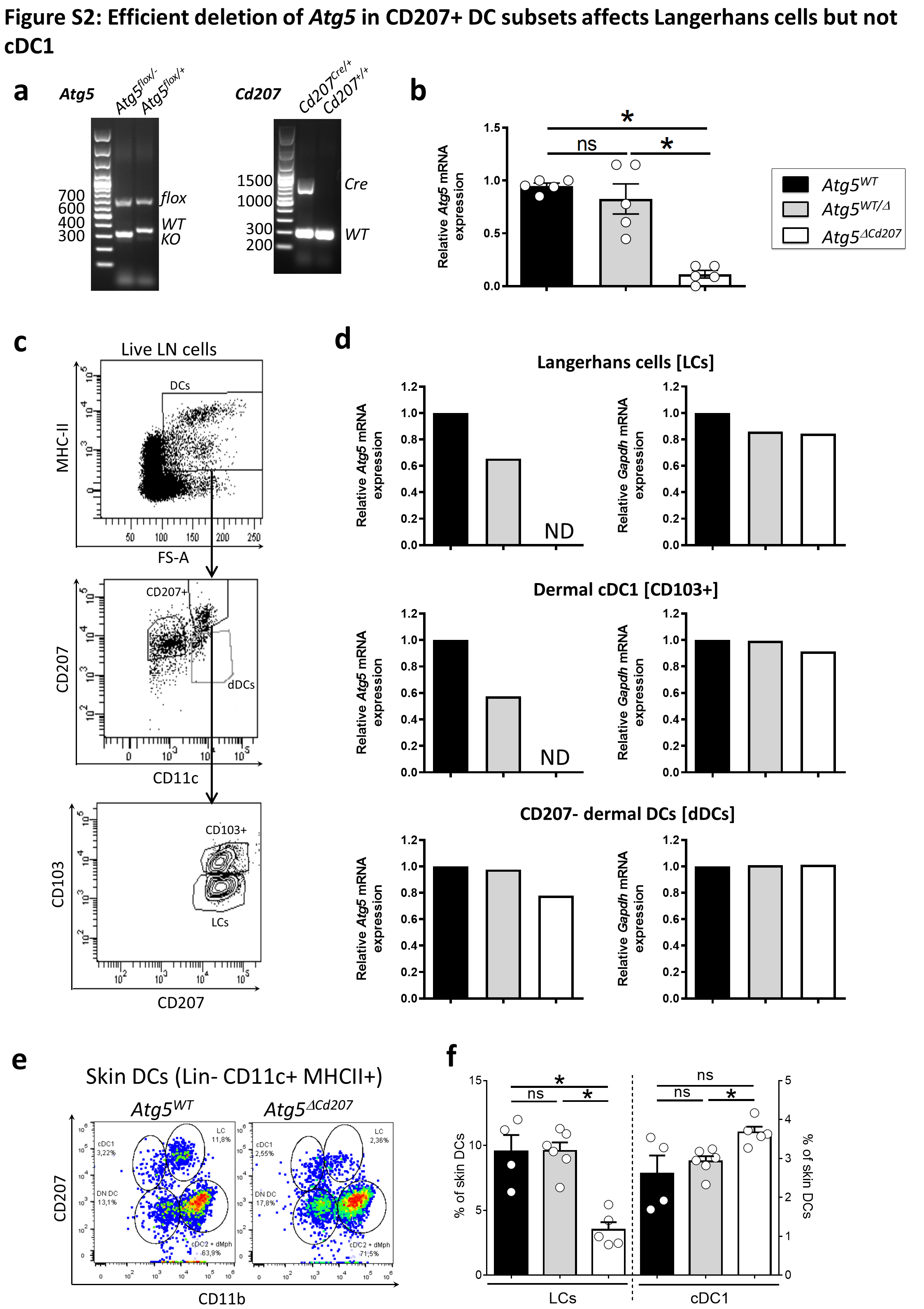

### Supplementary Figure S3

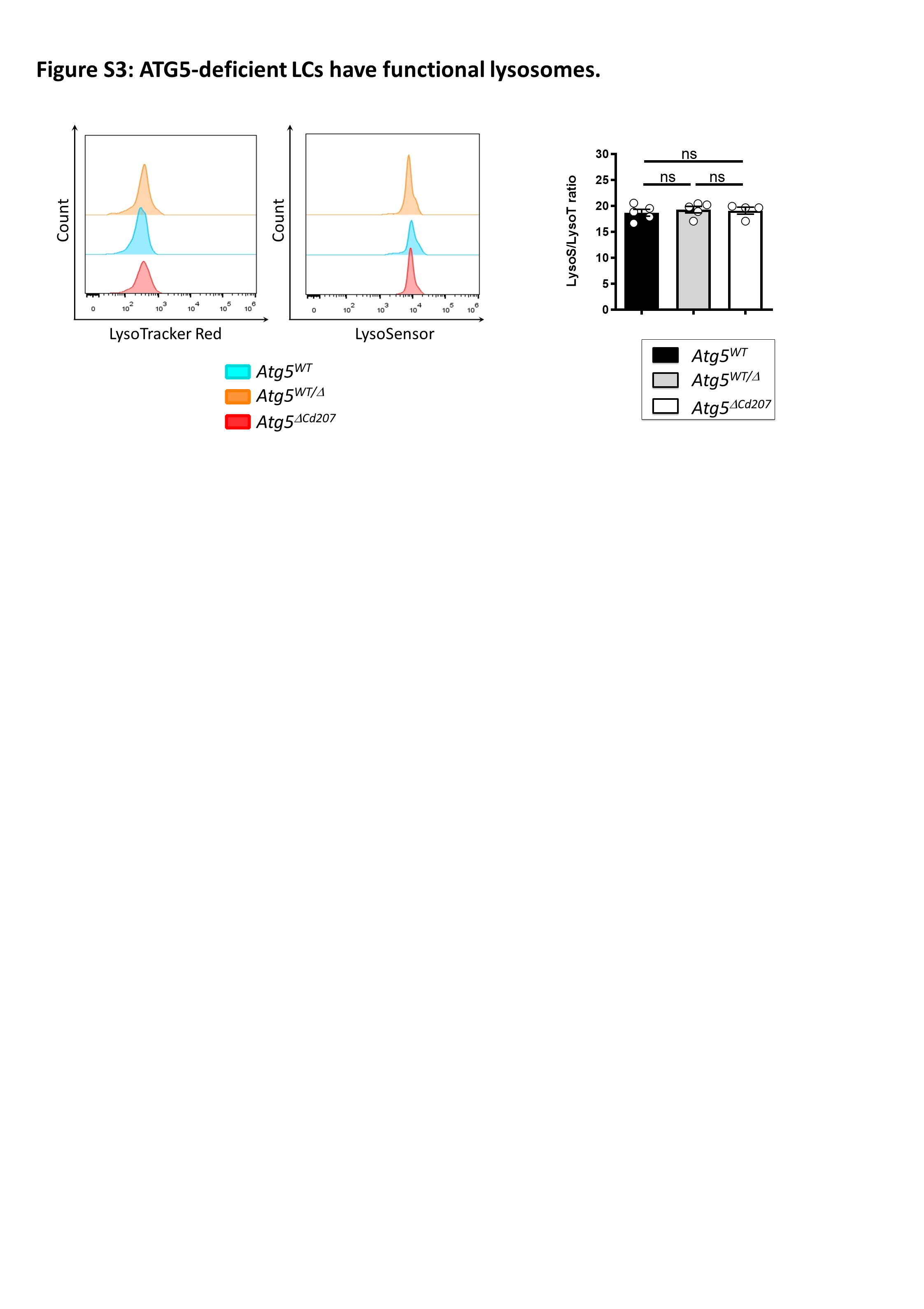

### Supplementary Figure S4

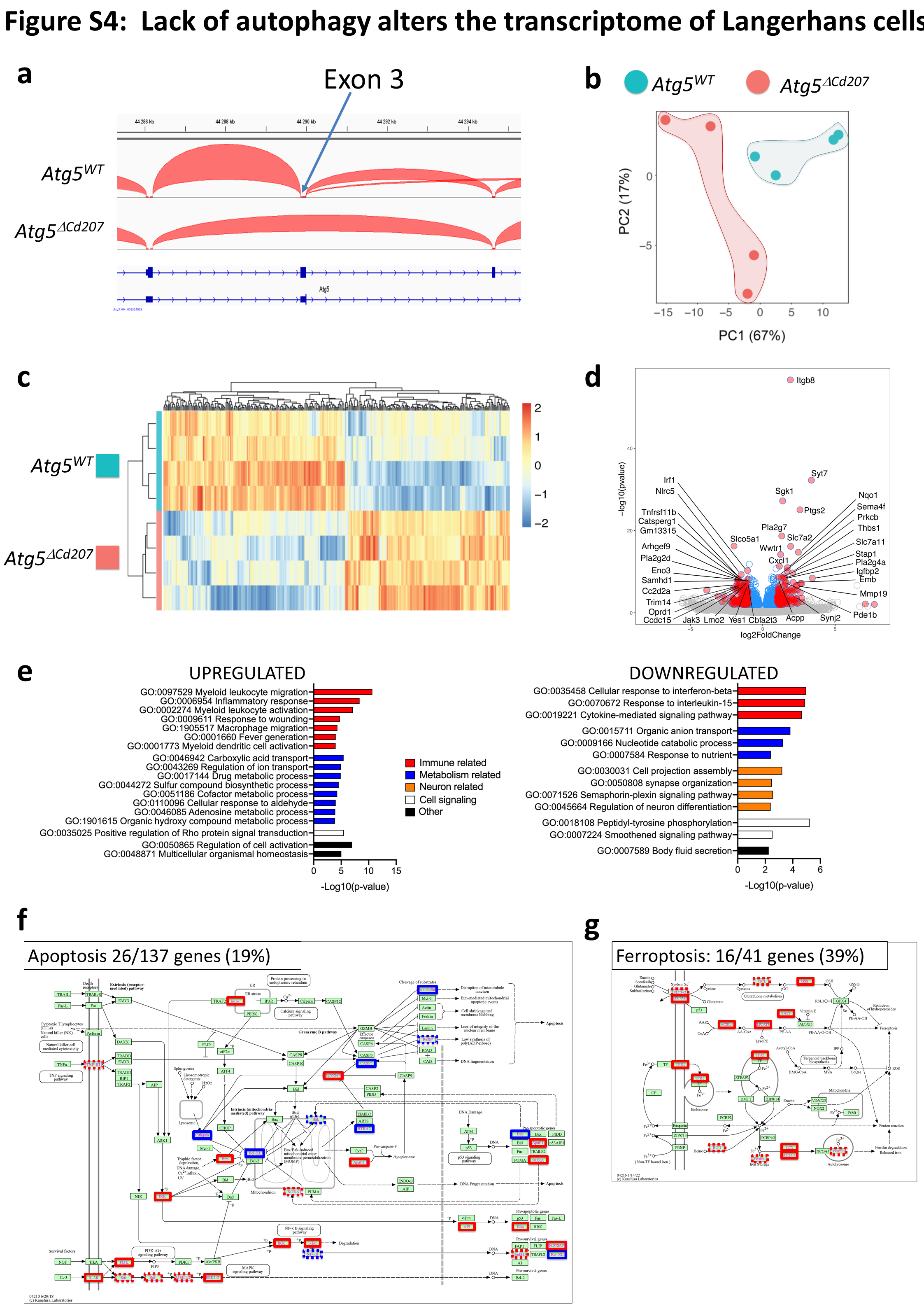

### Supplementary Figure S5

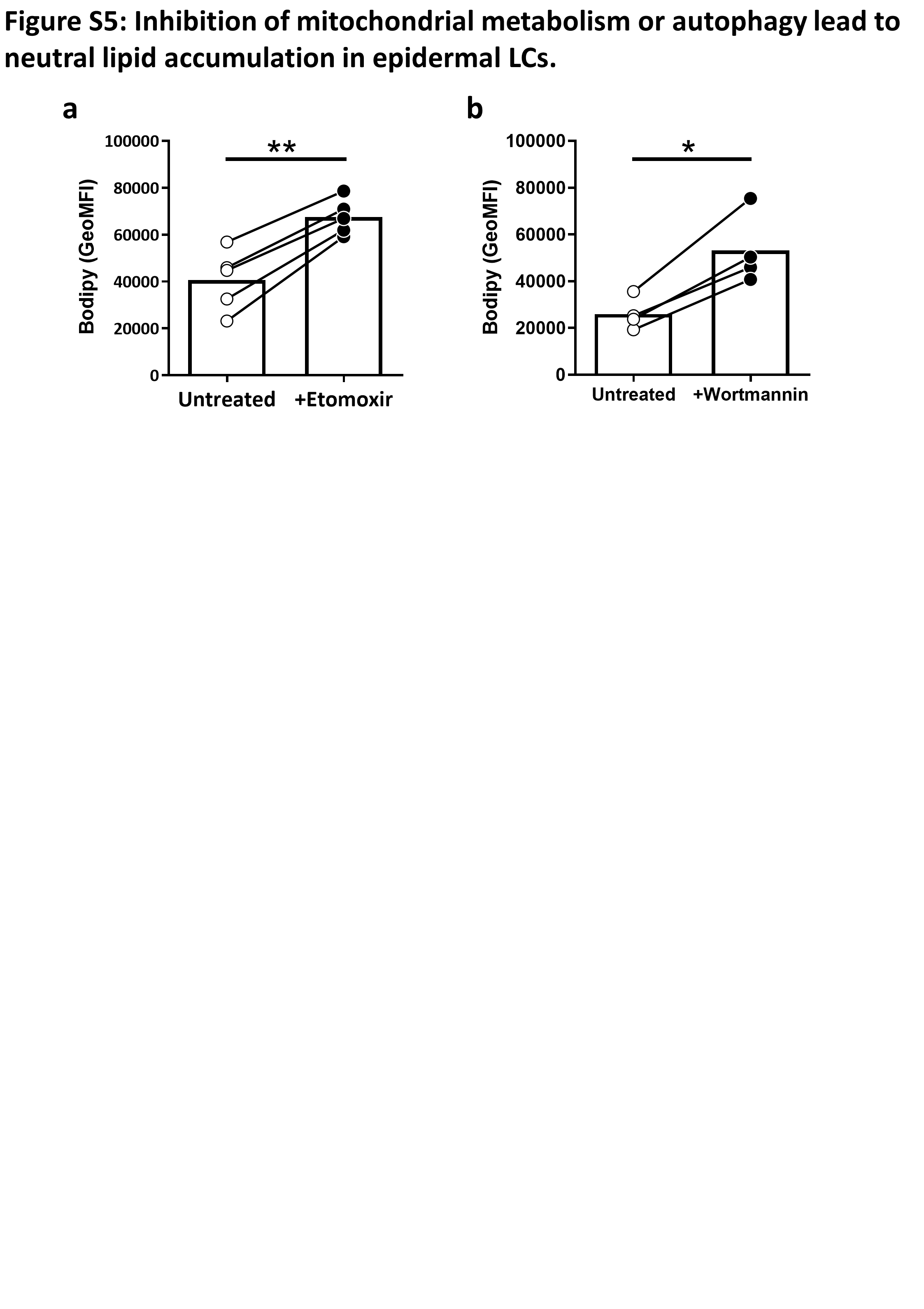

### Supplementary Figure S6

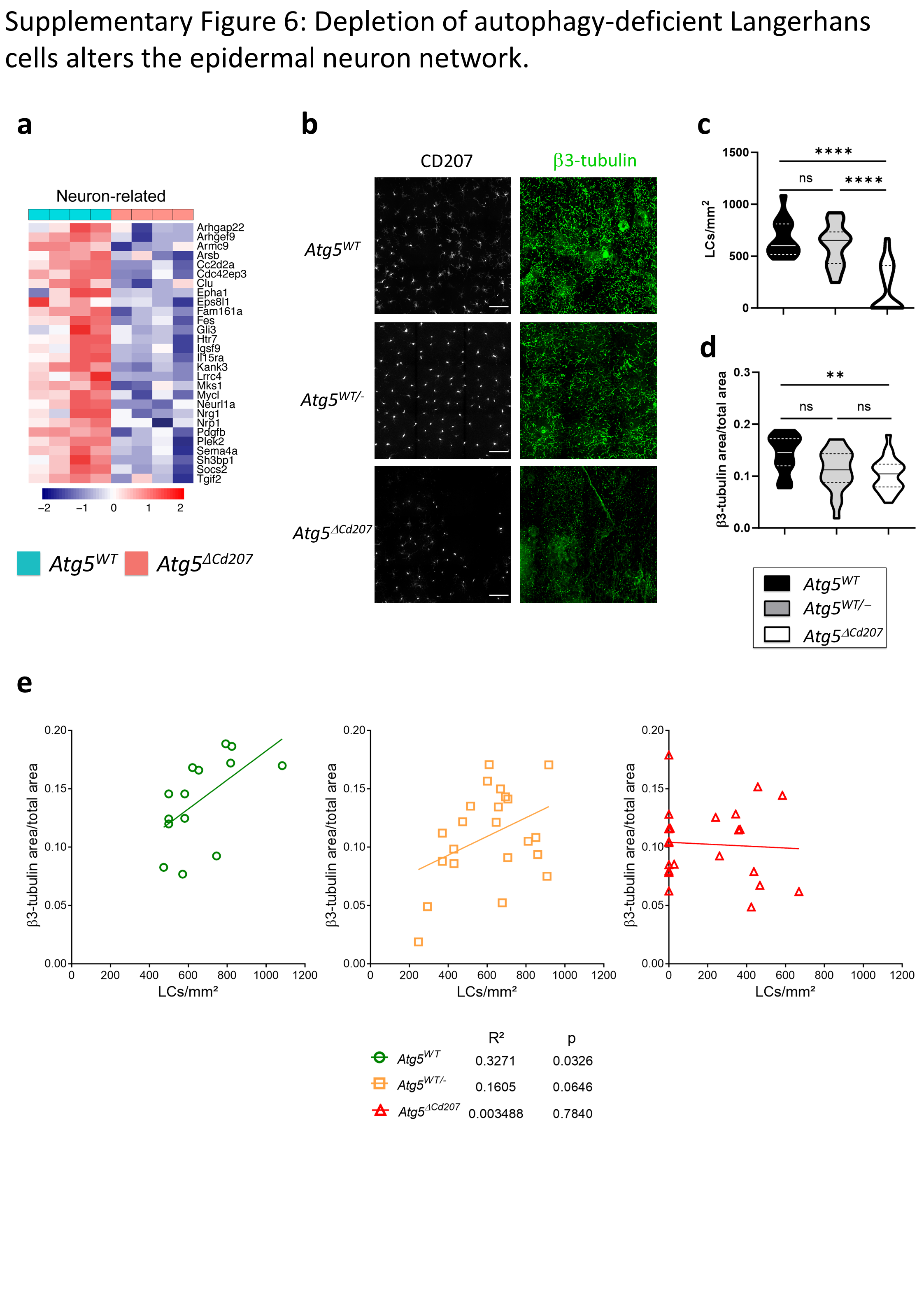
